# Global untreated wastewater hosts a vast reservoir of previously uncharacterized microbial lineages

**DOI:** 10.64898/2026.07.28.741345

**Authors:** Nelson Ruth, Daryl Domman, Brett Youtsey, Po-E Li, Andrew Hatch, Patrick Chain, Migun Shakya

**Author notes:** Corresponding Author: Migun Shakya, Bioscience Division, Los Alamos National Laboratory Los Alamos, New Mexico, USA, LA-UR-25-31023.

## Abstract

Untreated wastewater are reservoirs of microbial communities originating from different sources within an urbanized human-populated area and is now regularly used for tracking and assessing levels of certain human pathogens like SARS-CoV-2, *Salmonella,* and Poliomyelitis. Wastewater microbial communities can exhibit high diversity, comprised of mostly non-pathogens alongside a smaller population of pathogens. Much of the research on wastewater has focused on either pathogens or microbiomes in the treatment plants, but not on the microbiome of untreated wastewater, which is perhaps more reflective of community health. Moreover, as wastewater usage expands to more known and unknown pathogen detection, a deeper understanding of both wastewater’s overall genetic diversity and diversity over time is pivotal. Towards that goal, we characterized the observed microbial diversity of global untreated wastewater by analyzing 1,344 publicly available metagenomes, representing 6 continents, and created a global wastewater database hosted at https://doi.org/10.5281/zenodo.17794790. We elucidated the full spectrum of prokaryotic diversity by recovering and characterizing high quality bins, contigs, and reads from all the data. We provide the first comprehensive, publicly available database describing global wastewater by uncovering previously uncharacterized microbes, identifying geographically specific microbial populations, and creating a high-quality dataset to advance future microbiome, epidemiological, and biosurveillance research.

## Introduction

Untreated wastewater is an ecologically dense and complex environment, whose microbiome remains understudied despite its demonstrated usefulness to public health and biosurveillance, especially during the SARS-CoV-2 pandemic^1–3^. Historically, wastewater microbiome research has relied on cultured or focused analyses to explore taxonomic diversity, track pathogens, and characterize AMR genes^4–6^. While more recent sequencing efforts have significantly expanded our knowledge of the wastewater microbiome, these studies remain constrained by a limited geographical sampling breadth^7^. Metagenomic sequencing has allowed researchers to explore environments without any *a priori* knowledge, leading to discoveries of novel genes and organisms^8^. It also allows for the creation of metagenome-assembled genomes (MAGs), linking genomes to functional^9^ content and improving detection, analysis, and interpretation of potential pathogens, novel AMR genes, and taxonomic diversity^6^.

Several studies have surveyed global wastewater in recent years using read-based metagenomic analyses, but MAG-based investigations remain comparatively limited^10–14^. Previous studies cataloging and analyzing prokaryotic and viral MAGs from wastewater have been relatively small in scale, focusing on specific taxa in a WWTP or wastewater in a single city^15–18^. Here, we present a publicly available database [https://doi.org/10.5281/zenodo.17794790] of wastewater MAGs constructed from untreated global wastewater samples. We processed 1,344 publicly available metagenomes covering 79 countries and 6 continents and created a database containing 9,768 MAGs, reduced to 4,545 prokaryotic species-level genome bins (SGBs), many of which represent novel species, providing a highly specific database of global untreated wastewater taxa. We then showcase that this dataset enables deeper interrogation of pathogenic content in wastewater by characterizing pathogen associated genes and resolving relatedness to closely related strains.

## Results

### Recovery of thousands of prokaryotic genomes from wastewater metagenomes

To characterize the global wastewater microbiome, we processed 1,344 short-read wastewater metagenomic samples from the Sequence Read Archive (SRA) as of May 17^th^, 2022, representing samples from 79 countries across six continents (**Fig. 1A and Supplementary Table 1**). Among these samples are 106 metagenomes that underwent a viral concentration step prior to sequencing, but we included these samples as they likely still contained prokaryotic DNA. All of our samples were from untreated wastewater, most of which were collected from WWTPs or sewers representing larger urban areas (83%, n=1,117) or designated areas like hospitals (11%, n=152) and agricultural sites (4%, n=53) (**Supplementary Table 2)**. The largest contributing continent was Europe (44%, n=588) followed by Africa (14%, n=194), and Asia (14%, n=188). Notably, 55% (n= 324) of the European samples were sequencing runs from a single longitudinal study by a group from the Technical University of Denmark in Copenhagen, Denmark^19^.

**Figure 1.**
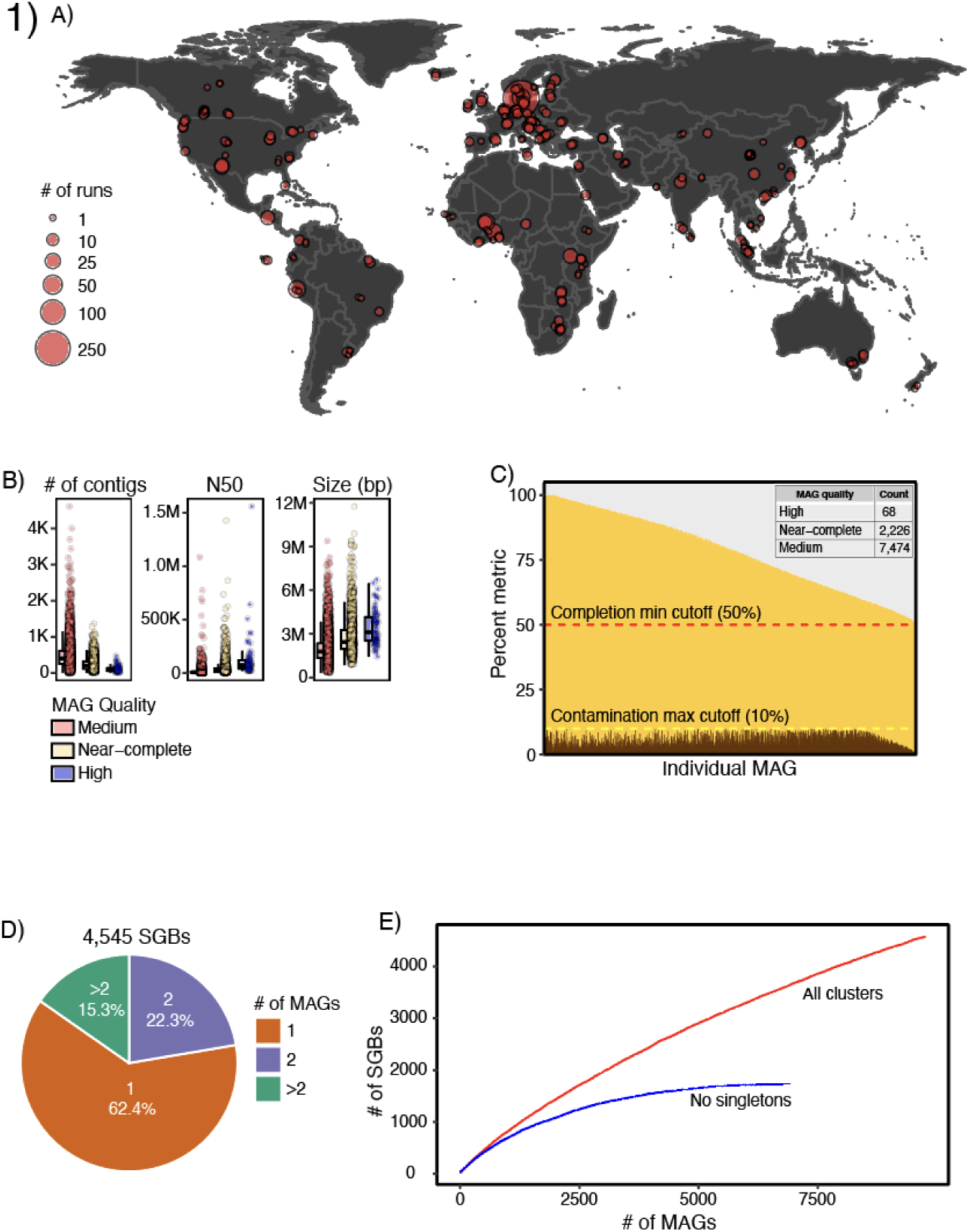
4,545 SGBs recovered from assembled contigs. (A) Geographic distribution of untreated metagenomic wastewater samples used to recover MAGS. (B) Distribution of MAG sizes using number of contigs, N50, and total bp, colored by MAG quality. (C) Non-chimeric MAG genome completeness and contamination. (D) Results from cluster all non-chimeric >= medium quality MAGs to 95% ANI. Accumulation curves of MAG species (E) created by random sampling without replacement from MAGS.

We assembled the sequence data (outlined in **Supplementary Figure 1**) by combining and co-assembling samples from same site based on available metadata (see Methods). This approach yielded 139 successful co-assemblies (comprising 1,062 samples) and 129 individual assemblies, generating over 56 million contigs larger than 1kb and 516 million contigs smaller than 1kb (**Supplementary Figure 2**) ^20^ ^20^. Prior to binning, we removed all contigs identified as lytic viruses to minimize the formation of host–virus chimeric bins to ensure the recovery of high-quality MAGs. The majority of these viral contigs were bacteriophages with 88% classified as *Caudoviricetes*, which were excluded from downstream MAG analyses. As a result, we recovered a total of 12,869 MAGs comprising over 5.5 million contigs. After removing any potentially chimeric MAGs as flagged by GUNC ^21^, we retained a total of 9,768 MAGs (representing over 4 million contigs). The recovered MAGs varied widely in assembly contiguity and size, with contig counts ranging from 4 to 4,612 contigs, N50 values from 1.37 Kbp to 1.6 Mbp, and total genome sizes from 326 Kbp to 11.8 Mbp per MAG **(Figure 1B)**. According to the MIMAG standards^22^, 68 MAGs were classified as high quality (>90% completeness, <5% contamination, and the presence of 16S, 23S, and 5S rRNA genes along with at least 18 tRNAs). A total of 9,670 MAGs were considered medium quality (50–90% completeness and <10% contamination). Among these, 2,196 genomes can be regarded as near-complete, as they exhibit >90% completeness and <5% contamination but lack one or more of the required rRNA genes and/or the full complement of tRNAs necessary to meet the high-quality designation^23^ (**Figure 1C**).

### Species-level genomic bins reveal redundancy in recovered genomes

Prokaryotic MAGs were clustered at 95% average nucleotide identity (ANI) to delineate species-level genomic bins (SGBs) and collapse redundant genomes, resulting in 4,545 SGBs. Although over 9,000 MAGs were recovered, these genomes corresponded to approximately half as many species, reflecting substantial redundancy at the species level. While the majority of SGBs (62%, n=2,835) were represented by a single MAG, the remaining 1,710 SGBs contained 72% of all recovered MAGs. This uneven distribution indicates that multiple MAGs were recovered from many prevalent wastewater species, providing wide representation within these lineages (**Figure 1D**).

To evaluate how comprehensively our genome recovery captured species-level diversity, we performed a species accumulation analysis using SGBs (**Figure 1E)**. When all SGBs were considered, the accumulation curve did not reach saturation, indicating that our dataset, despite being extensive, has not fully captured the total diversity present in wastewater. However, after excluding singleton SGBs, the curve approached a plateau at approximately 3,247 MAGs (1,360 species), corresponding to the recovery of one additional species for every 10 MAGs sampled. Species saturation reached 95% after sampling 5,068 MAGs (1,622 species), suggesting that despite heterogeneous sampling and sequencing strategies, our dataset captures the genomes of many abundant and widely distributed prokaryotic species in wastewater.

### The majority of SGBs represent novel species

To assess the novelty of the recovered SGBs, we compared their similarity to reference genomes using two complementary approaches. First, we compared all SGBs to 429,327 prokaryotic reference genomes comprised of NCBI RefSeq release 227 (n=389,830 genomes) as well as genomes from other environments relevant to wastewater microbial diversity, including human gut (n=13,714 genomes), activated sludge (n=4,706 genomes), and soil (n=21,077 genomes)^24–28^. Using a species-level threshold of 95% ANI, only 874 SGBs clustered with any reference genome, indicating recovered SGBs lack representative species in current genome databases **(Figure 2A)**. Second, we assigned taxonomy to all SGBs using GTDB-Tk^29^. This approach assigned 38% of SGBs (n=1,718) to species-level classification, 750 of which were already assigned by species level clustering. In total, we were able to assign 1,841 SGBs to species a level classification, further underscoring the high number of novel species within the dataset (**Figure 2B**). We then classified an SGB as “known” (kSGB) if it was assigned a species-level designation by either GTDB-Tk or clustered at ≥95% with a reference genome. Based on this definition, 41% (n=1,841) of SGBs were considered known while the remaining 59% (n=2,704) were designated as “unknown” SGBs (uSGBs) (**Figure 2C**). Notably, only 202 SGBs matched species-level references from the human gut, suggesting that much of the wastewater microbial community is distinct from gut-associated taxa.

**Figure 2.**
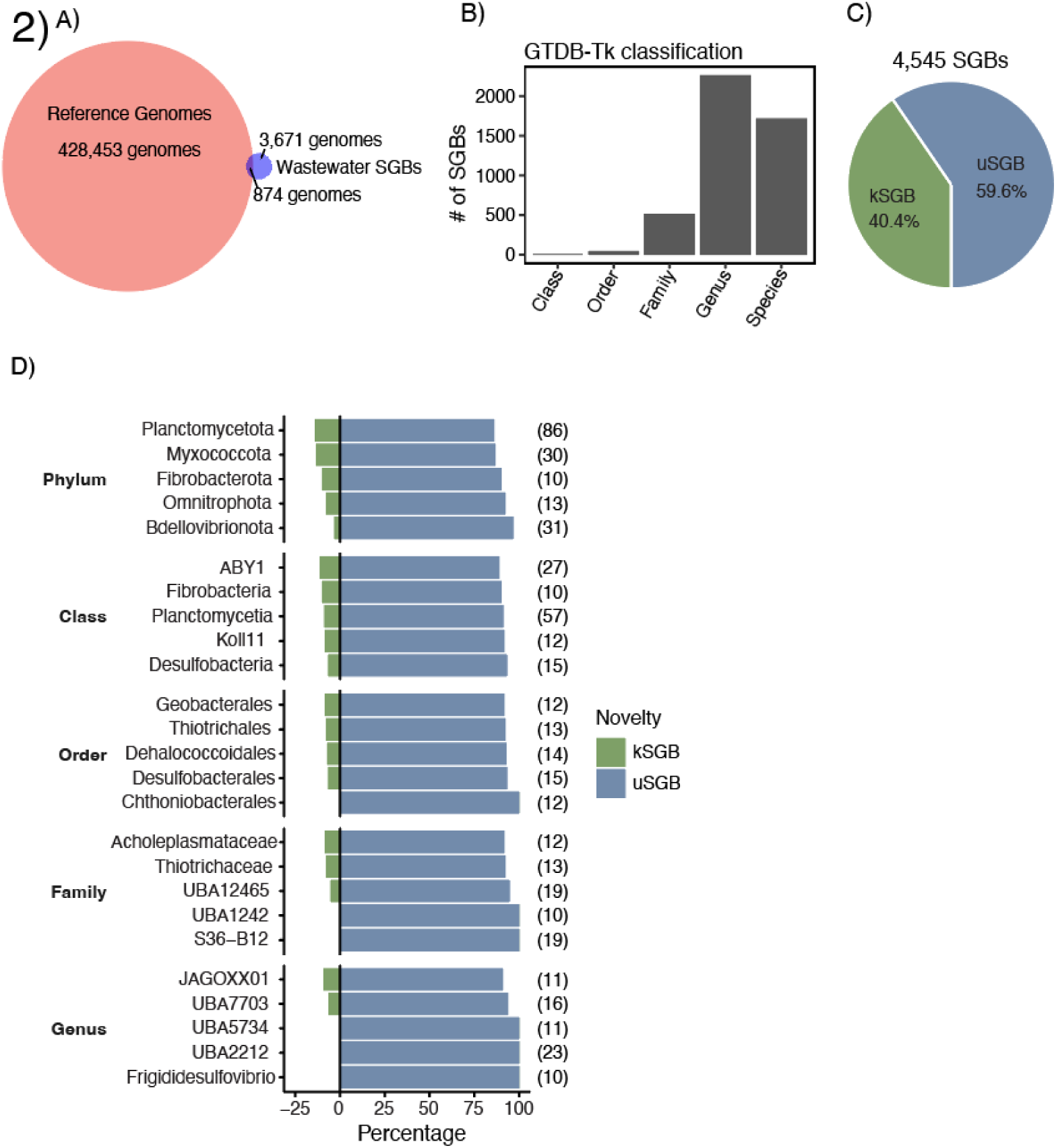
Many novel species recovered from wastewater. (A) Depth of GTDB-Tk classification of all SGBs. (B) Summary of SGBs with or without an SGB in reference databases from RefSeq v227, soil, human gut, and activated sludge. (C) Percentages of kSGBs and uSGBs within all SGBs, where kSGBs were either classified to species level by GTDB-Tk or clustered with a reference genome. (D) Top five most enriched taxa in uSGBs as a percentage of total SGBs per taxonomic level. Numbers in parentheses indicate the number of total SGBs for each taxon.

### uSGBs are from well-established lineages

Despite the novelty at a species level, our recovered uSGBs are predominantly affiliated with well-characterized microbial lineages. From phylum-to family-level, the most common taxa remain surprisingly similar (**Supplementary Figure 3**), indicating that species-level novelty primarily arises from unsampled diversity within established lineages rather than from deeply divergent or rare lineages. Consistent with this interpretation, the majority of uSGBs (79%) were assigned to established genera by GTDB-Tk (**Figure 2B; Supplementary Figure 4).** Despite these similarities, certain taxa from phylum-to genus-level still contributed far more uSGBs than kSGBs, revealing significant unexplored diversity of wastewater microorganisms **(Figure 2D)**.

To assure that the novelty assessment based on GTDB-Tk classification in uSGBs are not due to quality of genomes, we compared uSGBs to kSGBs in genome quality metrics using completeness and contamination. Quality profiles were comparable between known and unknown SGBs (**Supplementary Figure 5**), demonstrating that species novelty is not driven by low-quality genomes, but instead reflects the underrepresentation of wastewater-specific microbial genomes in reference databases.

### Diverse genomic diversity is represented in SGBs

To assess the genomic diversity of the recovered SGBs, we constructed separate phylogenies for bacterial and archaeal SGBs using GTDB-Tk (**Figure 3; Supplementary Figure 6**). We found that bacterial kSGBs and uSGBs were, for the most part, evenly distributed across the phylogenetic tree, indicating that uncharacterized diversity is not confined to a few lineages but rather spans much of the diversity across the bacterial tree. At every taxonomic level below kingdom, uSGBs encompassed a broader span of unique taxa than kSGBs. For instance, uSGBs were distributed across more phyla (53 vs. 38) and genera (1,026 vs. 768), with similar trends observed at intermediate ranks including class, order, and family. Although uSGBs and kSGBs shared similar high-level taxonomic composition, we found that there were several taxa where recovered uSGBs were enriched compared to kSGBs (**Figure 2D**), highlighting some of the abundant members of the wastewater microbiome lacking representative genomes in databases. This broad distribution suggests that wastewater contains representatives of many lineages that remain poorly characterized at many taxonomic levels, highlighting both the limitations of current reference genome databases and the need for additional characterization of genomes from wastewaters.

**Figure 3.**
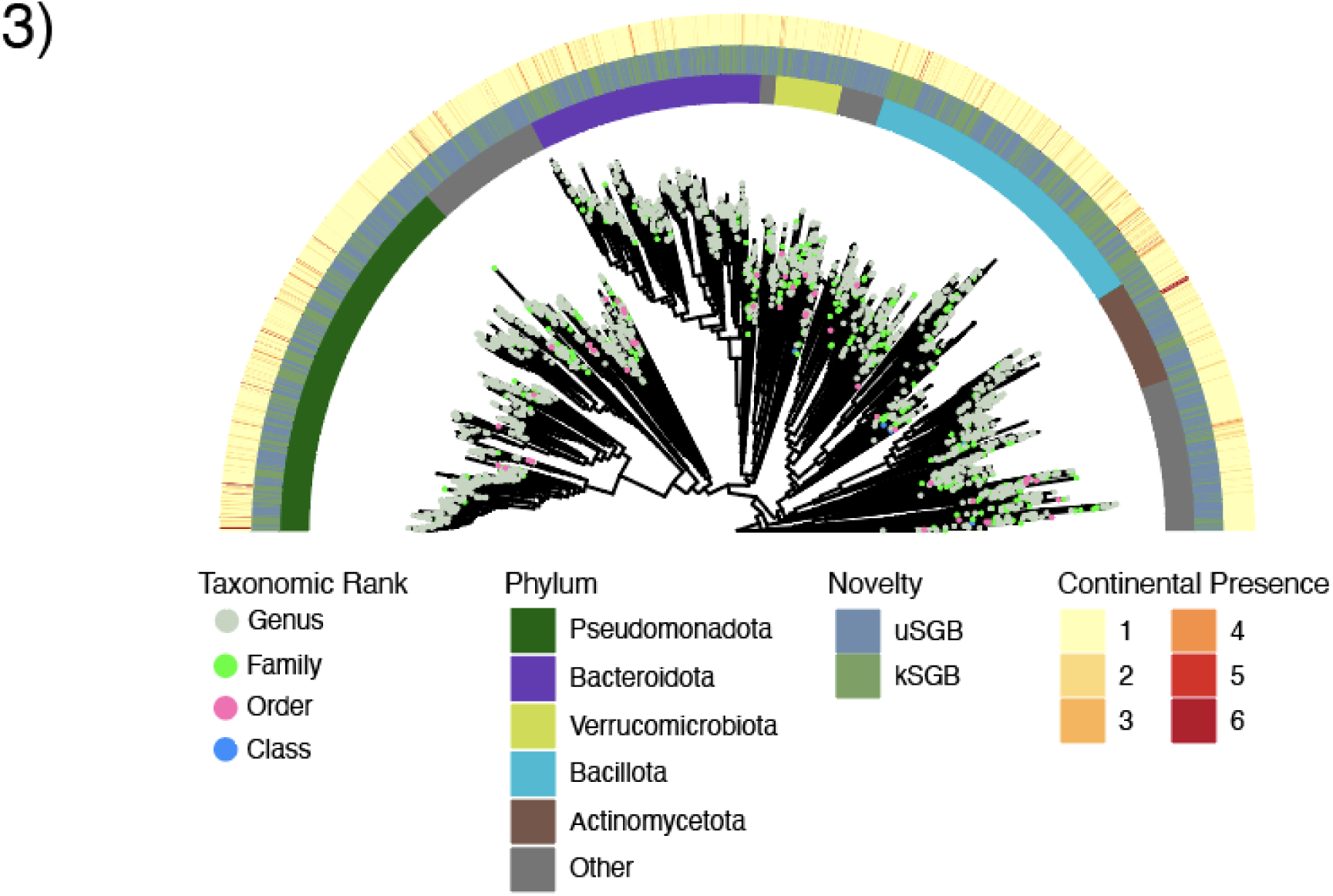
Phylogeny of SGBs in wastewater. Tip colors represent the assigned taxonomic rank of each SGB excluding species. The inner ring indicated the top five most common phyla, the middle represents known or unknown status of the SGB, and the outer ring indicates on how many continents the species was present.

### Putative virulence genes in subset of recovered SGBs

Wastewaters contain inputs from diverse human, animal, and environmental sources, making it a reservoir where pathogenic microbes are also present^30^. Although we cannot determine pathogenicity with certainty using solely bioinformatic analyses, screening for virulence genes provides an important first step in identifying organisms potentially relevant to public health. Detecting SGBs that harbor virulence-associated genes allow us to highlight potential risks and prioritize them for experimental validation and clinical monitoring. To this end, we assessed all SGBs for virulence potential using ABRicate^31^ to screen for the presence of known virulence-associated genes from the VFDB database^32^. Overall, 236 SGBs (5%) contained at least one virulence gene, with an average of 4.5 genes per genome (median = 1) (**Figure 4A**). These putative pathogenic SGBs were distributed across both uSGBs and kSGBs (**Figure 4B**), suggesting that the diversity of organisms in general and potential pathogens remains substantially underrepresented in existing genome resources. The majority of uSGBs harboring virulence genes belonged to the *Pseudomonadota* phylum (93%, n=105) spanning diverse genera within the Alpha- and Gamma-proteobacteria classes (**Figure 4C**).

**Figure 4.**
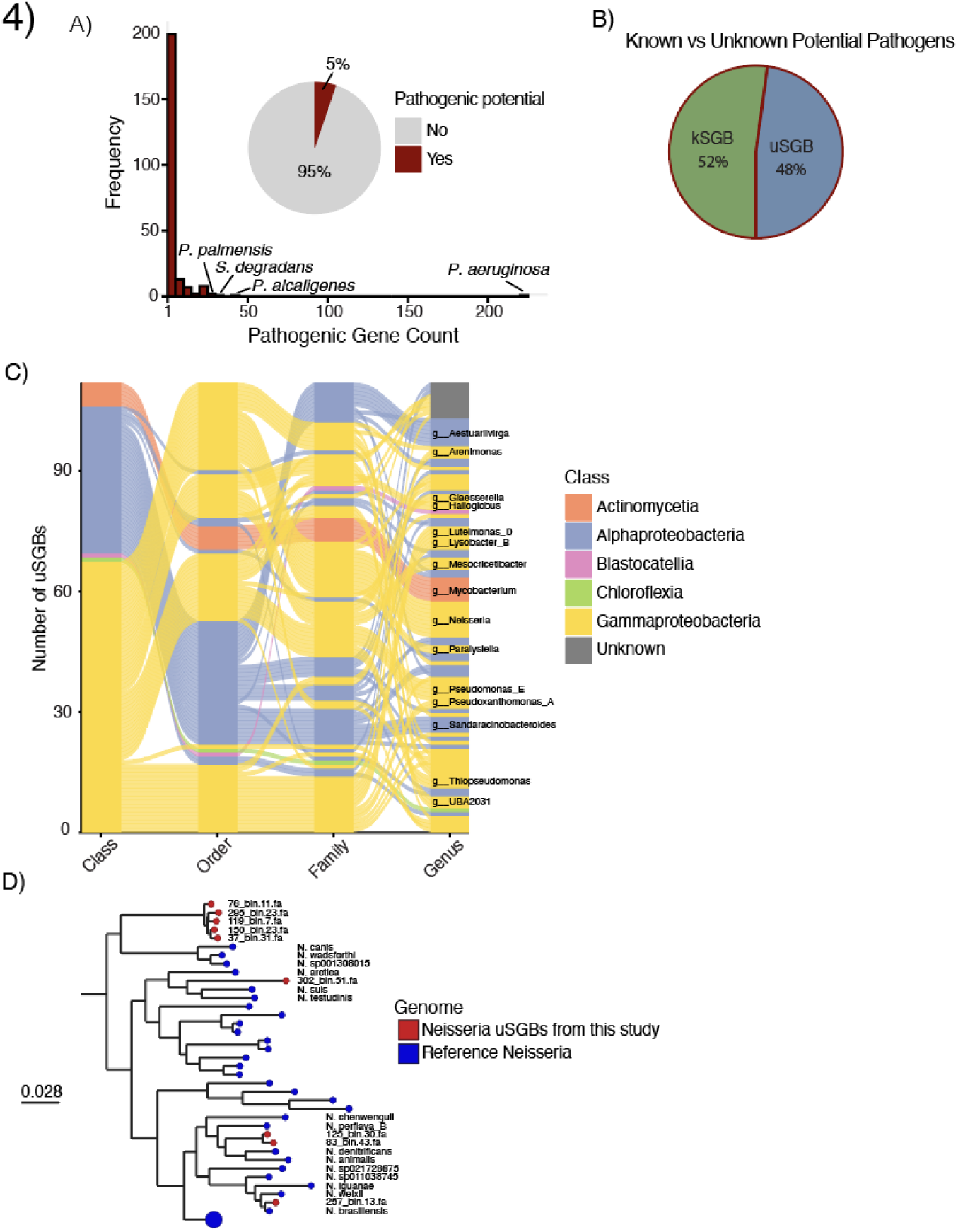
Pathogenic potential of SGBs. (A) Histogram of pathogenic gene counts from each SGB. A *Pseudomonas aeruginosa* genome contained 223 pathogenic genes. (B) Potential pathogens show a near 50/50 split between known and unknown SGBs. (C) Taxonomy of the 113 unknown potential pathogenic species with the most common 15 genera labelled. (D) Phylogenetic placement of unknown Neisseria species with reference Neisseria genomes and Moraxella as the outgroup.

### Potential novel pathogenic lineages in wastewater

While gene-based screening provides an initial indication of pathogenic potential, phylogenetic context offers additional evidence on how closely recovered SGBs are related to known pathogens. To explore this, we focused on the genus *Neisseria*, which includes human and zoonotic pathogens. Among the recovered uSGBs, nine *Neisseria* genomes encoded at least one virulence-associated genes. Their phylogenetic placement with publicly available genomes revealed two distinct patterns, five *Neisseria* uSGBs formed a deeply branching clade that was clearly separated from publicly available *Neisseria* genomes, whereas the remaining uSGBs were interspersed among known *Neisseria* spp. that are zoonotic or human related bacterium (**Figure 4D**). For example, one of the uSGB clustered with a clade that contain zoonotic *Neisseria* species like *N. suis*^33^*, N. testudinis*^34^, and *N.arctica*^34^, while others grouped with known human pathogen *N. brasilensis*^35^. Taken together, the example with *Neisseria* suggests that many uSGBs carrying virulent genes may correspond to either close relatives of known pathogens or to previously undescribed clades within taxonomic lineages.

### Most SGBs are continent-specific

To assess global distribution of SGBs, we examined their occurrence at the continental level. Continental presence was determined for each SGB based on the number of continents in which its member MAGs were detected, with detection defined as the recovery of at least one MAG from a sample collected on that continent. We then classified SGBs as continent-specific (detected in one continent), multi-continental (detected in two to five continents), or globally distributed (detected across all six continents). The majority of recovered SGBs were continent-specific (n=3,981), whereas 538 were multi-continental, and only 26 were globally distributed (**Figure 5A**). Continental specificity was quite pronounced as 83% of SGBs in North America, 86% in Europe, 90% in Asia, 92% in Africa, 78% in South America, and 73% in Oceania were unique to their continent of origin. Asia contained the largest number of unique SGBs (n = 1,555) despite representing only 13% of total samples, reflecting broader sampling coverage from 19 countries.

**Figure 5.**
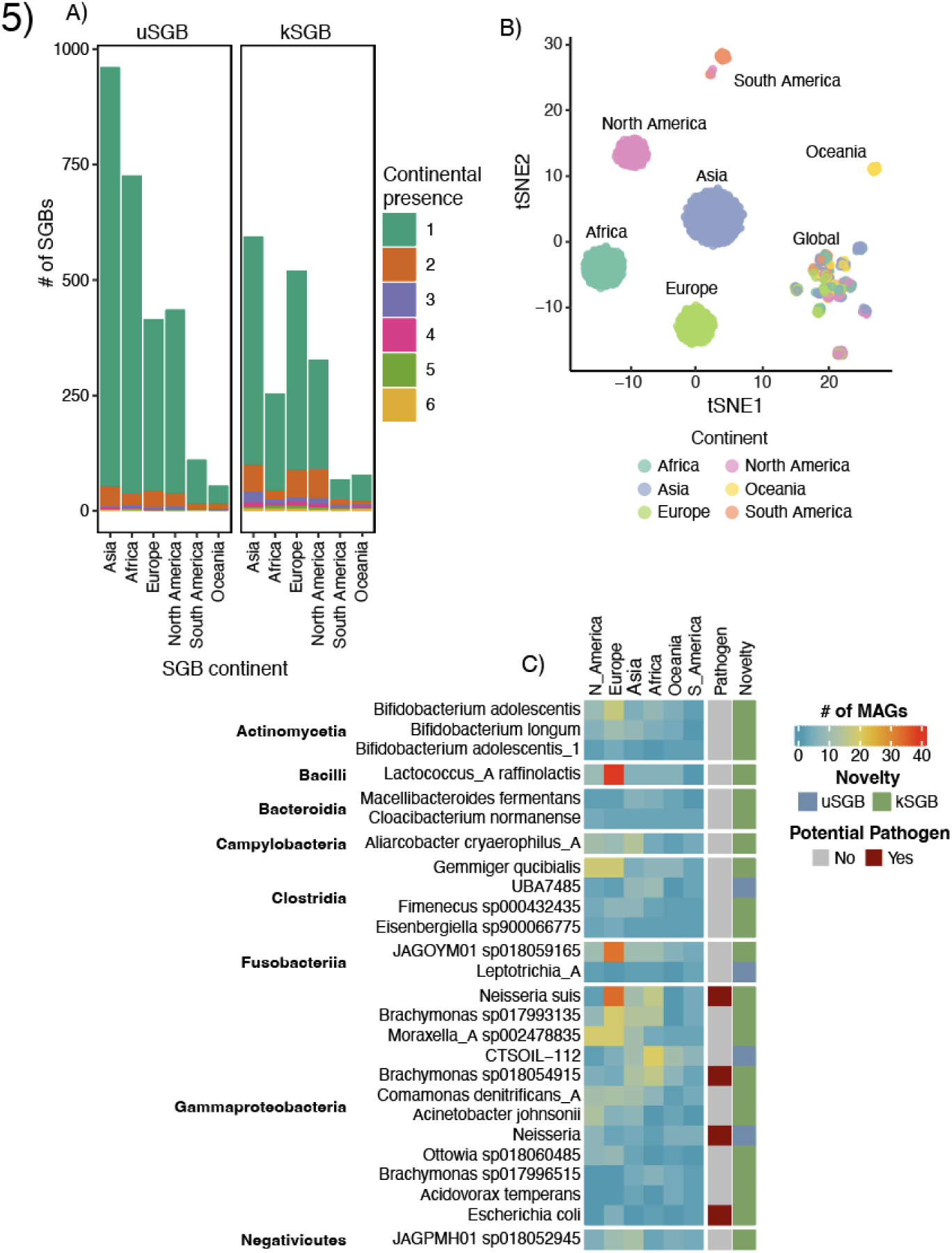
Global and continental cores of wastewater. (A) continental presence of SGBs, determined by the MAGs within each SGB. (B) UMAP made from binary presence/absence matrix for each SGB. (C) Heatmap showing frequencies of 26 core SGBs on each continent.

In contrast, Oceania (133 SGBs, 3% of samples) and South America (179 SGBs, 8% of samples) contributed the fewest due to more limited sampling, highlighting geographic biases in the dataset. To further explore these geographic patterns, we constructed a tSNE plot based on a binary matrix of SGB presence across continents (**Figure 5B**). This analysis revealed clustering influenced by most SGBs being continent-specific, and additionally we observed a distinct core of multi-continent and globally distributed SGBs, suggesting broad differences in SGB’s species level diversity that were recovered among continents. We used raw counts for a more detailed examination of continental distributions (**Supplementary Figure 7)**, and the results aligned with Figures 5A and 5B, showing a clustering of SGBs found on most continents and high continental specificity outside of that core. Likewise, the majority of uSGBs within each continent were continent specific as well. For example, Africa had the highest proportion of uSGBs (77% of 900), whereas Oceania had the lowest (30% of 133). These results indicate that continent-specific SGBs were enriched in uSGBs, consistent with the expectation that many uSGBs represent low-prevalence species in wastewater. The high proportion of uSGBs that are continent-specific, particularly in Africa, also points to substantial regional microbial diversity that is absent from current reference databases.

Screening for virulence genes showed that potential pathogenic SGBs did not exhibit a strong geographic bias. Instead, the proportion of pathogenic SGBs was relatively consistent across continent, ranging from 8% in Oceania to 4% in North America, Africa, and South America (**Supplementary Table 3**). A heatmap of only pathogenic SGBs and their presence on each continent showed similar patterns **(Supplementary Figure 8**). Together, these results indicate that most wastewater species are geographically restricted, with only a small set of globally distributed core taxa.

### A small set of SGBs represent the global core

Despite the predominance of continent-specific SGBs, a small subset of 26 SGBs was consistently recovered from wastewater samples across all six continents (**Figure 5C**). These globally distributed SGBs represent a robust core of wastewater-associated species that were detectable across highly heterogeneous sampling efforts and geographic regions. The majority of these core SGBs were classified as known species (kSGBs), indicating that widely distributed and consistently recoverable wastewater taxa are better represented in current reference genome databases. Remarkably, one *Neisseria* SGB that was not classified to a species encoded virulence-associated genes, suggesting that even among the most prevalent wastewater species, genomic diversity and potential public-health relevance remain incompletely characterized.

## Discussion

Our analysis of 1,344 wastewater metagenomes spanning 79 countries has revealed an unprecedented genome-resolved view of microbial diversity in untreated wastewater from around the globe, despite remaining focused fraction of microbial populations that could be assembled at sufficient resolution to form MAGs. The recovery of 9,768 MAGs representing 4,545 species-level genome bins (SGBs) constitutes one of the largest collections of wastewater-associated prokaryotic genomes to date. Most notably, our findings demonstrate that wastewater harbors extensive microbial diversity that remains largely uncharacterized, with 60% of recovered SGBs representing previously unknown species.

### Novel Microbial Diversity in Wastewater

The high proportion of novel species identified in this study highlights wastewater as a largely uncharacterized reservoir of microbial diversity. Despite comparing our SGBs against an extensive reference database of over 429,000 prokaryotic genomes, including those from environments relevant to wastewater (human gut, activated sludge, and soil), only 40% could be classified as known species. The broad phylogenetic distribution of these novel species, which spans 53 phyla compared to 38 for known species, indicates that this uncharacterized diversity is not confined to specific lineages but rather represents a systematic gap in our understanding of wastewater microbiomes. Interestingly, only 4.5% (n=202) SGBs matched species-level references from the human gut microbiome, challenging the common assumption that wastewater microbial communities primarily reflect human fecal inputs.

### Geographic Distribution and Continental Specificity

Understanding the geographic distribution of wastewater SGBs is important for guiding public health efforts including pathogen detection, antimicrobial resistance surveillance, and wastewater treatment optimization^13,36–38^. Our findings reveal pronounced patterns of geographic specificity in wastewater microbiomes, with 88% of SGBs restricted to a single continent. This high degree of endemicity suggests that regional factors strongly influence wastewater microbial communities. The observation that Africa harbors the highest proportion of unknown SGBs (77%) is particularly noteworthy and likely reflects historical sampling biases in microbial genomics, which has focused disproportionately on high-income regions. This emphasizes the need for geographically broad sampling to capture the full extent of global wastewater microbiomes.

Our analysis revealed a comparatively small set of globally distributed (core) SGBs which contrasts with a recent study that reported a much larger core community^13^. This difference largely arises from how “core” is defined and measured. Whereas the authors identified a prevalence-based core comprising taxa present in over 70% of individual samples, our study defines a biogeographic core requiring species-level SGBs to be detected across six continents. Our more stringent, distribution-based definition emphasizes true global ubiquity rather than regional dominance. Together, these differences highlight that widespread wastewater taxa are relatively rare, and most microbial species exhibit strong continental specificity, underscoring pronounced regional structure within the global wastewater microbiome.

### Wastewater as a pathogen reservoir

We identified 236 SGBs harboring virulence genes highlighting wastewater’s potential role as a reservoir for pathogen diversity. Nearly half of these putative pathogenic SGBs represent previously unknown species, which highlights the utility of using wastewater to monitor for new and emerging threats (**Figure 4B**). Importantly, the relatively even distribution of putative pathogenic SGBs across continents indicates that pathogen presence in wastewater is a global phenomenon rather than regionally restricted. This underscores the potential of wastewater surveillance as a tool for pathogen monitoring worldwide.

### Future directions

This study establishes a foundation for future research into wastewater microbiomes and their applications. The extensive catalog of novel SGBs provides reference genomes that can improve metagenomic analyses of wastewater samples worldwide. Future work should focus on characterizing the functional capabilities of these novel species, particularly their roles in nutrient cycling, pollutant degradation, and microbial ecology within wastewater systems. For public health applications, experimental validation of the pathogenic potential of novel SGBs identified in this study should also be investigated. It is also worth noting that although this study presents the largest publicly available collection of archaeal and bacterial MAGs recovered from wastewater, a substantial amount of diversity likely remains undiscovered. Future work should also focus on further characterization of wastewater microbiome diversity through equitable sampling across the globe. Expanding these analyses to examine strain-level assemblies and binning would allow greater resolution for tracking pathogens, virulence factors, and antimicrobial resistance mechanisms. Finally, it is important to note that our analyses focused on MAGs which represent only a subset of the assembled sequence space. The majority of contigs ≥1kb (∼55 million) were not incorporated into MAGs (∼5 million), likely corresponding to microbial populations present at an abundance insufficient for genome reconstruction. This observation suggests that substantial microbial diversity in wastewater remains uncharacterized and highlights the need for deeper and more intentional sampling strategies to more fully resolve the background wastewater microbiome.

In conclusion, our study reveals wastewater as a vast reservoir of previously uncharacterized microbial diversity with significant implications for microbial ecology, wastewater treatment, and public health monitoring. The global wastewater microbiome represents not only a challenge for comprehensive characterization but also an opportunity for discovering novel microorganisms, functions, and potential pathogens that impact human and environmental health worldwide.

### Funding and Acknowledgements

This work was funded by the Laboratory Directed Research and Development grants 202205855ECR to M.S and 20240674ER to M.S and P.C.

## Methods

### Sequence Read Archive data selection

From the NCBI Sequence Read Archive (SRA), all datasets with metadata containing keywords “metagenome” and “wastewater” were selected on May 17^th^ 2022. Approximately 40K biosamples were downloaded. Biosamples were removed if descriptors included keywords such as 16S, effluent, soil, sludge, and others that indicated targeted sequencing or samples from non-wastewater or treated wastewater (e.g. effluent wastewater), detailed in **(Supplementary Table 2**). Removed biosamples were also manually confirmed to ensure correct removal. This process did not remove all unwanted biosamples due to incomplete metadata, so we screened the remaining samples manually using available metadata and corresponding literature verifications, when available. The entire filtering process reduced our database to high quality 1,344 biosamples that exclusively contains untreated wastewater **(Supplementary Table 1)**.

### Data QC and assembly

Raw fastq files were downloaded using prefetch and fasterq-dump commands with default parameters from sratoolkit v3.0.0^39^. FaQCs v2.09^40^ was run with trim level 20 on reads to be single-assembled and not run on reads to be co-assembled. Specifically, we co-assembled samples, when they belonged to the same NCBI Bioproject and were collected from the same sources and locations (**Supplementary Table 1**). However, samples with incomplete location metadata were assembled individually, even if from the same Bioproject. Reads were single-assembled using Spades v3.13.0^41^ with the –meta flag. MetaHipMer v2.2.0^41^ with default parameters was used for co-assembly of paired-end reads and MEGAHIT v1.2.9^42^ with the – meta-large parameter was used for co-assembly of single-end reads.

### Removing viral contigs

Putative viral contigs were identified from assemblies using VIBRANT v1.2.1^20^ on default settings. The contigs labeled as lytic and medium, high, or complete were removed from the original assembly to increase bin quality, as viral contigs may share similar characteristics with bacterial contigs such as abundance, GC content, and k-mer composition which can lead to chimeric MAGs.

### Generation of MAGs

Non-viral contigs were binned using metaWRAP v1.3.2^43^ with the *binning* and *bin_refinement* modules to recover MAGs. To assign taxonomy to MAGs, GTDB-Tk v2.3.0^29^ *de_novo_wf* was used with default parameters and *GTDB release 214*. For additional QC on the refined bin set, GUNC v1.0.5^21^ was run on default parameters to detect and remove chimeric MAGs from the dataset. Quality of the bins were assessed using CheckM v1.0.2^44^.

### Genome clustering to SGBs

Genomes clustered to species-level bins with 95% ANI using dRep v3.4.3^45^ with parameters *–pa 0.9, -sa 0.95, -nc 0.30, -cm larger, and –S_algorithm fastANI.* After clustering MAGs at the species level, representative SGB genomes were selected based on maximal taxonomic resolution and highest genome quality. For clusters containing a single MAG, that MAG was automatically designated as the SGB representative.

### Comparison of SGBs to relevant database

An SGB was considered known if it had 95% ANI to a reference genome or was given a species-level classification by GTDB-Tk. Specifically, representative SGBs were compared to database containing relevant genomes and MAGs using MASH v2.3^46^ and dRep v3.4.3. First, all genomes were sketched using mash sketch with default parameters, then sketches of all genomes from the same database merged into a single database (using mash-paste) on default parameters to allow faster pairwise comparisons. Mash distances were calculated between SGBs and reference genomes using *mash dist* on default parameters, and hits were filtered to a mash distance <= 0.5. If an SGB had multiple hits to a reference genome at this threshold, the lower mash distance value was chosen. Each SGB was then compared to its reference genome partner using dRep parameters detailed above to more accurately assess similarity.

### Predicting potential pathogens

We ran ABRicate v1.0.1^31^ on all SGBs using the *-vfdb* parameter. SGBs containing at least one pathogenic gene were considered to be a potential pathogen.

### Phylogenetic analysis

To construct phylogenies from the bacterial and archaeal SGBs, we used GTDB-Tk v2.3.0. “*gtdbtk identify*” to identify marker genes and their respective proteins in all MAGs, which were then aligned with “*gtdbtk align”*. Both GTDB-Tk commands were run with default parameters except when excluding the reference genomes during the align step. The tree was visualized with ggtree v3.10.1^47^. For construction of the *Neisseria* phylogeny, we used GTDB-Tk v2.3.0 “de novo” workflow with *g Moraxella* as the outgroup and visualized using Microreact^48^. Branches with only reference genomes were collapsed, with the full tree in **Supplementary Figure 9.**

### tSNE clustering

In R v4.4.1^49^, we made a tSNE of SGBs based on a presence or absence matrix of continental presence using Rtsne package v0.17^50^ with perplexity=30. To ensure confidence, we tested 10 different seeds and altered parameters and the general pattern of clustering remained consistent for each iteration.

## References

1 Raymond, S. et al. How local health departments use wastewater surveillance data for public health planning and intervention in New York State. BMC Public Health 25, 2842 (2025). 10.1186/s12889-025-24250-6

2 van der Drift, A. R. et al. Wastewater surveillance studies on pathogens and their use in public health decision-making: a scoping review. Sci Total Environ 993, 179982 (2025). 10.1016/j.scitotenv.2025.179982

3 Diamond, M. B. et al. Wastewater surveillance of pathogens can inform public health responses. Nat Med 28, 1992–1995 (2022). 10.1038/s41591-022-01940-x

4 Lee, B. K. et al. Influence of wastewater type on the distribution of microbial community compositions including pathogenic bacteria within wastewater treatment processes. Sustain Environ Res 35 (2025). ARTN17 10.1186/s42834-025-00253-1

5 Ko, K. K. K., Chng, K. R. & Nagarajan, N. Metagenomics-enabled microbial surveillance. Nat Microbiol 7, 486–496 (2022). 10.1038/s41564-022-01089-w

6 Munk, P. et al. Genomic analysis of sewage from 101 countries reveals global landscape of antimicrobial resistance. Nat Commun 13, 7251 (2022). 10.1038/s41467-022-34312-7

7 Tierney, B. T. et al. Towards geospatially-resolved public-health surveillance via wastewater sequencing. Nat Commun 15, 8386 (2024). 10.1038/s41467-024-52427-x

8 Brown, C. T. et al. Unusual biology across a group comprising more than 15% of domain Bacteria. Nature 523, 208–211 (2015). 10.1038/nature14486

9 Pavlopoulos, G. A. et al. Unraveling the functional dark matter through global metagenomics. Nature 622, 594–602 (2023). 10.1038/s41586-023-06583-7

10 Teudt, F., Otani, S. & Aarestrup, F. M. Global Distribution and Diversity of Prevalent Sewage Water Plasmidomes. mSystems 7, e0019122 (2022). 10.1128/msystems.00191-22

11 Dueholm, M. K. D. et al. MiDAS 4: A global catalogue of full-length 16S rRNA gene sequences and taxonomy for studies of bacterial communities in wastewater treatment plants. Nat Commun 13, 1908 (2022). 10.1038/s41467-022-29438-7

12 Dueholm, M. K. D. et al. MiDAS 5: Global diversity of bacteria and archaea in anaerobic digesters. Nat Commun 15, 5361 (2024). 10.1038/s41467-024-49641-y

13 Yan, Y. et al. Global wastewater microbiome reveals core bacterial community and viral diversity with regional antibiotic resistance patterns. mSystems 10, e0142824 (2025). 10.1128/msystems.01428-24

14 Wyler, E. et al. Pathogen dynamics and discovery of novel viruses and enzymes by deep nucleic acid sequencing of wastewater. Environ Int 190, 108875 (2024). 10.1016/j.envint.2024.108875

15 Begmatov, S. et al. Metagenomic insights into the wastewater resistome before and after purification at large-scale wastewater treatment plants in the Moscow city. Sci Rep 14, 6349 (2024). 10.1038/s41598-024-56870-0

16 Pearson, V. M., Caudle, S. B. & Rokyta, D. R. Viral recombination blurs taxonomic lines: examination of single-stranded DNA viruses in a wastewater treatment plant. PeerJ 4, e2585 (2016). 10.7717/peerj.2585

17 Haryono, M. A. S. et al. Recovery of High Quality Metagenome-Assembled Genomes From Full-Scale Activated Sludge Microbial Communities in a Tropical Climate Using Longitudinal Metagenome Sampling. Front Microbiol 13, 869135 (2022). 10.3389/fmicb.2022.869135

18 Calderon-Franco, D. et al. Metagenomic profiling and transfer dynamics of antibiotic resistance determinants in a full-scale granular sludge wastewater treatment plant. Water Res 219, 118571 (2022). 10.1016/j.watres.2022.118571

19 Brinch, C. et al. Long-Term Temporal Stability of the Resistome in Sewage from Copenhagen. mSystems 5 (2020). 10.1128/mSystems.00841-20

20 Kieft, K., Zhou, Z. & Anantharaman, K. VIBRANT: automated recovery, annotation and curation of microbial viruses, and evaluation of viral community function from genomic sequences. Microbiome 8, 90 (2020). 10.1186/s40168-020-00867-0

21 Orakov, A. et al. GUNC: detection of chimerism and contamination in prokaryotic genomes. Genome Biol 22, 178 (2021). 10.1186/s13059-021-02393-0

22 Bowers, R. M. et al. Minimum information about a single amplified genome (MISAG) and a metagenome-assembled genome (MIMAG) of bacteria and archaea. Nat Biotechnol 35, 725–731 (2017). 10.1038/nbt.3893

23 Parks, D. H. et al. Recovery of nearly 8,000 metagenome-assembled genomes substantially expands the tree of life. Nat Microbiol 2, 1533–1542 (2017). 10.1038/s41564-017-0012-7

24 O’Leary, N. A. et al. Reference sequence (RefSeq) database at NCBI: current status, taxonomic expansion, and functional annotation. Nucleic Acids Res 44, D733–745 (2016). 10.1093/nar/gkv1189

25 Gurbich, T. A. et al. MGnify Genomes: A Resource for Biome-specific Microbial Genome Catalogues. J Mol Biol 435, 168016 (2023). 10.1016/j.jmb.2023.168016

26 Ye, L., Mei, R., Liu, W. T., Ren, H. & Zhang, X. X. Machine learning-aided analyses of thousands of draft genomes reveal specific features of activated sludge processes. Microbiome 8, 16 (2020). 10.1186/s40168-020-0794-3

27 Abdulkadir, N. et al. Genome-centric analyses of 165 metagenomes show that mobile genetic elements are crucial for the transmission of antimicrobial resistance genes to pathogens in activated sludge and wastewater. Microbiol Spectr 12, e0291823 (2024). 10.1128/spectrum.02918-23

28 Ma, B. et al. A genomic catalogue of soil microbiomes boosts mining of biodiversity and genetic resources. Nat Commun 14, 7318 (2023). 10.1038/s41467-023-43000-z

29 Chaumeil, P. A., Mussig, A. J., Hugenholtz, P. & Parks, D. H. GTDB-Tk v2: memory friendly classification with the genome taxonomy database. Bioinformatics 38, 5315–5316 (2022). 10.1093/bioinformatics/btac672

30 Corrin, T. et al. A scoping review of human pathogens detected in untreated human wastewater and sludge. J Water Health 22, 436–449 (2024). 10.2166/wh.2024.326

31 ABRicate (GitHub, 2020).

32 Zhou, S., Liu, B., Zheng, D., Chen, L. & Yang, J. VFDB 2025: an integrated resource for exploring anti-virulence compounds. Nucleic Acids Res 53, D871–D877 (2025). 10.1093/nar/gkae968

33 Busse, H. J., Kampfer, P., Szostak, M. P., Ruckert, C. & Spergser, J. Paralysiella testudinis gen. nov., sp. nov., isolated from the cloaca of a toad-headed turtle (Mesoclemmys nasuta). Int J Syst Evol Microbiol 71 (2021). 10.1099/ijsem.0.005114

34 Hansen, C. M. et al. Neisseria arctica sp. nov., isolated from nonviable eggs of greater white-fronted geese (Anser albifrons) in Arctic Alaska. Int J Syst Evol Microbiol 67, 1115–1119 (2017). 10.1099/ijsem.0.001773

35 Mustapha, M. M. et al. Two Cases of Newly Characterized Neisseria Species, Brazil. Emerg Infect Dis 26, 366–369 (2020). 10.3201/eid2602.190191

36 Khan, M. et al. Significance of wastewater surveillance in detecting the prevalence of SARS-CoV-2 variants and other respiratory viruses in the community - A multi-site evaluation. One Health 16, 100536 (2023). 10.1016/j.onehlt.2023.100536

37 Foxman, B. et al. Wastewater surveillance of antibiotic-resistant bacteria for public health action: potential and challenges. Am J Epidemiol 194, 1192–1199 (2025). 10.1093/aje/kwae419

38 Shi, K. et al. Microbiome regulation for sustainable wastewater treatment. Biotechnol Adv 77, 108458 (2024). 10.1016/j.biotechadv.2024.108458

39 Information, N. C. f. B. *SRA Toolkit*, <https://github.com/ncbi/sra-tools> (

40 Lo, C. C. & Chain, P. S. Rapid evaluation and quality control of next generation sequencing data with FaQCs. BMC Bioinformatics 15, 366 (2014). 10.1186/s12859-014-0366-2

41 Hofmeyr, S. et al. Terabase-scale metagenome coassembly with MetaHipMer. Sci Rep 10, 10689 (2020). 10.1038/s41598-020-67416-5

42 Li, D., Liu, C. M., Luo, R., Sadakane, K. & Lam, T. W. MEGAHIT: an ultra-fast single-node solution for large and complex metagenomics assembly via succinct de Bruijn graph. Bioinformatics 31, 1674–1676 (2015). 10.1093/bioinformatics/btv033

43 Uritskiy, G. V., DiRuggiero, J. & Taylor, J. MetaWRAP-a flexible pipeline for genome-resolved metagenomic data analysis. Microbiome 6, 158 (2018). 10.1186/s40168-018-0541-1

44 Parks, D. H., Imelfort, M., Skennerton, C. T., Hugenholtz, P. & Tyson, G. W. CheckM: assessing the quality of microbial genomes recovered from isolates, single cells, and metagenomes. Genome Res 25, 1043–1055 (2015). 10.1101/gr.186072.114

45 Olm, M. R., Brown, C. T., Brooks, B. & Banfield, J. F. dRep: a tool for fast and accurate genomic comparisons that enables improved genome recovery from metagenomes through de-replication. ISME J 11, 2864–2868 (2017). 10.1038/ismej.2017.126

46 Ondov, B. D. et al. Mash: fast genome and metagenome distance estimation using MinHash. Genome Biol 17, 132 (2016). 10.1186/s13059-016-0997-x

47 Xu, S. et al. Ggtree: A serialized data object for visualization of a phylogenetic tree and annotation data. Imeta 1, e56 (2022). 10.1002/imt2.56

48 Argimon, S. et al. Microreact: visualizing and sharing data for genomic epidemiology and phylogeography. Microb Genom 2, e000093 (2016). 10.1099/mgen.0.000093

49 R: A Language and Environment for Statistical Computing v. 4.4.1 (2024).

50. T-Distributed Stochastic Neighbor Embedding using a Barnes-Hut Implementation v. 0.17 (2023).

