## Supplementary Tables and Figures for "Global untreated wastewater hosts a vast reservoir of previously uncharacterized microbial lineages": global_ww_supp_all.pdf

1)

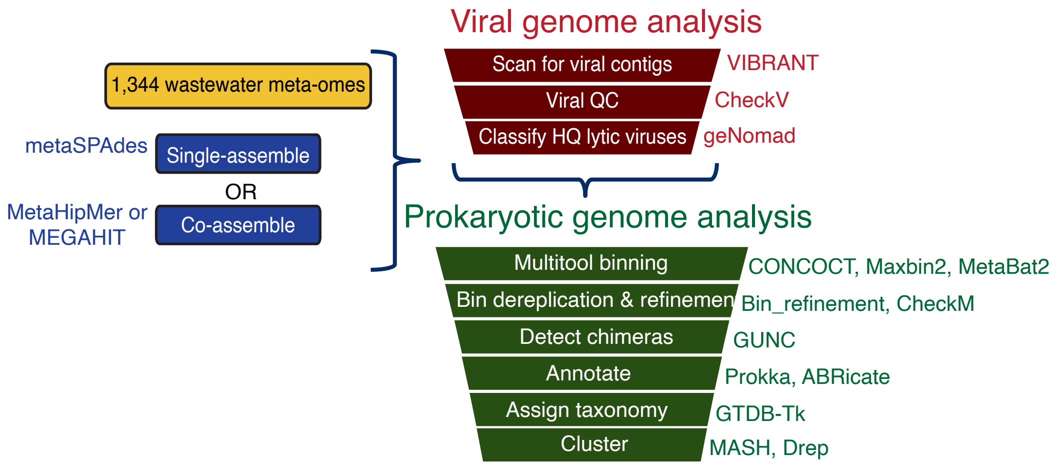

**Supplementary Figure 1**: Bioinformatic pipeline for assembling shotgun metagenomic samples, scanning for viruses, binning, and MAG analysis.

2)

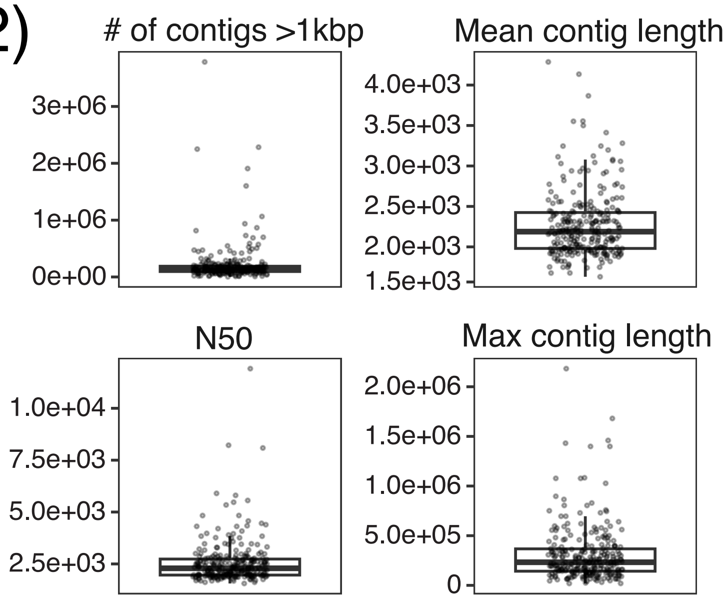

**Supplementary Figure 2:** Statistics of all assembled reads. Each dot represents an assembly.

3)

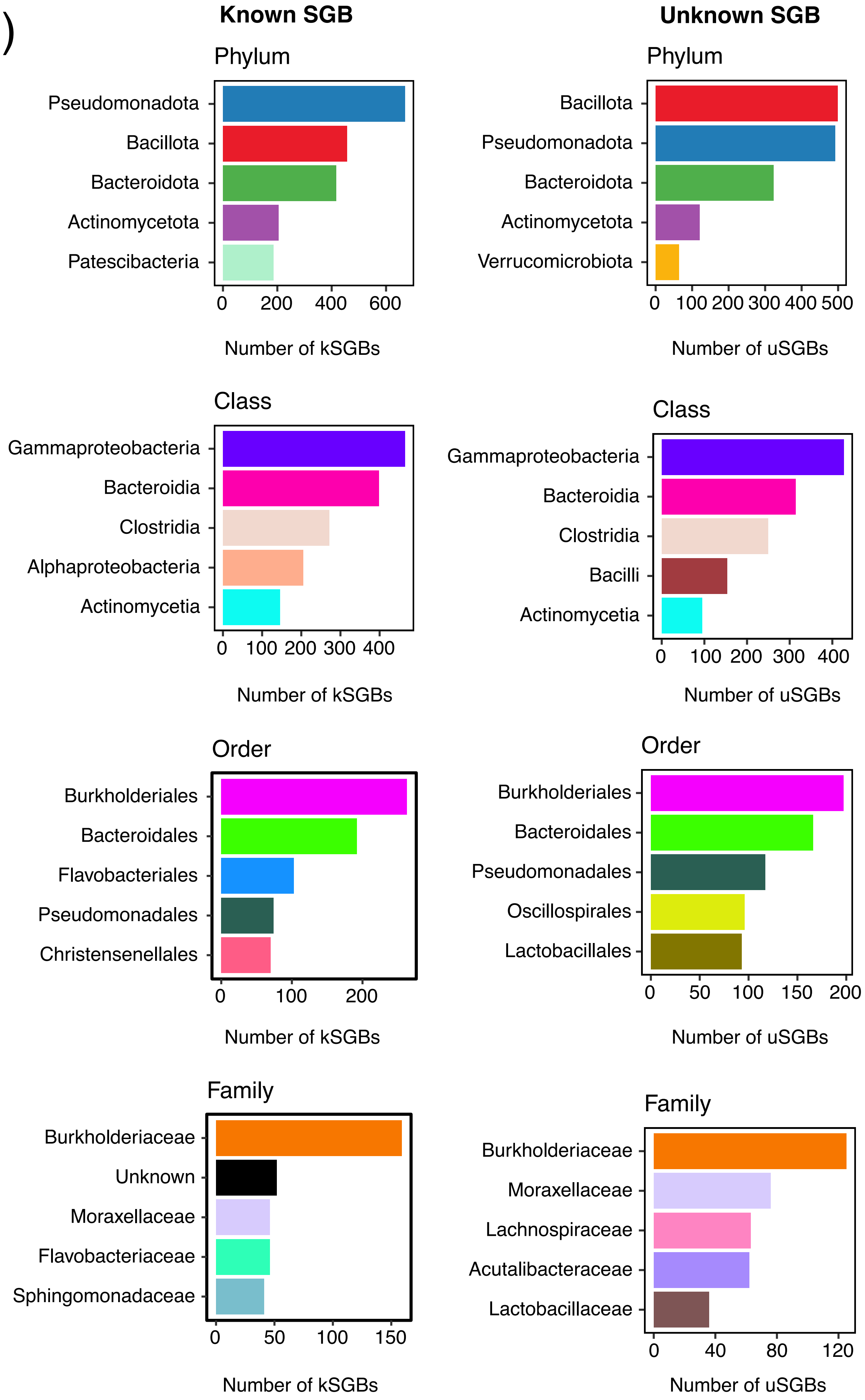

**Supplementary Figure 3:** Top 5 common taxonomic classifications of kSGBs and uSGBs. Genus-level classifications were excluded due to large diversity within each group.

4)

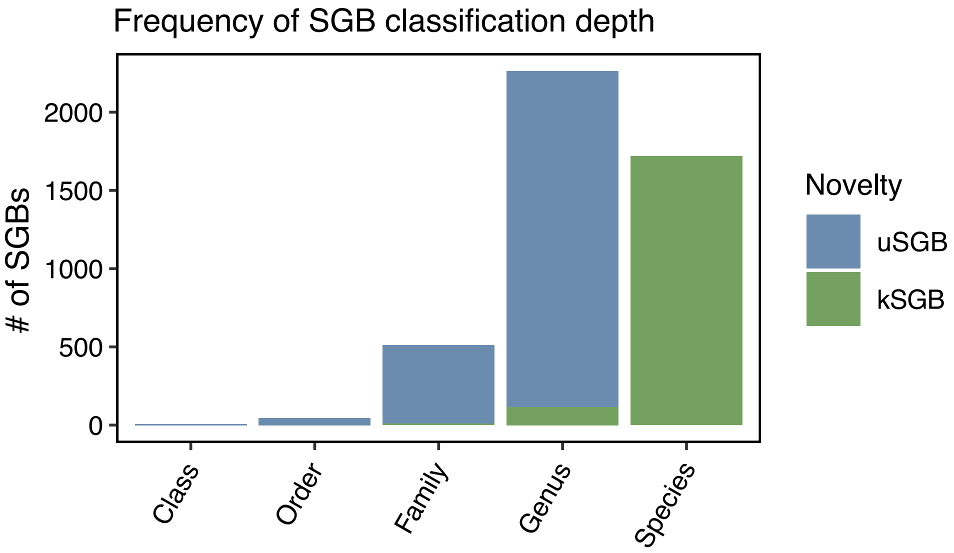

**Supplementary Figure 4:** Frequency of SGB classification depth, colored by known or unknown status. All SGBs clustered to 95% ANI with reference genomes or given a species-level taxonomy by GTDB-Tk were considered known.

5)

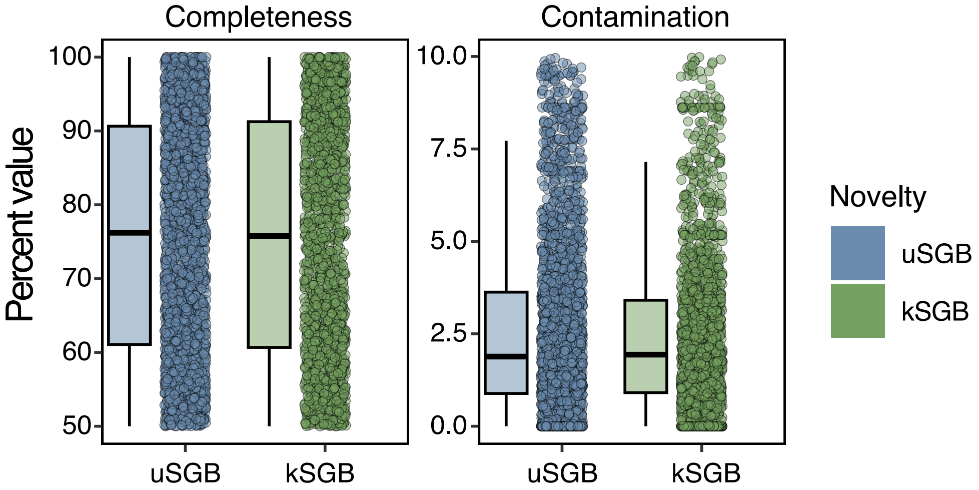

**Supplementary Figure 5:** Completeness and contamination metrics for uSGBs and kSGBs, predicted by CheckM2.

6)

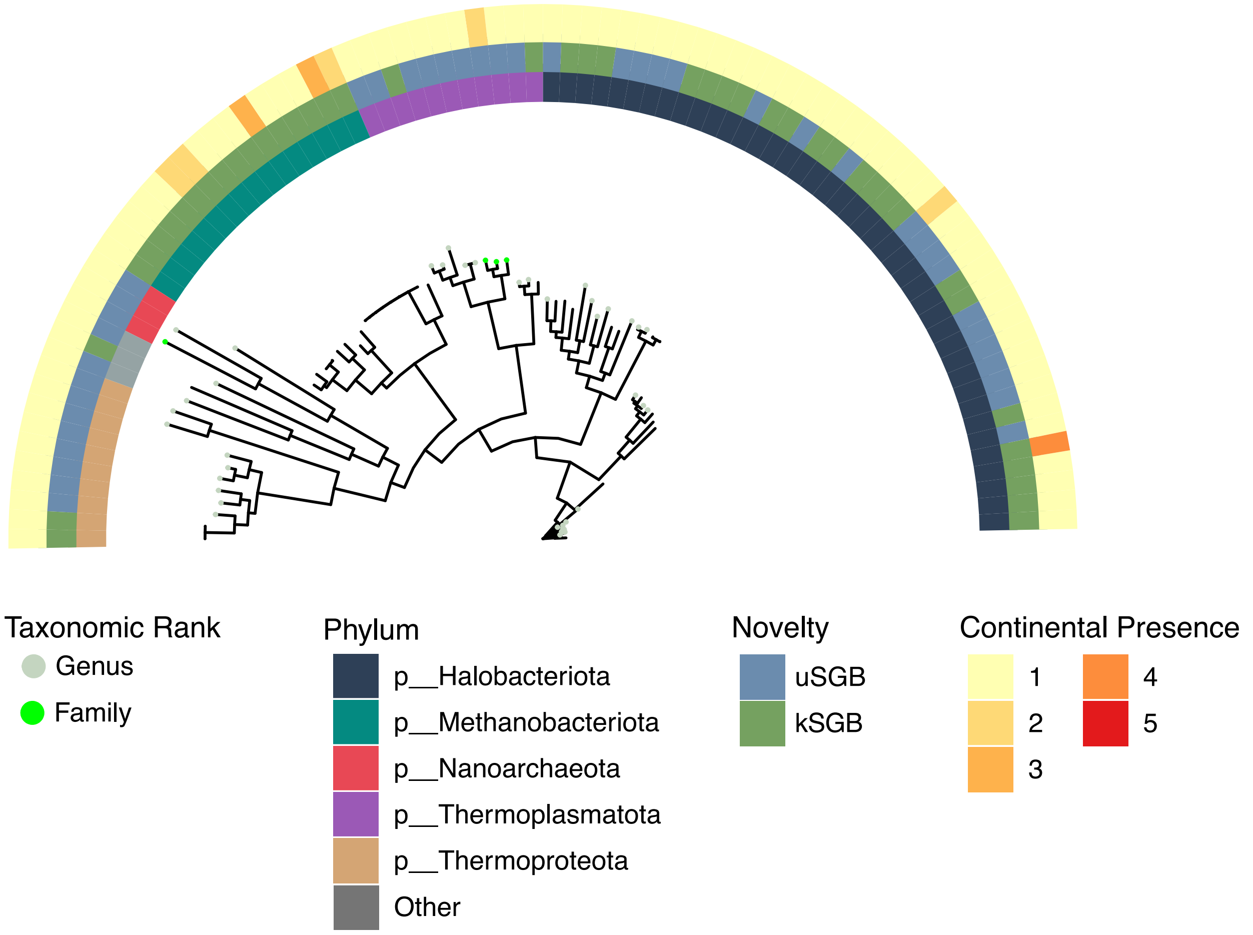

**Supplementary Figure 6:** Archaeal SGB phylogeny. Tip colors represent the assigned taxonomic rank of each SGB excluding species. The inner ring indicates the five most common phyla, the middle represents known or unknown status of the SGB, and the outer ring indicates on how many continents the species was present.

7)

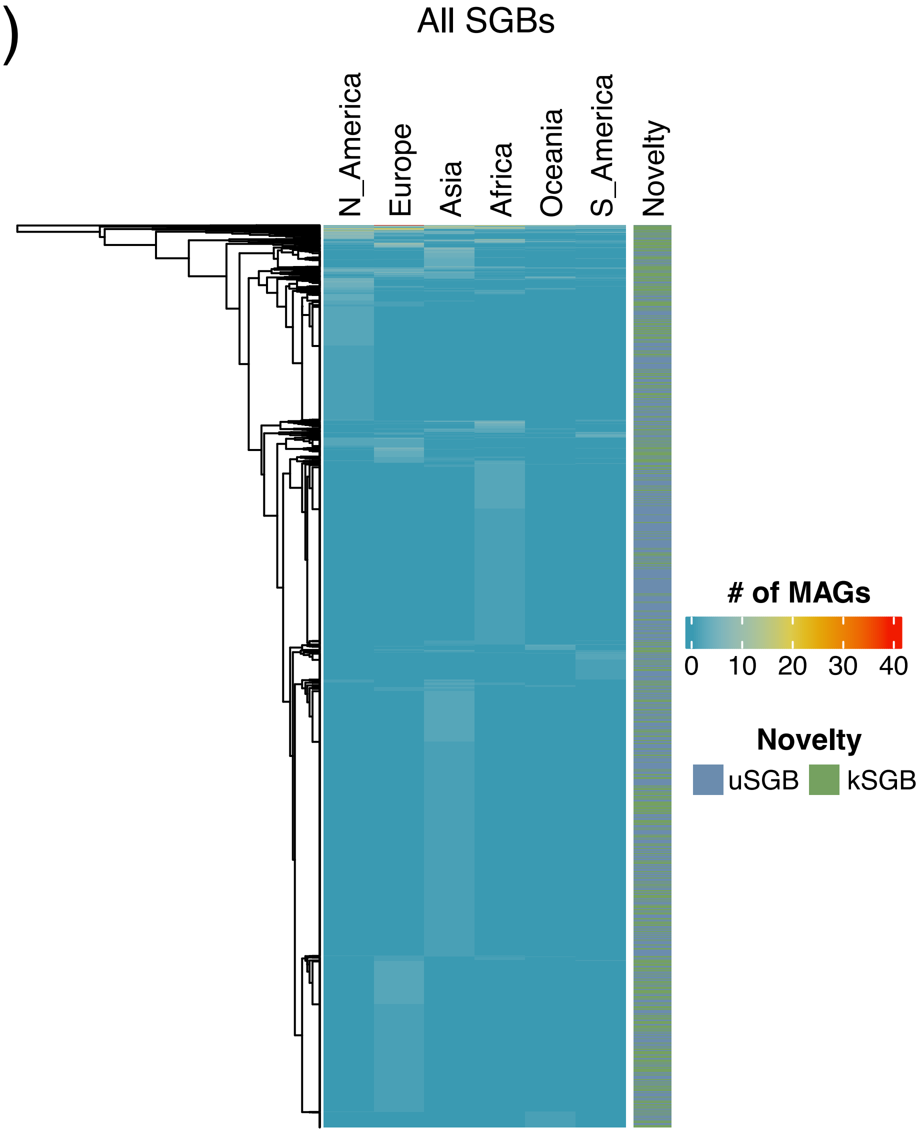

**Supplementary Figure 7:** Heatmap of all SGBs using continuous continental presence.

8)

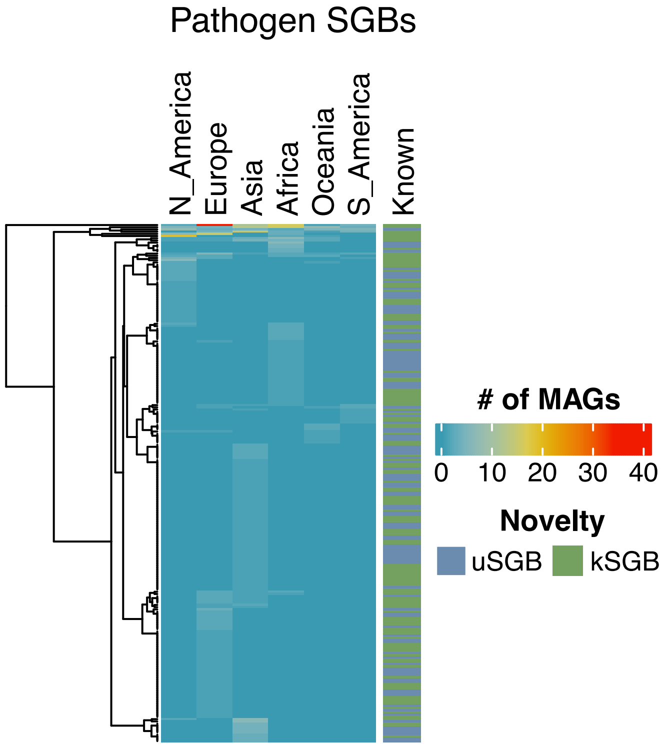

**Supplementary Figure 8:** Heatmap of all pathogenic SGBs using binary continental presence.

9)

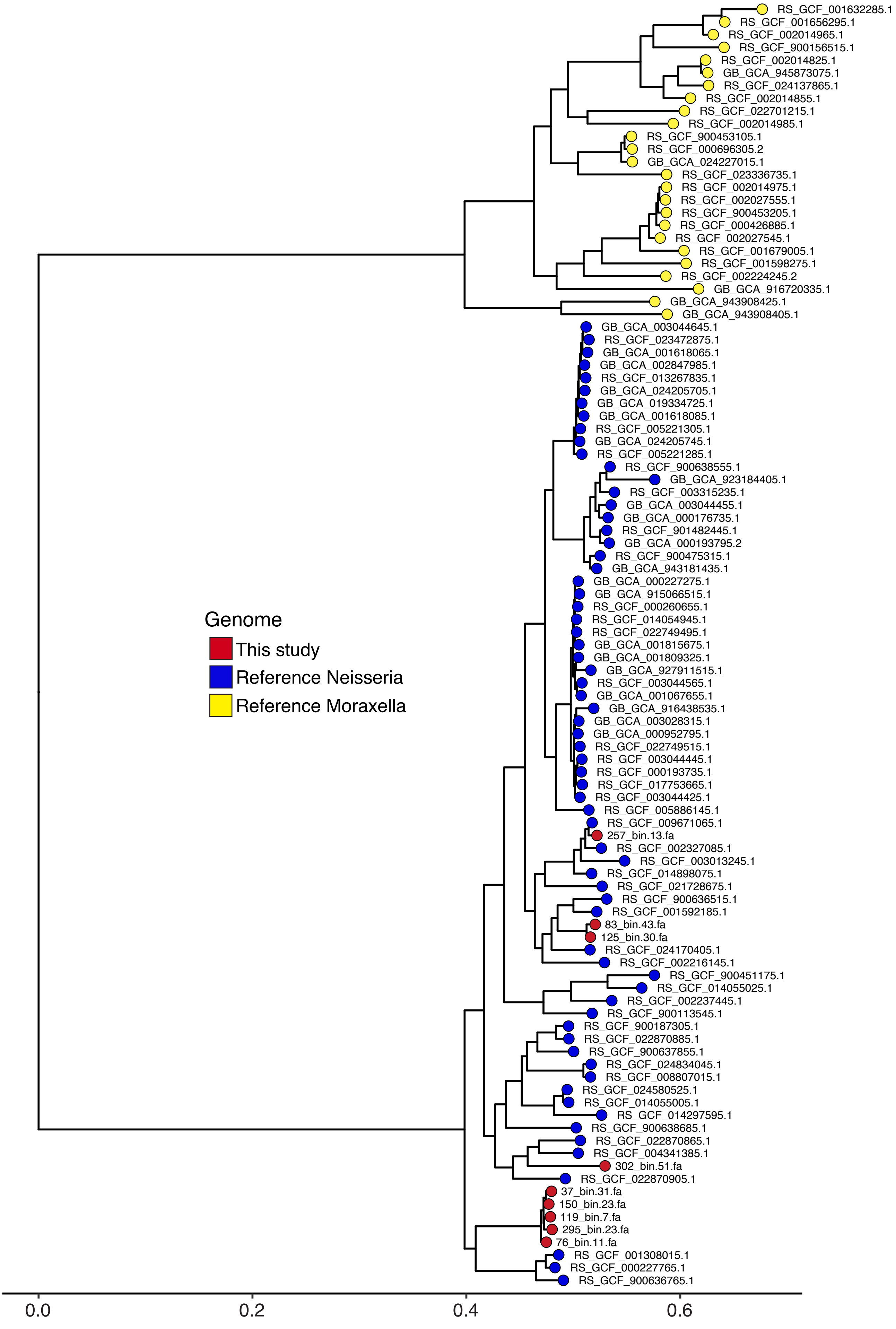

**Supplementary Figure 9:** Phylogeny of unknown Neisseria SGBs and reference Neisseria using Moraxella genomes as the outgroup.

**Supplementary Table 1: List of SRR runs used in the study along with corresponding metadata**

| name | bioproject | sra_run_acc | notes | location | assembling | Seq | subseq | country |
|---|---|---|---|---|---|---|---|---|
| 1 | PRJNA801794 | SRR17818322 | sewage | Milwaukee, V | co | Metagenome | MetaG | USA |
| 1 | PRJNA801794 | SRR17818323 | sewage | Milwaukee, V | co | Metagenome | MetaG | USA |
| 1 | PRJNA801794 | SRR17818325 | sewage | Milwaukee, V | co | Metagenome | MetaG | USA |
| 2 | PRJNA801794 | SRR17818328 | sewage | Palo Alto, CA | co | Metagenome | MetaG | USA |
| 2 | PRJNA801794 | SRR17818321 | sewage | Palo Alto, CA | co | Metagenome | MetaG | USA |
| 2 | PRJNA801794 | SRR17818316 | sewage | Palo Alto, CA | co | Metagenome | MetaG | USA |
| 3 | PRJNA801794 | SRR17818330 | sewage | Gloucester, N | co | Metagenome | MetaG | USA |
| 3 | PRJNA801794 | SRR17818327 | sewage | Gloucester, N | co | Metagenome | MetaG | USA |
| 3 | PRJNA801794 | SRR17818318 | sewage | Gloucester, N | co | Metagenome | MetaG | USA |
| 4 | PRJNA801794 | SRR17818317 | sewage | Key West, FL | co | Metagenome | MetaG | USA |
| 4 | PRJNA801794 | SRR17818319 | sewage | Key West, FL | co | Metagenome | MetaG | USA |
| 4 | PRJNA801794 | SRR17818314 | sewage | Key West, FL | co | Metagenome | MetaG | USA |
| 5 | PRJNA801794 | SRR17818329 | sewage | Laramie, WY | co | Metagenome | MetaG | USA |
| 5 | PRJNA801794 | SRR17818320 | sewage | Laramie, WY | co | Metagenome | MetaG | USA |
| 5 | PRJNA801794 | SRR17818315 | sewage | Laramie, WY | co | Metagenome | MetaG | USA |
| 6 | PRJNA801794 | SRR17818326 | sewage | Oak Creek, W | co | Metagenome | MetaG | USA |
| 6 | PRJNA801794 | SRR17818331 | sewage | Oak Creek, W | co | Metagenome | MetaG | USA |
| 6 | PRJNA801794 | SRR17818324 | sewage | Oak Creek, W | co | Metagenome | MetaG | USA |
| 7 | PRJNA774511 | SRR16642476 | ww from pig farm | China | single | Metagenome | MetaG | China |
| 8 | PRJNA774511 | SRR16642472 | ww from pig farm | China | single | Metagenome | MetaG | China |
| 9 | PRJNA774511 | SRR16642459 | ww from pig farm | China | single | Metagenome | MetaG | China |
| 10 | PRJNA774511 | SRR16642446 | ww from pig farm | China | single | Metagenome | MetaG | China |
| 11 | PRJNA774511 | SRR16642433 | ww from pig farm | China | single | Metagenome | MetaG | China |
| 12 | PRJNA774511 | SRR16642419 | ww from pig farm | China | single | Metagenome | MetaG | China |
| 13 | PRJNA70623 | SRR315458 | sewage | Pittsburgh, PA | single | Metagenome | MetaG | USA |
| 14 | PRJNA70623 | SRR315459 | sewage | Addis Ababa, | single | Metagenome | MetaG | Ethiopia |
| 15 | PRJNA70623 | SRR315457 | sewage viromes | Barcelona, Sp | single | Metagenome | MetaG | Spain |
| 22 | PRJNA300541 | SRR2938061 | sewage | El Salvador | co | Metagenome | MetaG | El Salvador |
| 22 | PRJNA300541 | SRR2938063 | sewage | El Salvador | co | Metagenome | MetaG | El Salvador |
| 22 | PRJNA300541 | SRR2938064 | sewage | El Salvador | co | Metagenome | MetaG | El Salvador |
| 22 | PRJNA300541 | SRR2938065 | sewage | El Salvador | co | Metagenome | MetaG | El Salvador |
| 22 | PRJNA300541 | SRR2938066 | sewage | El Salvador | co | Metagenome | MetaG | El Salvador |
| 22 | PRJNA300541 | SRR2938067 | sewage | El Salvador | co | Metagenome | MetaG | El Salvador |
| 22 | PRJNA300541 | SRR2938268 | sewage | El Salvador | co | Metagenome | MetaG | El Salvador |
| 22 | PRJNA300541 | SRR2938270 | sewage | El Salvador | co | Metagenome | MetaG | El Salvador |
| 22 | PRJNA300541 | SRR2938271 | sewage | El Salvador | co | Metagenome | MetaG | El Salvador |
| 22 | PRJNA300541 | SRR2938274 | sewage | El Salvador | co | Metagenome | MetaG | El Salvador |
| 22 | PRJNA300541 | SRR2938277 | sewage | El Salvador | co | Metagenome | MetaG | El Salvador |
| 22 | PRJNA300541 | SRR2938285 | sewage | El Salvador | co | Metagenome | MetaG | El Salvador |
| 22 | PRJNA300541 | SRR2938286 | sewage | El Salvador | co | Metagenome | MetaG | El Salvador |
| 22 | PRJNA300541 | SRR2938287 | sewage | El Salvador | co | Metagenome | MetaG | El Salvador |
| 22 | PRJNA300541 | SRR2938288 | sewage | El Salvador | co | Metagenome | MetaG | El Salvador |
| 22 | PRJNA300541 | SRR2938291 | sewage | El Salvador | co | Metagenome | MetaG | El Salvador |
| 22 | PRJNA300541 | SRR2938294 | sewage | El Salvador | co | Metagenome | MetaG | El Salvador |
| 22 | PRJNA300541 | SRR2938304 | sewage | El Salvador | co | Metagenome | MetaG | El Salvador |
| 22 | PRJNA300541 | SRR2938307 | sewage | El Salvador | co | Metagenome | MetaG | El Salvador |
| 23 | PRJNA300541 | SRR2938311 | sewage | Lima, Peru | co | Metagenome | MetaG | Peru |
| 23 | PRJNA300541 | SRR2938312 | sewage | Lima, Peru | co | Metagenome | MetaG | Peru |
| 23 | PRJNA300541 | SRR2938313 | sewage | Lima, Peru | co | Metagenome | MetaG | Peru |
| 23 | PRJNA300541 | SRR2938314 | sewage | Lima, Peru | co | Metagenome | MetaG | Peru |
| 23 | PRJNA300541 | SRR2938315 | sewage | Lima, Peru | co | Metagenome | MetaG | Peru |
| 23 | PRJNA300541 | SRR2938316 | sewage | Lima, Peru | co | Metagenome | MetaG | Peru |
| 23 | PRJNA300541 | SRR2938317 | sewage | Lima, Peru | co | Metagenome | MetaG | Peru |
| 23 | PRJNA300541 | SRR2938318 | sewage | Lima, Peru | co | Metagenome | MetaG | Peru |
| 23 | PRJNA300541 | SRR2938319 | sewage | Lima, Peru | co | Metagenome | MetaG | Peru |
| 23 | PRJNA300541 | SRR2938320 | sewage | Lima, Peru | co | Metagenome | MetaG | Peru |
| 23 | PRJNA300541 | SRR2938321 | sewage | Lima, Peru | co | Metagenome | MetaG | Peru |
| 23 | PRJNA300541 | SRR2938322 | sewage | Lima, Peru | co | Metagenome | MetaG | Peru |
| 23 | PRJNA300541 | SRR2938323 | sewage | Lima, Peru | co | Metagenome | MetaG | Peru |
| 23 | PRJNA300541 | SRR2938324 | sewage | Lima, Peru | co | Metagenome | MetaG | Peru |
| 23 | PRJNA300541 | SRR2938325 | sewage | Lima, Peru | co | Metagenome | MetaG | Peru |
| 23 | PRJNA300541 | SRR2938326 | sewage | Lima, Peru | co | Metagenome | MetaG | Peru |
| 23 | PRJNA300541 | SRR2938327 | sewage | Lima, Peru | co | Metagenome | MetaG | Peru |
| 23 | PRJNA300541 | SRR2938329 | sewage | Lima, Peru | co | Metagenome | MetaG | Peru |
| 23 | PRJNA300541 | SRR2938330 | sewage | Lima, Peru | co | Metagenome | MetaG | Peru |
| 23 | PRJNA300541 | SRR2938331 | sewage | Lima, Peru | co | Metagenome | MetaG | Peru |
| 23 | PRJNA300541 | SRR2938332 | sewage | Lima, Peru | co | Metagenome | MetaG | Peru |
| 23 | PRJNA300541 | SRR2938333 | sewage | Lima, Peru | co | Metagenome | MetaG | Peru |
| 23 | PRJNA300541 | SRR2938334 | sewage | Lima, Peru | co | Metagenome | MetaG | Peru |
| 23 | PRJNA300541 | SRR2938335 | sewage | Lima, Peru | co | Metagenome | MetaG | Peru |
| 23 | PRJNA300541 | SRR2938336 | sewage | Lima, Peru | co | Metagenome | MetaG | Peru |
| 23 | PRJNA300541 | SRR2938112 | sewage | Lima, Peru | co | Metagenome | MetaG | Peru |
| 23 | PRJNA300541 | SRR2938125 | sewage | Lima, Peru | co | Metagenome | MetaG | Peru |

| | | | | | | | | |
|---|---|---|---|---|---|---|---|---|
| 23 | PRJNA300541 | SRR2938126 | sewage | Lima, Peru | co | Metagenome | MetaG | Peru |
| 23 | PRJNA300541 | SRR2938127 | sewage | Lima, Peru | co | Metagenome | MetaG | Peru |
| 23 | PRJNA300541 | SRR2938128 | sewage | Lima, Peru | co | Metagenome | MetaG | Peru |
| 23 | PRJNA300541 | SRR2938129 | sewage | Lima, Peru | co | Metagenome | MetaG | Peru |
| 23 | PRJNA300541 | SRR2938130 | sewage | Lima, Peru | co | Metagenome | MetaG | Peru |
| 23 | PRJNA300541 | SRR2938131 | sewage | Lima, Peru | co | Metagenome | MetaG | Peru |
| 23 | PRJNA300541 | SRR2938132 | sewage | Lima, Peru | co | Metagenome | MetaG | Peru |
| 23 | PRJNA300541 | SRR2938232 | sewage | Lima, Peru | co | Metagenome | MetaG | Peru |
| 24 | PRJEB34690 | ERR3569137 | hospital ww | Pretoria, Sout | co | Metagenome | MetaG | South Africa |
| 24 | PRJEB34690 | ERR3569138 | hospital ww | Pretoria, Sout | co | Metagenome | MetaG | South Africa |
| 24 | PRJEB34690 | ERR3569139 | hospital ww | Pretoria, Sout | co | Metagenome | MetaG | South Africa |
| 25 | PRJEB34690 | ERR3569140 | hospital ww | Pretoria, Sout | co | Metagenome | MetaG | South Africa |
| 25 | PRJEB34690 | ERR3569141 | hospital ww | Pretoria, Sout | co | Metagenome | MetaG | South Africa |
| 25 | PRJEB34690 | ERR3569142 | hospital ww | Pretoria, Sout | co | Metagenome | MetaG | South Africa |
| 26 | PRJNA616359 | SRR11461807 | ww viral | USA | single | Metagenome | MetaG | USA |
| 27 | PRJNA616359 | SRR11461808 | ww viral | USA | single | Metagenome | MetaG | USA |
| 28 | PRJNA616359 | SRR11461806 | ww viral | USA | single | Metagenome | MetaG | USA |
| 29 | PRJNA616359 | SRR11461805 | ww viral | USA | single | Metagenome | MetaG | USA |
| 30 | PRJNA616359 | SRR11461803 | ww viral | USA | single | Metagenome | MetaG | USA |
| 31 | PRJNA616359 | SRR11461804 | ww viral | USA | single | Metagenome | MetaG | USA |
| 32 | PRJNA616359 | SRR11461810 | ww viral | USA | single | Metagenome | MetaG | USA |
| 33 | PRJNA616359 | SRR11461809 | ww viral | USA | single | Metagenome | MetaG | USA |
| 34 | PRJNA264280 | SRR1616983 | medical sewage | Germany, sev | co | Metagenome | MetaG | Germany |
| 34 | PRJNA264280 | SRR1616984 | medical sewage | Germany, sev | co | Metagenome | MetaG | Germany |
| 34 | PRJNA264280 | SRR1616985 | medical sewage | Germany, sev | co | Metagenome | MetaG | Germany |
| 35 | PRJNA264280 | SRR1616986 | medical sewage | Germany, sev | co | Metagenome | MetaG | Germany |
| 35 | PRJNA264280 | SRR1616987 | medical sewage | Germany, sev | co | Metagenome | MetaG | Germany |
| 35 | PRJNA264280 | SRR1616988 | medical sewage | Germany, sev | co | Metagenome | MetaG | Germany |
| 36 | PRJNA415974 | SRR6231144 | hospital ww | Sugar Creek, | co | Metagenome | MetaG | USA |
| 36 | PRJNA415974 | SRR6231176 | hospital ww | Sugar Creek, | co | Metagenome | MetaG | USA |
| 37 | PRJNA415974 | SRR6231209 | hospital ww | Mallard Creel | co | Metagenome | MetaG | USA |
| 37 | PRJNA415974 | SRR6231213 | hospital ww | Mallard Creel | co | Metagenome | MetaG | USA |
| 37 | PRJNA415974 | SRR6231145 | hospital ww | Mallard Creel | co | Metagenome | MetaG | USA |
| 37 | PRJNA415974 | SRR6231162 | hospital ww | Mallard Creel | co | Metagenome | MetaG | USA |
| 38 | PRJNA515946 | SRR8573788 | sewage | Santa Catalin | co | Metagenome | MetaG | Uraguay |
| 38 | PRJNA515946 | SRR8573789 | sewage | Santa Catalin | co | Metagenome | MetaG | Uraguay |
| 38 | PRJNA515946 | SRR8573790 | sewage | Santa Catalin | co | Metagenome | MetaG | Uraguay |
| 38 | PRJNA515946 | SRR8573791 | sewage | Santa Catalin | co | Metagenome | MetaG | Uraguay |
| 39 | PRJNA515946 | SRR8573793 | sewage | Zabala, Uragı | single | Metagenome | MetaG | Uraguay |
| 40 | PRJNA515946 | SRR8573796 | sewage | Montevideo, | single | Metagenome | MetaG | Uruguay |
| 41 | PRJNA515946 | SRR8573804 | sewage | Villa Arejo, U | single | Metagenome | MetaG | Uruguay |
| 42 | PRJNA515946 | SRR8573807 | sewage | Cerro, Urugua | single | Metagenome | MetaG | Uruguay |
| 43 | PRJNA824545 | SRR18715497 | wwtp influent | China | single | Metagenome | MetaG | China |
| 44 | PRJNA824545 | SRR18715500 | wwtp influent | China | single | Metagenome | MetaG | China |
| 45 | PRJNA824545 | SRR18715503 | wwtp influent | China | single | Metagenome | MetaG | China |
| 46 | PRJNA824545 | SRR18715513 | wwtp influent | China | single | Metagenome | MetaG | China |
| 47 | PRJNA693629 | SRR13612679 | wwtp influent | Johore, Malay | single | Metagenome | MetaG | Malaysia |
| 48 | PRJNA693629 | SRR13612675 | wwtp influent | Johore, Malay | single | Metagenome | MetaG | Malaysia |
| 49 | PRJNA693629 | SRR13612673 | wwtp influent | Johore, Malay | single | Metagenome | MetaG | Malaysia |
| 50 | PRJNA693629 | SRR13612671 | wwtp influent | Johore, Malay | single | Metagenome | MetaG | Malaysia |
| 51 | PRJNA693629 | SRR13612669 | wwtp influent | Johore, Malay | single | Metagenome | MetaG | Malaysia |
| 52 | PRJNA693629 | SRR13612677 | wwtp influent | Johore, Malay | single | Metagenome | MetaG | Malaysia |
| 53 | PRJNA612238 | SRR11296558 | hospital sewage | China | single | Metagenome | MetaG | China |
| 54 | PRJNA612238 | SRR11296557 | hospital sewage | China | single | Metagenome | MetaG | China |
| 55 | PRJNA612238 | SRR11296562 | hospital sewage | China | single | Metagenome | MetaG | China |
| 56 | PRJNA612238 | SRR11296561 | hospital sewage | China | single | Metagenome | MetaG | China |
| 57 | PRJNA612238 | SRR11296560 | hospital sewage | China | single | Metagenome | MetaG | China |
| 58 | PRJNA612238 | SRR11296559 | hospital sewage | China | single | Metagenome | MetaG | China |
| 59 | PRJEB34410 | ERR3519521 | hospital ww | Edinburgh, Sc | co | Metagenome | MetaG | Scotland |
| 59 | PRJEB34410 | ERR3519522 | hospital ww | Edinburgh, Sc | co | Metagenome | MetaG | Scotland |
| 59 | PRJEB34410 | ERR3519523 | hospital ww | Edinburgh, Sc | co | Metagenome | MetaG | Scotland |
| 59 | PRJEB34410 | ERR3519524 | hospital ww | Edinburgh, Sc | co | Metagenome | MetaG | Scotland |
| 59 | PRJEB34410 | ERR3519526 | hospital ww | Edinburgh, Sc | co | Metagenome | MetaG | Scotland |
| 59 | PRJEB34410 | ERR3519525 | hospital ww | Edinburgh, Sc | co | Metagenome | MetaG | Scotland |
| 59 | PRJEB34410 | ERR3519528 | hospital ww | Edinburgh, Sc | co | Metagenome | MetaG | Scotland |
| 59 | PRJEB34410 | ERR3519527 | hospital ww | Edinburgh, Sc | co | Metagenome | MetaG | Scotland |
| 60 | PRJEB23496 | ERR2596700 | sewage viromes | Atlanta, GA, I | single | Metavirome | MetaV: WGS | USA |
| 61 | PRJEB23496 | ERR2596699 | sewage viromes | Belgrade, Ser | single | Metavirome | MetaV: WGS | Serbia |
| 62 | PRJEB23496 | ERR2596698 | sewage viromes | Riga, Latvia | single | Metavirome | MetaV: WGS | Latvia |
| 63 | PRJEB23496 | ERR2596697 | sewage viromes | Helsinki, Finla | single | Metavirome | MetaV: WGS | Finland |
| 64 | PRJEB23496 | ERR2596696 | sewage viromes | Toronto, Cana | single | Metavirome | MetaV: WGS | Canada |
| 65 | PRJEB23496 | ERR2596695 | sewage viromes | N'Djamena, ( | single | Metavirome | MetaV: WGS | Chad |
| 66 | PRJEB13832 | ERR1525434 | sewage | Copenhagen, | co | Metagenome | MetaG | Denmark |
| 66 | PRJEB13832 | ERR1525435 | sewage | Copenhagen, | co | Metagenome | MetaG | Denmark |
| 66 | PRJEB13832 | ERR1525437 | sewage | Copenhagen, | co | Metagenome | MetaG | Denmark |

| | | | | | | | | |
|---|---|---|---|---|---|---|---|---|
| 66 | PRJEB13832 | ERR1525436 | sewage | Copenhagen, | co | Metagenome | MetaG | Denmark |
| 66 | PRJEB13832 | ERR1525446 | sewage | Copenhagen, | co | Metagenome | MetaG | Denmark |
| 66 | PRJEB13832 | ERR1525440 | sewage | Copenhagen, | co | Metagenome | MetaG | Denmark |
| 66 | PRJEB13832 | ERR1525441 | sewage | Copenhagen, | co | Metagenome | MetaG | Denmark |
| 66 | PRJEB13832 | ERR1525439 | sewage | Copenhagen, | co | Metagenome | MetaG | Denmark |
| 66 | PRJEB13832 | ERR1525445 | sewage | Copenhagen, | co | Metagenome | MetaG | Denmark |
| 66 | PRJEB13832 | ERR1525444 | sewage | Copenhagen, | co | Metagenome | MetaG | Denmark |
| 66 | PRJEB13832 | ERR1525438 | sewage | Copenhagen, | co | Metagenome | MetaG | Denmark |
| 66 | PRJEB13832 | ERR1525443 | sewage | Copenhagen, | co | Metagenome | MetaG | Denmark |
| 66 | PRJEB13832 | ERR1525442 | sewage | Copenhagen, | co | Metagenome | MetaG | Denmark |
| 66 | PRJEB13832 | ERR1512995 | sewage | Copenhagen, | co | Metagenome | MetaG | Denmark |
| 66 | PRJEB13832 | ERR1512996 | sewage | Copenhagen, | co | Metagenome | MetaG | Denmark |
| 66 | PRJEB13832 | ERR1512994 | sewage | Copenhagen, | co | Metagenome | MetaG | Denmark |
| 66 | PRJEB13832 | ERR1512993 | sewage | Copenhagen, | co | Metagenome | MetaG | Denmark |
| 66 | PRJEB13832 | ERR1513007 | sewage | Copenhagen, | co | Metagenome | MetaG | Denmark |
| 66 | PRJEB13832 | ERR1513003 | sewage | Copenhagen, | co | Metagenome | MetaG | Denmark |
| 66 | PRJEB13832 | ERR1512997 | sewage | Copenhagen, | co | Metagenome | MetaG | Denmark |
| 66 | PRJEB13832 | ERR1512991 | sewage | Copenhagen, | co | Metagenome | MetaG | Denmark |
| 66 | PRJEB13832 | ERR1512992 | sewage | Copenhagen, | co | Metagenome | MetaG | Denmark |
| 66 | PRJEB13832 | ERR1512998 | sewage | Copenhagen, | co | Metagenome | MetaG | Denmark |
| 66 | PRJEB13832 | ERR1513004 | sewage | Copenhagen, | co | Metagenome | MetaG | Denmark |
| 66 | PRJEB13832 | ERR1513000 | sewage | Copenhagen, | co | Metagenome | MetaG | Denmark |
| 66 | PRJEB13832 | ERR1513006 | sewage | Copenhagen, | co | Metagenome | MetaG | Denmark |
| 66 | PRJEB13832 | ERR1513005 | sewage | Copenhagen, | co | Metagenome | MetaG | Denmark |
| 66 | PRJEB13832 | ERR1512999 | sewage | Copenhagen, | co | Metagenome | MetaG | Denmark |
| 66 | PRJEB13832 | ERR1512990 | sewage | Copenhagen, | co | Metagenome | MetaG | Denmark |
| 66 | PRJEB13832 | ERR1512989 | sewage | Copenhagen, | co | Metagenome | MetaG | Denmark |
| 66 | PRJEB13832 | ERR1513001 | sewage | Copenhagen, | co | Metagenome | MetaG | Denmark |
| 66 | PRJEB13832 | ERR1512987 | sewage | Copenhagen, | co | Metagenome | MetaG | Denmark |
| 66 | PRJEB13832 | ERR1512988 | sewage | Copenhagen, | co | Metagenome | MetaG | Denmark |
| 66 | PRJEB13832 | ERR1513002 | sewage | Copenhagen, | co | Metagenome | MetaG | Denmark |
| 66 | PRJEB13832 | ERR1512984 | sewage | Copenhagen, | co | Metagenome | MetaG | Denmark |
| 66 | PRJEB13832 | ERR1512985 | sewage | Copenhagen, | co | Metagenome | MetaG | Denmark |
| 66 | PRJEB13832 | ERR1512986 | sewage | Copenhagen, | co | Metagenome | MetaG | Denmark |
| 66 | PRJEB13832 | ERR1514429 | sewage | Copenhagen, | co | Metagenome | MetaG | Denmark |
| 66 | PRJEB13832 | ERR1514430 | sewage | Copenhagen, | co | Metagenome | MetaG | Denmark |
| 66 | PRJEB13832 | ERR1514428 | sewage | Copenhagen, | co | Metagenome | MetaG | Denmark |
| 66 | PRJEB13832 | ERR1514427 | sewage | Copenhagen, | co | Metagenome | MetaG | Denmark |
| 83 | PRJEB13831 | ERR1725965 | sewage | Abidjan, Cote | co | Metagenome | MetaG | Cote d'Ivoire |
| 83 | PRJEB13831 | ERR1713345 | sewage | Abidjan, Cote | co | Metagenome | MetaG | Cote d'Ivoire |
| 83 | PRJEB13831 | ERR2592252 | sewage | Abidjan, Cote | co | Metagenome | MetaG | Cote d'Ivoire |
| 84 | PRJEB13831 | ERR1713355 | sewage | Addis Ababa, | single | Metagenome | MetaG | Ethiopia |
| 85 | PRJEB13831 | ERR1713368 | sewage | Almaty, Kaza | single | Metagenome | MetaG | Kazakhstan |
| 86 | PRJEB13831 | ERR1726009 | sewage | Ankara, Turke | co | Metagenome | MetaG | Turkey |
| 86 | PRJEB13831 | ERR1713395 | sewage | Ankara, Turke | co | Metagenome | MetaG | Turkey |
| 86 | PRJEB13831 | ERR2592273 | sewage | Ankara, Turke | co | Metagenome | MetaG | Turkey |
| 87 | PRJEB13831 | ERR1713397 | sewage | Atlanta, GA, I | single | Metagenome | MetaG | USA |
| 88 | PRJEB13831 | ERR1713359 | sewage | Banjul, Gamb | single | Metagenome | MetaG | Gambia |
| 89 | PRJEB13831 | ERR1725972 | sewage | Barcelona, Sp | co | Metagenome | MetaG | Spain |
| 89 | PRJEB13831 | ERR1725971 | sewage | Barcelona, Sp | co | Metagenome | MetaG | Spain |
| 89 | PRJEB13831 | ERR1725970 | sewage | Barcelona, Sp | co | Metagenome | MetaG | Spain |
| 89 | PRJEB13831 | ERR1725969 | sewage | Barcelona, Sp | co | Metagenome | MetaG | Spain |
| 89 | PRJEB13831 | ERR1713354 | sewage | Barcelona, Sp | co | Metagenome | MetaG | Spain |
| 89 | PRJEB13831 | ERR2592334 | sewage | Barcelona, Sp | co | Metagenome | MetaG | Spain |
| 90 | PRJEB13831 | ERR1725995 | sewage | Bedong, Mala | co | Metagenome | MetaG | Malaysia |
| 90 | PRJEB13831 | ERR1725994 | sewage | Bedong, Mala | co | Metagenome | MetaG | Malaysia |
| 90 | PRJEB13831 | ERR1713377 | sewage | Bedong, Mala | co | Metagenome | MetaG | Malaysia |
| 90 | PRJEB13831 | ERR2592337 | sewage | Bedong, Mala | co | Metagenome | MetaG | Malaysia |
| 90 | PRJEB13831 | ERR2592264 | sewage | Bedong, Mala | co | Metagenome | MetaG | Malaysia |
| 91 | PRJEB13831 | ERR1725951 | sewage | Belem, Brasil | co | Metagenome | MetaG | Brasil |
| 91 | PRJEB13831 | ERR1725950 | sewage | Belem, Brasil | co | Metagenome | MetaG | Brasil |
| 91 | PRJEB13831 | ERR1725949 | sewage | Belem, Brasil | co | Metagenome | MetaG | Brasil |
| 91 | PRJEB13831 | ERR1713337 | sewage | Belem, Brasil | co | Metagenome | MetaG | Brasil |
| 91 | PRJEB13831 | ERR1713336 | sewage | Belem, Brasil | co | Metagenome | MetaG | Brasil |
| 91 | PRJEB13831 | ERR2592248 | sewage | Belem, Brasil | co | Metagenome | MetaG | Brasil |
| 92 | PRJEB13831 | ERR1726002 | sewage | Belgrade, Ser | co | Metagenome | MetaG | Serbia |
| 92 | PRJEB13831 | ERR1713388 | sewage | Belgrade, Ser | co | Metagenome | MetaG | Serbia |
| 92 | PRJEB13831 | ERR2592270 | sewage | Belgrade, Ser | co | Metagenome | MetaG | Serbia |
| 93 | PRJEB13831 | ERR1713372 | sewage | Belvaux, Luxe | single | Metagenome | MetaG | Luxembourg |
| 94 | PRJEB13831 | ERR1725968 | sewage | Berlin, Germa | co | Metagenome | MetaG | Germany |
| 94 | PRJEB13831 | ERR1725967 | sewage | Berlin, Germa | co | Metagenome | MetaG | Germany |
| 94 | PRJEB13831 | ERR1713348 | sewage | Berlin, Germa | co | Metagenome | MetaG | Germany |
| 94 | PRJEB13831 | ERR2592332 | sewage | Berlin, Germa | co | Metagenome | MetaG | Germany |
| 94 | PRJEB13831 | ERR2592254 | sewage | Berlin, Germa | co | Metagenome | MetaG | Germany |
| 95 | PRJEB13831 | ERR1725998 | sewage | Bilthoven, Th | co | Metagenome | MetaG | The Netherlands |

| | | | | | | | | |
|---|---|---|---|---|---|---|---|---|
| 95 | PRJEB13831 | ERR1725997 | sewage | Bilthoven, The | co | Metagenome | MetaG | The Netherlands |
| 95 | PRJEB13831 | ERR1713379 | sewage | Bilthoven, The | co | Metagenome | MetaG | The Netherlands |
| 95 | PRJEB13831 | ERR2592338 | sewage | Bilthoven, The | co | Metagenome | MetaG | The Netherlands |
| 95 | PRJEB13831 | ERR2592266 | sewage | Bilthoven, The | co | Metagenome | MetaG | The Netherlands |
| 96 | PRJEB13831 | ERR1713406 | sewage | Boulder, CO, | single | Metagenome | MetaG | USA |
| 97 | PRJEB13831 | ERR1726003 | sewage | Bratislava, Sl | co | Metagenome | MetaG | Slovakia |
| 97 | PRJEB13831 | ERR1713389 | sewage | Bratislava, Sl | co | Metagenome | MetaG | Slovakia |
| 98 | PRJEB13831 | ERR1725966 | sewage | Brno, Czech R | co | Metagenome | MetaG | Czech Republic |
| 98 | PRJEB13831 | ERR1713347 | sewage | Brno, Czech R | co | Metagenome | MetaG | Czech Republic |
| 98 | PRJEB13831 | ERR2592253 | sewage | Brno, Czech R | co | Metagenome | MetaG | Czech Republic |
| 99 | PRJEB13831 | ERR1713361 | sewage | Budapest, Hu | single | Metagenome | MetaG | Hungary |
| 100 | PRJEB13831 | ERR1725958 | sewage | Calgary, Cana | co | Metagenome | MetaG | Canada |
| 100 | PRJEB13831 | ERR1725957 | sewage | Calgary, Cana | co | Metagenome | MetaG | Canada |
| 100 | PRJEB13831 | ERR1725956 | sewage | Calgary, Cana | co | Metagenome | MetaG | Canada |
| 100 | PRJEB13831 | ERR1725955 | sewage | Calgary, Cana | co | Metagenome | MetaG | Canada |
| 100 | PRJEB13831 | ERR1713340 | sewage | Calgary, Cana | co | Metagenome | MetaG | Canada |
| 101 | PRJEB13831 | ERR1726015 | sewage | Chicago, IL, U | co | Metagenome | MetaG | USA |
| 101 | PRJEB13831 | ERR1713399 | sewage | Chicago, IL, U | co | Metagenome | MetaG | USA |
| 101 | PRJEB13831 | ERR2592340 | sewage | Chicago, IL, U | co | Metagenome | MetaG | USA |
| 101 | PRJEB13831 | ERR2592275 | sewage | Chicago, IL, U | co | Metagenome | MetaG | USA |
| 101 | PRJEB13831 | ERR1726014 | sewage | Chicago, IL, U | co | Metagenome | MetaG | USA |
| 101 | PRJEB13831 | ERR1726013 | sewage | Chicago, IL, U | co | Metagenome | MetaG | USA |
| 101 | PRJEB13831 | ERR1726012 | sewage | Chicago, IL, U | co | Metagenome | MetaG | USA |
| 102 | PRJEB13831 | ERR1725989 | sewage | Chisinau, Mol | co | Metagenome | MetaG | Moldova |
| 102 | PRJEB13831 | ERR1725988 | sewage | Chisinau, Mol | co | Metagenome | MetaG | Moldova |
| 102 | PRJEB13831 | ERR1713374 | sewage | Chisinau, Mol | co | Metagenome | MetaG | Moldova |
| 103 | PRJEB13831 | ERR1725978 | sewage | Cochin, India | co | Metagenome | MetaG | India |
| 103 | PRJEB13831 | ERR1713362 | sewage | Cochin, India | co | Metagenome | MetaG | India |
| 103 | PRJEB13831 | ERR2592258 | sewage | Cochin, India | co | Metagenome | MetaG | India |
| 104 | PRJEB13831 | ERR1713371 | sewage | Colombo, Sri | single | Metagenome | MetaG | Sri Lanka |
| 105 | PRJEB13831 | ERR1713351 | sewage | Copenhagen, | co | Metagenome | MetaG | Denmark |
| 105 | PRJEB13831 | ERR1713350 | sewage | Copenhagen, | co | Metagenome | MetaG | Denmark |
| 105 | PRJEB13831 | ERR1713349 | sewage | Copenhagen, | co | Metagenome | MetaG | Denmark |
| 106 | PRJEB13831 | ERR1713386 | sewage | Dakar, Seneg | single | Metagenome | MetaG | Senegal |
| 107 | PRJEB13831 | ERR1713382 | sewage | Dunedin, Nev | single | Metagenome | MetaG | New Zealand |
| 108 | PRJEB13831 | ERR1726029 | sewage | El Paso, TX, U | co | Metagenome | MetaG | USA |
| 108 | PRJEB13831 | ERR1713404 | sewage | El Paso, TX, U | co | Metagenome | MetaG | USA |
| 108 | PRJEB13831 | ERR1713403 | sewage | El Paso, TX, U | co | Metagenome | MetaG | USA |
| 108 | PRJEB13831 | ERR1713402 | sewage | El Paso, TX, U | co | Metagenome | MetaG | USA |
| 108 | PRJEB13831 | ERR1713400 | sewage | El Paso, TX, U | co | Metagenome | MetaG | USA |
| 108 | PRJEB13831 | ERR2592342 | sewage | El Paso, TX, U | co | Metagenome | MetaG | USA |
| 108 | PRJEB13831 | ERR2592341 | sewage | El Paso, TX, U | co | Metagenome | MetaG | USA |
| 108 | PRJEB13831 | ERR2592280 | sewage | El Paso, TX, U | co | Metagenome | MetaG | USA |
| 108 | PRJEB13831 | ERR2592279 | sewage | El Paso, TX, U | co | Metagenome | MetaG | USA |
| 108 | PRJEB13831 | ERR2592278 | sewage | El Paso, TX, U | co | Metagenome | MetaG | USA |
| 108 | PRJEB13831 | ERR1726028 | sewage | El Paso, TX, U | co | Metagenome | MetaG | USA |
| 108 | PRJEB13831 | ERR1726027 | sewage | El Paso, TX, U | co | Metagenome | MetaG | USA |
| 108 | PRJEB13831 | ERR1726026 | sewage | El Paso, TX, U | co | Metagenome | MetaG | USA |
| 108 | PRJEB13831 | ERR1726025 | sewage | El Paso, TX, U | co | Metagenome | MetaG | USA |
| 108 | PRJEB13831 | ERR1726024 | sewage | El Paso, TX, U | co | Metagenome | MetaG | USA |
| 108 | PRJEB13831 | ERR1726023 | sewage | El Paso, TX, U | co | Metagenome | MetaG | USA |
| 108 | PRJEB13831 | ERR1726022 | sewage | El Paso, TX, U | co | Metagenome | MetaG | USA |
| 108 | PRJEB13831 | ERR1726021 | sewage | El Paso, TX, U | co | Metagenome | MetaG | USA |
| 108 | PRJEB13831 | ERR1726020 | sewage | El Paso, TX, U | co | Metagenome | MetaG | USA |
| 108 | PRJEB13831 | ERR1726019 | sewage | El Paso, TX, U | co | Metagenome | MetaG | USA |
| 108 | PRJEB13831 | ERR1726018 | sewage | El Paso, TX, U | co | Metagenome | MetaG | USA |
| 108 | PRJEB13831 | ERR2592276 | sewage | El Paso, TX, U | co | Metagenome | MetaG | USA |
| 109 | PRJEB13831 | ERR1725980 | sewage | Galway, Irela | co | Metagenome | MetaG | Ireland |
| 109 | PRJEB13831 | ERR1725979 | sewage | Galway, Irela | co | Metagenome | MetaG | Ireland |
| 109 | PRJEB13831 | ERR1713363 | sewage | Galway, Irela | co | Metagenome | MetaG | Ireland |
| 110 | PRJEB13831 | ERR1726006 | sewage | Gothenburg, : | co | Metagenome | MetaG | Sweden |
| 110 | PRJEB13831 | ERR1726005 | sewage | Gothenburg, : | co | Metagenome | MetaG | Sweden |
| 110 | PRJEB13831 | ERR1713392 | sewage | Gothenburg, : | co | Metagenome | MetaG | Sweden |
| 111 | PRJEB13831 | ERR1713344 | sewage | Guangzhou, C | single | Metagenome | MetaG | China |
| 112 | PRJEB13831 | ERR1725964 | sewage | Herrenschwa | co | Metagenome | MetaG | Switzerland |
| 112 | PRJEB13831 | ERR1725963 | sewage | Herrenschwa | co | Metagenome | MetaG | Switzerland |
| 112 | PRJEB13831 | ERR1725962 | sewage | Herrenschwa | co | Metagenome | MetaG | Switzerland |
| 112 | PRJEB13831 | ERR1725961 | sewage | Herrenschwa | co | Metagenome | MetaG | Switzerland |
| 112 | PRJEB13831 | ERR1713343 | sewage | Herrenschwa | co | Metagenome | MetaG | Switzerland |
| 112 | PRJEB13831 | ERR2592331 | sewage | Herrenschwa | co | Metagenome | MetaG | Switzerland |
| 112 | PRJEB13831 | ERR2592251 | sewage | Herrenschwa | co | Metagenome | MetaG | Switzerland |
| 113 | PRJEB13831 | ERR1713407 | sewage | Ho Chi Minh, | single | Metagenome | MetaG | Vietnam |
| 114 | PRJEB13831 | ERR1725996 | sewage | Ibadan, Niger | co | Metagenome | MetaG | Nigeria |
| 114 | PRJEB13831 | ERR1713378 | sewage | Ibadan, Niger | co | Metagenome | MetaG | Nigeria |
| 114 | PRJEB13831 | ERR2592265 | sewage | Ibadan, Niger | co | Metagenome | MetaG | Nigeria |

| | | | | | | | | |
|---|---|---|---|---|---|---|---|---|
| 115 | PRJEB13831 | ERR1725984 | sewage | Jerusalem, Is | co | Metagenome | MetaG | Israel |
| 115 | PRJEB13831 | ERR1725983 | sewage | Jerusalem, Is | co | Metagenome | MetaG | Israel |
| 115 | PRJEB13831 | ERR1713366 | sewage | Jerusalem, Is | co | Metagenome | MetaG | Israel |
| 116 | PRJEB13831 | ERR1726030 | sewage | Johannesburg | co | Metagenome | MetaG | South Africa |
| 116 | PRJEB13831 | ERR1713409 | sewage | Johannesburg | co | Metagenome | MetaG | South Africa |
| 116 | PRJEB13831 | ERR2592281 | sewage | Johannesburg | co | Metagenome | MetaG | South Africa |
| 117 | PRJEB13831 | ERR1713384 | sewage | Karachi, Pakis | single | Metagenome | MetaG | Pakistan |
| 118 | PRJEB13831 | ERR1713381 | sewage | Kathmandu, I | single | Metagenome | MetaG | Nepal |
| 119 | PRJEB13831 | ERR1726035 | sewage | Kitwe, Zambi | co | Metagenome | MetaG | Zambia |
| 119 | PRJEB13831 | ERR1713411 | sewage | Kitwe, Zambi | co | Metagenome | MetaG | Zambia |
| 119 | PRJEB13831 | ERR2592343 | sewage | Kitwe, Zambi | co | Metagenome | MetaG | Zambia |
| 119 | PRJEB13831 | ERR2592283 | sewage | Kitwe, Zambi | co | Metagenome | MetaG | Zambia |
| 119 | PRJEB13831 | ERR1726034 | sewage | Kitwe, Zambi | co | Metagenome | MetaG | Zambia |
| 119 | PRJEB13831 | ERR1726033 | sewage | Kitwe, Zambi | co | Metagenome | MetaG | Zambia |
| 119 | PRJEB13831 | ERR1726032 | sewage | Kitwe, Zambi | co | Metagenome | MetaG | Zambia |
| 120 | PRJEB13831 | ERR1713385 | sewage | Lima, Peru | single | Metagenome | MetaG | Peru |
| 121 | PRJEB13831 | ERR1726004 | sewage | Ljubljana, Slo | co | Metagenome | MetaG | Slovenia |
| 121 | PRJEB13831 | ERR1713390 | sewage | Ljubljana, Slo | co | Metagenome | MetaG | Slovenia |
| 121 | PRJEB13831 | ERR2592271 | sewage | Ljubljana, Slo | co | Metagenome | MetaG | Slovenia |
| 122 | PRJEB13831 | ERR1713394 | sewage | Lome, Togo | single | Metagenome | MetaG | Togo |
| 123 | PRJEB13831 | ERR1726031 | sewage | Lusaka, Zamb | co | Metagenome | MetaG | Zambia |
| 123 | PRJEB13831 | ERR1713410 | sewage | Lusaka, Zamb | co | Metagenome | MetaG | Zambia |
| 123 | PRJEB13831 | ERR2592282 | sewage | Lusaka, Zamb | co | Metagenome | MetaG | Zambia |
| 124 | PRJEB13831 | ERR1725944 | sewage | Melbourne, A | co | Metagenome | MetaG | Australia |
| 124 | PRJEB13831 | ERR1725943 | sewage | Melbourne, A | co | Metagenome | MetaG | Australia |
| 124 | PRJEB13831 | ERR1725942 | sewage | Melbourne, A | co | Metagenome | MetaG | Australia |
| 124 | PRJEB13831 | ERR1725941 | sewage | Melbourne, A | co | Metagenome | MetaG | Australia |
| 124 | PRJEB13831 | ERR1713333 | sewage | Melbourne, A | co | Metagenome | MetaG | Australia |
| 125 | PRJEB13831 | ERR1713396 | sewage | Moshi Urban, | single | Metagenome | MetaG | Tanzania |
| 126 | PRJEB13831 | ERR1713346 | sewage | Mosquera, Co | single | Metagenome | MetaG | Colombia |
| 127 | PRJEB13831 | ERR1726008 | sewage | N'Djamena, C | co | Metagenome | MetaG | Chad |
| 127 | PRJEB13831 | ERR1726007 | sewage | N'Djamena, C | co | Metagenome | MetaG | Chad |
| 127 | PRJEB13831 | ERR1713393 | sewage | N'Djamena, C | co | Metagenome | MetaG | Chad |
| 127 | PRJEB13831 | ERR2592339 | sewage | N'Djamena, C | co | Metagenome | MetaG | Chad |
| 127 | PRJEB13831 | ERR2592272 | sewage | N'Djamena, C | co | Metagenome | MetaG | Chad |
| 128 | PRJEB13831 | ERR1725986 | sewage | Nairobi, Keny | co | Metagenome | MetaG | Kenya |
| 128 | PRJEB13831 | ERR1725985 | sewage | Nairobi, Keny | co | Metagenome | MetaG | Kenya |
| 128 | PRJEB13831 | ERR1713369 | sewage | Nairobi, Keny | co | Metagenome | MetaG | Kenya |
| 128 | PRJEB13831 | ERR2592261 | sewage | Nairobi, Keny | co | Metagenome | MetaG | Kenya |
| 129 | PRJEB13831 | ERR1725999 | sewage | Oslo, Norway | co | Metagenome | MetaG | Norway |
| 129 | PRJEB13831 | ERR1713380 | sewage | Oslo, Norway | co | Metagenome | MetaG | Norway |
| 129 | PRJEB13831 | ERR2592267 | sewage | Oslo, Norway | co | Metagenome | MetaG | Norway |
| 130 | PRJEB13831 | ERR1713342 | sewage | Ottawa, Cana | single | Metagenome | MetaG | Canada |
| 131 | PRJEB13831 | ERR1725953 | sewage | Palapye, Bots | co | Metagenome | MetaG | Botswana |
| 131 | PRJEB13831 | ERR1725952 | sewage | Palapye, Bots | co | Metagenome | MetaG | Botswana |
| 131 | PRJEB13831 | ERR1713338 | sewage | Palapye, Bots | co | Metagenome | MetaG | Botswana |
| 131 | PRJEB13831 | ERR2592249 | sewage | Palapye, Bots | co | Metagenome | MetaG | Botswana |
| 132 | PRJEB13831 | ERR1713370 | sewage | Phnom Penh, | single | Metagenome | MetaG | Cambodia |
| 133 | PRJEB13831 | ERR1726017 | sewage | Portland, OR, | co | Metagenome | MetaG | USA |
| 133 | PRJEB13831 | ERR1713401 | sewage | Portland, OR, | co | Metagenome | MetaG | USA |
| 133 | PRJEB13831 | ERR2592277 | sewage | Portland, OR, | co | Metagenome | MetaG | USA |
| 133 | PRJEB13831 | ERR1726016 | sewage | Portland, OR, | co | Metagenome | MetaG | USA |
| 134 | PRJEB13831 | ERR1713408 | sewage | Pristina, Koso | single | Metagenome | MetaG | Kosovo |
| 135 | PRJEB13831 | ERR1726000 | sewage | Pulawy, Polar | co | Metagenome | MetaG | Poland |
| 135 | PRJEB13831 | ERR1713383 | sewage | Pulawy, Polar | co | Metagenome | MetaG | Poland |
| 135 | PRJEB13831 | ERR2592268 | sewage | Pulawy, Polar | co | Metagenome | MetaG | Poland |
| 136 | PRJEB13831 | ERR1713352 | sewage | Quito, Ecuad | single | Metagenome | MetaG | Ecuador |
| 137 | PRJEB13831 | ERR1725954 | sewage | Regina, Cana | co | Metagenome | MetaG | Canada |
| 137 | PRJEB13831 | ERR1713339 | sewage | Regina, Cana | co | Metagenome | MetaG | Canada |
| 137 | PRJEB13831 | ERR2592329 | sewage | Regina, Cana | co | Metagenome | MetaG | Canada |
| 138 | PRJEB13831 | ERR1725982 | sewage | Reykjavik, Ice | co | Metagenome | MetaG | Iceland |
| 138 | PRJEB13831 | ERR1713365 | sewage | Reykjavik, Ice | co | Metagenome | MetaG | Iceland |
| 138 | PRJEB13831 | ERR2592260 | sewage | Reykjavik, Ice | co | Metagenome | MetaG | Iceland |
| 139 | PRJEB13831 | ERR1725987 | sewage | Riga, Latvia | co | Metagenome | MetaG | Latvia |
| 139 | PRJEB13831 | ERR1713373 | sewage | Riga, Latvia | co | Metagenome | MetaG | Latvia |
| 139 | PRJEB13831 | ERR2592336 | sewage | Riga, Latvia | co | Metagenome | MetaG | Latvia |
| 139 | PRJEB13831 | ERR2592262 | sewage | Riga, Latvia | co | Metagenome | MetaG | Latvia |
| 140 | PRJEB13831 | ERR1713367 | sewage | Rome, Italy | single | Metagenome | MetaG | Italy |
| 141 | PRJEB13831 | ERR1713353 | sewage | San Cristobal, | co | Metagenome | MetaG | Ecuador |
| 141 | PRJEB13831 | ERR2592333 | sewage | San Cristobal, | co | Metagenome | MetaG | Ecuador |
| 142 | PRJEB13831 | ERR1726011 | sewage | Seattle, WA, | co | Metagenome | MetaG | USA |
| 142 | PRJEB13831 | ERR1713398 | sewage | Seattle, WA, | co | Metagenome | MetaG | USA |
| 142 | PRJEB13831 | ERR2592274 | sewage | Seattle, WA, | co | Metagenome | MetaG | USA |
| 143 | PRJEB13831 | ERR1726001 | sewage | Singapore, Si | co | Metagenome | MetaG | Singapore |
| 143 | PRJEB13831 | ERR1713387 | sewage | Singapore, Si | co | Metagenome | MetaG | Singapore |

| | | | | | | | | |
|---|---|---|---|---|---|---|---|---|
| 143 | PRJEB13831 | ERR2592269 | sewage | Singapore, Si | co | Metagenome | MetaG | Singapore |
| 144 | PRJEB13831 | ERR1725991 | sewage | Skopje, Repul | co | Metagenome | MetaG | Republic of Macedonia |
| 144 | PRJEB13831 | ERR1725990 | sewage | Skopje, Repul | co | Metagenome | MetaG | Republic of Macedonia |
| 144 | PRJEB13831 | ERR1713375 | sewage | Skopje, Repul | co | Metagenome | MetaG | Republic of Macedonia |
| 144 | PRJEB13831 | ERR2592263 | sewage | Skopje, Repul | co | Metagenome | MetaG | Republic of Macedonia |
| 145 | PRJEB13831 | ERR1725948 | sewage | Sofia, Bulgari | co | Metagenome | MetaG | Bulgaria |
| 145 | PRJEB13831 | ERR1725947 | sewage | Sofia, Bulgari | co | Metagenome | MetaG | Bulgaria |
| 145 | PRJEB13831 | ERR1713335 | sewage | Sofia, Bulgari | co | Metagenome | MetaG | Bulgaria |
| 145 | PRJEB13831 | ERR2592247 | sewage | Sofia, Bulgari | co | Metagenome | MetaG | Bulgaria |
| 146 | PRJEB13831 | ERR1713405 | sewage | South Adams | single | Metagenome | MetaG | USA |
| 147 | PRJEB13831 | ERR1725993 | sewage | St. Venera, M | co | Metagenome | MetaG | Malta |
| 147 | PRJEB13831 | ERR1725992 | sewage | St. Venera, M | co | Metagenome | MetaG | Malta |
| 147 | PRJEB13831 | ERR1713376 | sewage | St. Venera, M | co | Metagenome | MetaG | Malta |
| 148 | PRJEB13831 | ERR1713358 | sewage | Tamale, Ghai | single | Metagenome | MetaG | Ghana |
| 149 | PRJEB13831 | ERR1725975 | sewage | Tbilisi, Georg | co | Metagenome | MetaG | Georgia |
| 149 | PRJEB13831 | ERR1725974 | sewage | Tbilisi, Georg | co | Metagenome | MetaG | Georgia |
| 149 | PRJEB13831 | ERR1713357 | sewage | Tbilisi, Georg | co | Metagenome | MetaG | Georgia |
| 149 | PRJEB13831 | ERR2592256 | sewage | Tbilisi, Georg | co | Metagenome | MetaG | Georgia |
| 150 | PRJEB13831 | ERR1725981 | sewage | Tehran, Iran | co | Metagenome | MetaG | Iran |
| 150 | PRJEB13831 | ERR1713364 | sewage | Tehran, Iran | co | Metagenome | MetaG | Iran |
| 150 | PRJEB13831 | ERR2592259 | sewage | Tehran, Iran | co | Metagenome | MetaG | Iran |
| 151 | PRJEB13831 | ERR1725938 | sewage | Tirana, Alban | co | Metagenome | MetaG | Albania |
| 151 | PRJEB13831 | ERR1713331 | sewage | Tirana, Alban | co | Metagenome | MetaG | Albania |
| 151 | PRJEB13831 | ERR2592244 | sewage | Tirana, Alban | co | Metagenome | MetaG | Albania |
| 152 | PRJEB13831 | ERR1725960 | sewage | Toronto, Cana | co | Metagenome | MetaG | Canada |
| 152 | PRJEB13831 | ERR1725959 | sewage | Toronto, Cana | co | Metagenome | MetaG | Canada |
| 152 | PRJEB13831 | ERR1713341 | sewage | Toronto, Cana | co | Metagenome | MetaG | Canada |
| 152 | PRJEB13831 | ERR2592330 | sewage | Toronto, Cana | co | Metagenome | MetaG | Canada |
| 152 | PRJEB13831 | ERR2592250 | sewage | Toronto, Cana | co | Metagenome | MetaG | Canada |
| 153 | PRJEB13831 | ERR1713391 | sewage | Uppsala, Swe | single | Metagenome | MetaG | Sweden |
| 154 | PRJEB13831 | ERR1725946 | sewage | Vienna, Austr | co | Metagenome | MetaG | Austria |
| 154 | PRJEB13831 | ERR1725945 | sewage | Vienna, Austr | co | Metagenome | MetaG | Austria |
| 154 | PRJEB13831 | ERR1713334 | sewage | Vienna, Austr | co | Metagenome | MetaG | Austria |
| 154 | PRJEB13831 | ERR2592246 | sewage | Vienna, Austr | co | Metagenome | MetaG | Austria |
| 155 | PRJEB13831 | ERR1725940 | sewage | Woden, Austr | co | Metagenome | MetaG | Australia |
| 155 | PRJEB13831 | ERR1725939 | sewage | Woden, Austr | co | Metagenome | MetaG | Australia |
| 155 | PRJEB13831 | ERR1713332 | sewage | Woden, Austr | co | Metagenome | MetaG | Australia |
| 155 | PRJEB13831 | ERR2592328 | sewage | Woden, Austr | co | Metagenome | MetaG | Australia |
| 155 | PRJEB13831 | ERR2592245 | sewage | Woden, Austr | co | Metagenome | MetaG | Australia |
| 156 | PRJEB13831 | ERR1725977 | sewage | Zagreb, Croat | co | Metagenome | MetaG | Croatia |
| 156 | PRJEB13831 | ERR1725976 | sewage | Zagreb, Croat | co | Metagenome | MetaG | Croatia |
| 156 | PRJEB13831 | ERR1713360 | sewage | Zagreb, Croat | co | Metagenome | MetaG | Croatia |
| 156 | PRJEB13831 | ERR2592335 | sewage | Zagreb, Croat | co | Metagenome | MetaG | Croatia |
| 156 | PRJEB13831 | ERR2592257 | sewage | Zagreb, Croat | co | Metagenome | MetaG | Croatia |
| 157 | PRJNA244282 | SRR1237780 | sewage | SW Germany | co | Metagenome | MetaG | Germany |
| 157 | PRJNA244282 | SRR1237782 | sewage | SW Germany | co | Metagenome | MetaG | Germany |
| 157 | PRJNA244282 | SRR1237781 | sewage | SW Germany | co | Metagenome | MetaG | Germany |
| 157 | PRJNA244282 | SRR1237783 | sewage | SW Germany | co | Metagenome | MetaG | Germany |
| 158 | PRJNA287481 | SRR5279631 | sewage (deep shale well) | China | single | Metagenome | MetaG | China |
| 159 | PRJNA348753 | SRR5279630 | water | China | single | Metagenome | MetaG | China |
| 160 | PRJNA423296 | SRR6429796 | wastewater river | Kampala, Uga | co | Metagenome | MetaG | Uganda |
| 160 | PRJNA423296 | SRR6429797 | wastewater river | Kampala, Uga | co | Metagenome | MetaG | Uganda |
| 160 | PRJNA423296 | SRR6429787 | wastewater river | Kampala, Uga | co | Metagenome | MetaG | Uganda |
| 160 | PRJNA423296 | SRR6429786 | wastewater river | Kampala, Uga | co | Metagenome | MetaG | Uganda |
| 160 | PRJNA423296 | SRR6429788 | wastewater river | Kampala, Uga | co | Metagenome | MetaG | Uganda |
| 160 | PRJNA423296 | SRR6429789 | wastewater river | Kampala, Uga | co | Metagenome | MetaG | Uganda |
| 160 | PRJNA423296 | SRR6429791 | wastewater river | Kampala, Uga | co | Metagenome | MetaG | Uganda |
| 160 | PRJNA423296 | SRR6429785 | wastewater river | Kampala, Uga | co | Metagenome | MetaG | Uganda |
| 160 | PRJNA423296 | SRR6429784 | wastewater river | Kampala, Uga | co | Metagenome | MetaG | Uganda |
| 160 | PRJNA423296 | SRR6429790 | wastewater river | Kampala, Uga | co | Metagenome | MetaG | Uganda |
| 160 | PRJNA423296 | SRR6429794 | wastewater river | Kampala, Uga | co | Metagenome | MetaG | Uganda |
| 160 | PRJNA423296 | SRR6429795 | wastewater river | Kampala, Uga | co | Metagenome | MetaG | Uganda |
| 160 | PRJNA423296 | SRR6429793 | wastewater river | Kampala, Uga | co | Metagenome | MetaG | Uganda |
| 160 | PRJNA423296 | SRR6429792 | wastewater river | Kampala, Uga | co | Metagenome | MetaG | Uganda |
| 160 | PRJNA423296 | SRR6429805 | wastewater river | Kampala, Uga | co | Metagenome | MetaG | Uganda |
| 160 | PRJNA423296 | SRR6429804 | wastewater river | Kampala, Uga | co | Metagenome | MetaG | Uganda |
| 160 | PRJNA423296 | SRR6429802 | wastewater river | Kampala, Uga | co | Metagenome | MetaG | Uganda |
| 160 | PRJNA423296 | SRR6429803 | wastewater river | Kampala, Uga | co | Metagenome | MetaG | Uganda |
| 160 | PRJNA423296 | SRR6429799 | wastewater river | Kampala, Uga | co | Metagenome | MetaG | Uganda |
| 160 | PRJNA423296 | SRR6429798 | wastewater river | Kampala, Uga | co | Metagenome | MetaG | Uganda |
| 160 | PRJNA423296 | SRR6429800 | wastewater river | Kampala, Uga | co | Metagenome | MetaG | Uganda |
| 160 | PRJNA423296 | SRR6429801 | wastewater river | Kampala, Uga | co | Metagenome | MetaG | Uganda |
| 161 | PRJNA482680 | SRR9030455 | sewage | Calgary, Cana | single | Metagenome | MetaG | Canada |
| 162 | PRJNA488992 | SRR8584358 | wastewater drain | Najafgarf dra | co | Metagenome | MetaG | India |
| 162 | PRJNA488992 | SRR8584355 | wastewater drain | Najafgarf dra | co | Metagenome | MetaG | India |

| | | | | | | | | |
|---|---|---|---|---|---|---|---|---|
| 163 | PRJNA505617 | SRR8583493 | wwtp influent | China | single | Metagenome | MetaG | China |
| 164 | PRJNA505617 | SRR8208347 | wwtp influent | China | single | Metagenome | MetaG | China |
| 165 | PRJNA505617 | SRR8208344 | wwtp influent | Shekwuhui, C | single | Metagenome | MetaG | China |
| 166 | PRJNA505617 | SRR8208343 | wwtp influent | China | single | Metagenome | MetaG | China |
| 167 | PRJNA505617 | SRR14455375 | wwtp influent | China | single | Metagenome | MetaG | China |
| 168 | PRJNA529711 | SRR9030906 | bovine feedlot catch-basin | Calgary, Cana | single | Metagenome | MetaG | Canada |
| 169 | PRJNA529711 | SRR9030900 | bovine feedlot catch-basin | Calgary, Cana | single | Metagenome | MetaG | Canada |
| 170 | PRJNA529711 | SRR9030911 | bovine feedlot catch-basin | Calgary, Cana | single | Metagenome | MetaG | Canada |
| 171 | PRJNA529711 | SRR9030912 | bovine feedlot catch-basin | Calgary, Cana | single | Metagenome | MetaG | Canada |
| 172 | PRJNA529711 | SRR9030908 | bovine feedlot catch-basin | Calgary, Cana | single | Metagenome | MetaG | Canada |
| 173 | PRJNA529711 | SRR9030902 | bovine feedlot catch-basin | Calgary, Cana | single | Metagenome | MetaG | Canada |
| 174 | PRJNA529711 | SRR9030901 | bovine feedlot catch–basin | Calgary, Cana | single | Metagenome | MetaG | Canada |
| 175 | PRJNA529711 | SRR9030907 | bovine feedlot catch-basin | Calgary, Cana | single | Metagenome | MetaG | Canada |
| 176 | PRJNA529711 | SRR9030903 | bovine feedlot catch-basin | Calgary, Cana | single | Metagenome | MetaG | Canada |
| 177 | PRJNA529711 | SRR9030909 | bovine feedlot catch-basin | Calgary, Cana | single | Metagenome | MetaG | Canada |
| 178 | PRJNA529711 | SRR9030910 | bovine feedlot catch-basin | Calgary, Cana | single | Metagenome | MetaG | Canada |
| 179 | PRJNA529711 | SRR9030904 | bovine feedlot catch-basin | Calgary, Cana | single | Metagenome | MetaG | Canada |
| 180 | PRJNA358310 | SRR5123285 | agriculture wastewater | Burkina Faso, | co | Metagenome | MetaG | Africa |
| 180 | PRJNA358310 | SRR5123286 | agriculture wastewater | Burkina Faso, | co | Metagenome | MetaG | Africa |
| 180 | PRJNA358310 | SRR5123284 | agriculture wastewater | Burkina Faso, | co | Metagenome | MetaG | Africa |
| 180 | PRJNA358310 | SRR5123283 | agriculture wastewater | Burkina Faso, | co | Metagenome | MetaG | Africa |
| 180 | PRJNA358310 | SRR5123279 | agriculture wastewater | Burkina Faso, | co | Metagenome | MetaG | Africa |
| 180 | PRJNA358310 | SRR5123280 | agriculture wastewater | Burkina Faso, | co | Metagenome | MetaG | Africa |
| 180 | PRJNA358310 | SRR5123282 | agriculture wastewater | Burkina Faso, | co | Metagenome | MetaG | Africa |
| 180 | PRJNA358310 | SRR5123281 | agriculture wastewater | Burkina Faso, | co | Metagenome | MetaG | Africa |
| 180 | PRJNA358310 | SRR5123289 | agriculture wastewater | Burkina Faso, | co | Metagenome | MetaG | Africa |
| 180 | PRJNA358310 | SRR5123290 | agriculture wastewater | Burkina Faso, | co | Metagenome | MetaG | Africa |
| 180 | PRJNA358310 | SRR5123292 | agriculture wastewater | Burkina Faso, | co | Metagenome | MetaG | Africa |
| 180 | PRJNA358310 | SRR5123278 | agriculture wastewater | Burkina Faso, | co | Metagenome | MetaG | Africa |
| 180 | PRJNA358310 | SRR5123291 | agriculture wastewater | Burkina Faso, | co | Metagenome | MetaG | Africa |
| 180 | PRJNA358310 | SRR5123295 | agriculture wastewater | Burkina Faso, | co | Metagenome | MetaG | Africa |
| 180 | PRJNA358310 | SRR5123288 | agriculture wastewater | Burkina Faso, | co | Metagenome | MetaG | Africa |
| 180 | PRJNA358310 | SRR5123287 | agriculture wastewater | Burkina Faso, | co | Metagenome | MetaG | Africa |
| 180 | PRJNA358310 | SRR5123294 | agriculture wastewater | Burkina Faso, | co | Metagenome | MetaG | Africa |
| 180 | PRJNA358310 | SRR5123293 | agriculture wastewater | Burkina Faso, | co | Metagenome | MetaG | Africa |
| 181 | PRJNA482680 | SRR9030454 | sewage | Calgary, Cana | single | Metagenome | MetaG | Canada |
| 182 | PRJNA482680 | SRR9030456 | sewage | Calgary, Cana | single | Metagenome | MetaG | Canada |
| 183 | PRJNA482680 | SRR9030457 | sewage | Calgary, Cana | single | Metagenome | MetaG | Canada |
| 184 | PRJNA482680 | SRR9030452 | sewage | Calgary, Cana | single | Metagenome | MetaG | Canada |
| 185 | PRJNA482680 | SRR9030453 | sewage | Calgary, Cana | single | Metagenome | MetaG | Canada |
| 186 | PRJNA573903 | SRR10178538 | wastewater | China | single | Metagenome | MetaG | China |
| 187 | PRJNA573903 | SRR10178536 | wastewater | China | single | Metagenome | MetaG | China |
| 188 | PRJNA573903 | SRR10178537 | wastewater | China | single | Metagenome | MetaG | China |
| 189 | PRJNA591725 | SRR10769898 | pig farm wastewater | China | single | Metagenome | MetaG | China |
| 190 | PRJNA591725 | SRR10769904 | pig farm wastewater | China | single | Metagenome | MetaG | China |
| 191 | PRJNA591725 | SRR10769914 | pig farm wastewater | China | single | Metagenome | MetaG | China |
| 192 | PRJNA673783 | SRR12978842 | wwtp influent | Spain | single | Metagenome | MetaG | Spain |
| 193 | PRJNA673783 | SRR12978838 | wwtp influent | Spain | single | Metagenome | MetaG | Spain |
| 194 | PRJNA673783 | SRR12978840 | wwtp influent | Spain | single | Metagenome | MetaG | Spain |
| 195 | PRJNA673783 | SRR12978828 | wwtp influent | Spain | single | Metagenome | MetaG | Spain |
| 196 | PRJNA673783 | SRR12978836 | wwtp influent | Spain | single | Metagenome | MetaG | Spain |
| 197 | PRJNA673783 | SRR12978830 | wwtp influent | Spain | single | Metagenome | MetaG | Spain |
| 198 | PRJNA673783 | SRR12978834 | wwtp influent | Spain | single | Metagenome | MetaG | Spain |
| 199 | PRJNA673783 | SRR12978832 | wwtp influent | Spain | single | Metagenome | MetaG | Spain |
| 200 | PRJNA723368 | SRR14297772 | hospital ww | Luzhou city, C | co | Metagenome | MetaG | China |
| 200 | PRJNA723368 | SRR14297780 | hospital ww | Luzhou city, C | co | Metagenome | MetaG | China |
| 200 | PRJNA723368 | SRR14297777 | hospital ww | Luzhou city, C | co | Metagenome | MetaG | China |
| 200 | PRJNA723368 | SRR14297778 | hospital ww | Luzhou city, C | co | Metagenome | MetaG | China |
| 200 | PRJNA723368 | SRR14297776 | hospital ww | Luzhou city, C | co | Metagenome | MetaG | China |
| 200 | PRJNA723368 | SRR14297775 | hospital ww | Luzhou city, C | co | Metagenome | MetaG | China |
| 200 | PRJNA723368 | SRR14297773 | hospital ww | Luzhou city, C | co | Metagenome | MetaG | China |
| 200 | PRJNA723368 | SRR14297774 | hospital ww | Luzhou city, C | co | Metagenome | MetaG | China |
| 201 | PRJNA770854 | SRR16308658 | hospital ww | Wenzhou, Ch | co | Metagenome | MetaG | China |
| 201 | PRJNA770854 | SRR16308654 | hospital ww | Wenzhou, Ch | co | Metagenome | MetaG | China |
| 201 | PRJNA770854 | SRR16308657 | hospital ww | Wenzhou, Ch | co | Metagenome | MetaG | China |
| 201 | PRJNA770854 | SRR16308649 | hospital ww | Wenzhou, Ch | co | Metagenome | MetaG | China |
| 201 | PRJNA770854 | SRR16308648 | hospital ww | Wenzhou, Ch | co | Metagenome | MetaG | China |
| 201 | PRJNA770854 | SRR16308650 | hospital ww | Wenzhou, Ch | co | Metagenome | MetaG | China |
| 202 | PRJEB34633 | ERR3562950 | sewage | Copenhagen, | co | Metagenome | MetaG | Denmark |
| 202 | PRJEB34633 | ERR3562976 | sewage | Copenhagen, | co | Metagenome | MetaG | Denmark |
| 202 | PRJEB34633 | ERR3562962 | sewage | Copenhagen, | co | Metagenome | MetaG | Denmark |
| 202 | PRJEB34633 | ERR3563030 | sewage | Copenhagen, | co | Metagenome | MetaG | Denmark |
| 202 | PRJEB34633 | ERR3563002 | sewage | Copenhagen, | co | Metagenome | MetaG | Denmark |
| 202 | PRJEB34633 | ERR3563016 | sewage | Copenhagen, | co | Metagenome | MetaG | Denmark |
| 202 | PRJEB34633 | ERR3563015 | sewage | Copenhagen, | co | Metagenome | MetaG | Denmark |

| | | | | | | | |
|---|---|---|---|---|---|---|---|
| 202 | PRJEB34633 | ERR3563001 | sewage | Copenhagen, co | Metagenome | MetaG | Denmark |
| 202 | PRJEB34633 | ERR3563029 | sewage | Copenhagen, co | Metagenome | MetaG | Denmark |
| 202 | PRJEB34633 | ERR3562961 | sewage | Copenhagen, co | Metagenome | MetaG | Denmark |
| 202 | PRJEB34633 | ERR3562975 | sewage | Copenhagen, co | Metagenome | MetaG | Denmark |
| 202 | PRJEB34633 | ERR3562949 | sewage | Copenhagen, co | Metagenome | MetaG | Denmark |
| 202 | PRJEB34633 | ERR3562839 | sewage | Copenhagen, co | Metagenome | MetaG | Denmark |
| 202 | PRJEB34633 | ERR3562963 | sewage | Copenhagen, co | Metagenome | MetaG | Denmark |
| 202 | PRJEB34633 | ERR3562977 | sewage | Copenhagen, co | Metagenome | MetaG | Denmark |
| 202 | PRJEB34633 | ERR3562990 | sewage | Copenhagen, co | Metagenome | MetaG | Denmark |
| 202 | PRJEB34633 | ERR3563017 | sewage | Copenhagen, co | Metagenome | MetaG | Denmark |
| 202 | PRJEB34633 | ERR3563003 | sewage | Copenhagen, co | Metagenome | MetaG | Denmark |
| 202 | PRJEB34633 | ERR3563004 | sewage | Copenhagen, co | Metagenome | MetaG | Denmark |
| 202 | PRJEB34633 | ERR3563018 | sewage | Copenhagen, co | Metagenome | MetaG | Denmark |
| 202 | PRJEB34633 | ERR3562989 | sewage | Copenhagen, co | Metagenome | MetaG | Denmark |
| 202 | PRJEB34633 | ERR3562978 | sewage | Copenhagen, co | Metagenome | MetaG | Denmark |
| 202 | PRJEB34633 | ERR3562964 | sewage | Copenhagen, co | Metagenome | MetaG | Denmark |
| 202 | PRJEB34633 | ERR3562840 | sewage | Copenhagen, co | Metagenome | MetaG | Denmark |
| 202 | PRJEB34633 | ERR3562968 | sewage | Copenhagen, co | Metagenome | MetaG | Denmark |
| 202 | PRJEB34633 | ERR3562974 | sewage | Copenhagen, co | Metagenome | MetaG | Denmark |
| 202 | PRJEB34633 | ERR3562999 | sewage | Copenhagen, co | Metagenome | MetaG | Denmark |
| 202 | PRJEB34633 | ERR3563014 | sewage | Copenhagen, co | Metagenome | MetaG | Denmark |
| 202 | PRJEB34633 | ERR3563008 | sewage | Copenhagen, co | Metagenome | MetaG | Denmark |
| 202 | PRJEB34633 | ERR3563007 | sewage | Copenhagen, co | Metagenome | MetaG | Denmark |
| 202 | PRJEB34633 | ERR3563013 | sewage | Copenhagen, co | Metagenome | MetaG | Denmark |
| 202 | PRJEB34633 | ERR3563000 | sewage | Copenhagen, co | Metagenome | MetaG | Denmark |
| 202 | PRJEB34633 | ERR3562973 | sewage | Copenhagen, co | Metagenome | MetaG | Denmark |
| 202 | PRJEB34633 | ERR3562967 | sewage | Copenhagen, co | Metagenome | MetaG | Denmark |
| 202 | PRJEB34633 | ERR3562971 | sewage | Copenhagen, co | Metagenome | MetaG | Denmark |
| 202 | PRJEB34633 | ERR3562965 | sewage | Copenhagen, co | Metagenome | MetaG | Denmark |
| 202 | PRJEB34633 | ERR3562959 | sewage | Copenhagen, co | Metagenome | MetaG | Denmark |
| 202 | PRJEB34633 | ERR3563005 | sewage | Copenhagen, co | Metagenome | MetaG | Denmark |
| 202 | PRJEB34633 | ERR3563011 | sewage | Copenhagen, co | Metagenome | MetaG | Denmark |
| 202 | PRJEB34633 | ERR3563039 | sewage | Copenhagen, co | Metagenome | MetaG | Denmark |
| 202 | PRJEB34633 | ERR3563040 | sewage | Copenhagen, co | Metagenome | MetaG | Denmark |
| 202 | PRJEB34633 | ERR3563012 | sewage | Copenhagen, co | Metagenome | MetaG | Denmark |
| 202 | PRJEB34633 | ERR3563006 | sewage | Copenhagen, co | Metagenome | MetaG | Denmark |
| 202 | PRJEB34633 | ERR3562960 | sewage | Copenhagen, co | Metagenome | MetaG | Denmark |
| 202 | PRJEB34633 | ERR3562966 | sewage | Copenhagen, co | Metagenome | MetaG | Denmark |
| 202 | PRJEB34633 | ERR3562972 | sewage | Copenhagen, co | Metagenome | MetaG | Denmark |
| 202 | PRJEB34633 | ERR3562859 | sewage | Copenhagen, co | Metagenome | MetaG | Denmark |
| 202 | PRJEB34633 | ERR3562871 | sewage | Copenhagen, co | Metagenome | MetaG | Denmark |
| 202 | PRJEB34633 | ERR3562917 | sewage | Copenhagen, co | Metagenome | MetaG | Denmark |
| 202 | PRJEB34633 | ERR3562903 | sewage | Copenhagen, co | Metagenome | MetaG | Denmark |
| 202 | PRJEB34633 | ERR3562865 | sewage | Copenhagen, co | Metagenome | MetaG | Denmark |
| 202 | PRJEB34633 | ERR3563101 | sewage | Copenhagen, co | Metagenome | MetaG | Denmark |
| 202 | PRJEB34633 | ERR3563063 | sewage | Copenhagen, co | Metagenome | MetaG | Denmark |
| 202 | PRJEB34633 | ERR3563077 | sewage | Copenhagen, co | Metagenome | MetaG | Denmark |
| 202 | PRJEB34633 | ERR3563090 | sewage | Copenhagen, co | Metagenome | MetaG | Denmark |
| 202 | PRJEB34633 | ERR3563089 | sewage | Copenhagen, co | Metagenome | MetaG | Denmark |
| 202 | PRJEB34633 | ERR3563078 | sewage | Copenhagen, co | Metagenome | MetaG | Denmark |
| 202 | PRJEB34633 | ERR3563064 | sewage | Copenhagen, co | Metagenome | MetaG | Denmark |
| 202 | PRJEB34633 | ERR3563102 | sewage | Copenhagen, co | Metagenome | MetaG | Denmark |
| 202 | PRJEB34633 | ERR3562904 | sewage | Copenhagen, co | Metagenome | MetaG | Denmark |
| 202 | PRJEB34633 | ERR3562866 | sewage | Copenhagen, co | Metagenome | MetaG | Denmark |
| 202 | PRJEB34633 | ERR3562872 | sewage | Copenhagen, co | Metagenome | MetaG | Denmark |
| 202 | PRJEB34633 | ERR3562918 | sewage | Copenhagen, co | Metagenome | MetaG | Denmark |
| 202 | PRJEB34633 | ERR3562860 | sewage | Copenhagen, co | Metagenome | MetaG | Denmark |
| 202 | PRJEB34633 | ERR3562930 | sewage | Copenhagen, co | Metagenome | MetaG | Denmark |
| 202 | PRJEB34633 | ERR3562868 | sewage | Copenhagen, co | Metagenome | MetaG | Denmark |
| 202 | PRJEB34633 | ERR3562902 | sewage | Copenhagen, co | Metagenome | MetaG | Denmark |
| 202 | PRJEB34633 | ERR3562916 | sewage | Copenhagen, co | Metagenome | MetaG | Denmark |
| 202 | PRJEB34633 | ERR3562874 | sewage | Copenhagen, co | Metagenome | MetaG | Denmark |
| 202 | PRJEB34633 | ERR3562899 | sewage | Copenhagen, co | Metagenome | MetaG | Denmark |
| 202 | PRJEB34633 | ERR3563104 | sewage | Copenhagen, co | Metagenome | MetaG | Denmark |
| 202 | PRJEB34633 | ERR3563050 | sewage | Copenhagen, co | Metagenome | MetaG | Denmark |
| 202 | PRJEB34633 | ERR3563076 | sewage | Copenhagen, co | Metagenome | MetaG | Denmark |
| 202 | PRJEB34633 | ERR3563062 | sewage | Copenhagen, co | Metagenome | MetaG | Denmark |
| 202 | PRJEB34633 | ERR3563061 | sewage | Copenhagen, co | Metagenome | MetaG | Denmark |
| 202 | PRJEB34633 | ERR3563075 | sewage | Copenhagen, co | Metagenome | MetaG | Denmark |
| 202 | PRJEB34633 | ERR3563049 | sewage | Copenhagen, co | Metagenome | MetaG | Denmark |
| 202 | PRJEB34633 | ERR3563103 | sewage | Copenhagen, co | Metagenome | MetaG | Denmark |
| 202 | PRJEB34633 | ERR3562900 | sewage | Copenhagen, co | Metagenome | MetaG | Denmark |
| 202 | PRJEB34633 | ERR3562915 | sewage | Copenhagen, co | Metagenome | MetaG | Denmark |
| 202 | PRJEB34633 | ERR3562873 | sewage | Copenhagen, co | Metagenome | MetaG | Denmark |
| 202 | PRJEB34633 | ERR3562867 | sewage | Copenhagen, co | Metagenome | MetaG | Denmark |

| | | | | | | | |
|---|---|---|---|---|---|---|---|
| 202 | PRJEB34633 | ERR3562901 | sewage | Copenhagen, co | Metagenome | MetaG | Denmark |
| 202 | PRJEB34633 | ERR3562929 | sewage | Copenhagen, co | Metagenome | MetaG | Denmark |
| 202 | PRJEB34633 | ERR3562905 | sewage | Copenhagen, co | Metagenome | MetaG | Denmark |
| 202 | PRJEB34633 | ERR3562863 | sewage | Copenhagen, co | Metagenome | MetaG | Denmark |
| 202 | PRJEB34633 | ERR3562877 | sewage | Copenhagen, co | Metagenome | MetaG | Denmark |
| 202 | PRJEB34633 | ERR3562911 | sewage | Copenhagen, co | Metagenome | MetaG | Denmark |
| 202 | PRJEB34633 | ERR3562939 | sewage | Copenhagen, co | Metagenome | MetaG | Denmark |
| 202 | PRJEB34633 | ERR3562890 | sewage | Copenhagen, co | Metagenome | MetaG | Denmark |
| 202 | PRJEB34633 | ERR3563107 | sewage | Copenhagen, co | Metagenome | MetaG | Denmark |
| 202 | PRJEB34633 | ERR3563071 | sewage | Copenhagen, co | Metagenome | MetaG | Denmark |
| 202 | PRJEB34633 | ERR3563065 | sewage | Copenhagen, co | Metagenome | MetaG | Denmark |
| 202 | PRJEB34633 | ERR3563059 | sewage | Copenhagen, co | Metagenome | MetaG | Denmark |
| 202 | PRJEB34633 | ERR3563060 | sewage | Copenhagen, co | Metagenome | MetaG | Denmark |
| 202 | PRJEB34633 | ERR3563066 | sewage | Copenhagen, co | Metagenome | MetaG | Denmark |
| 202 | PRJEB34633 | ERR3563072 | sewage | Copenhagen, co | Metagenome | MetaG | Denmark |
| 202 | PRJEB34633 | ERR3563108 | sewage | Copenhagen, co | Metagenome | MetaG | Denmark |
| 202 | PRJEB34633 | ERR3562889 | sewage | Copenhagen, co | Metagenome | MetaG | Denmark |
| 202 | PRJEB34633 | ERR3562940 | sewage | Copenhagen, co | Metagenome | MetaG | Denmark |
| 202 | PRJEB34633 | ERR3562878 | sewage | Copenhagen, co | Metagenome | MetaG | Denmark |
| 202 | PRJEB34633 | ERR3562912 | sewage | Copenhagen, co | Metagenome | MetaG | Denmark |
| 202 | PRJEB34633 | ERR3562906 | sewage | Copenhagen, co | Metagenome | MetaG | Denmark |
| 202 | PRJEB34633 | ERR3562864 | sewage | Copenhagen, co | Metagenome | MetaG | Denmark |
| 202 | PRJEB34633 | ERR3562914 | sewage | Copenhagen, co | Metagenome | MetaG | Denmark |
| 202 | PRJEB34633 | ERR3562876 | sewage | Copenhagen, co | Metagenome | MetaG | Denmark |
| 202 | PRJEB34633 | ERR3562862 | sewage | Copenhagen, co | Metagenome | MetaG | Denmark |
| 202 | PRJEB34633 | ERR3562908 | sewage | Copenhagen, co | Metagenome | MetaG | Denmark |
| 202 | PRJEB34633 | ERR3562850 | sewage | Copenhagen, co | Metagenome | MetaG | Denmark |
| 202 | PRJEB34633 | ERR3563112 | sewage | Copenhagen, co | Metagenome | MetaG | Denmark |
| 202 | PRJEB34633 | ERR3563106 | sewage | Copenhagen, co | Metagenome | MetaG | Denmark |
| 202 | PRJEB34633 | ERR3563068 | sewage | Copenhagen, co | Metagenome | MetaG | Denmark |
| 202 | PRJEB34633 | ERR3563074 | sewage | Copenhagen, co | Metagenome | MetaG | Denmark |
| 202 | PRJEB34633 | ERR3563099 | sewage | Copenhagen, co | Metagenome | MetaG | Denmark |
| 202 | PRJEB34633 | ERR3563100 | sewage | Copenhagen, co | Metagenome | MetaG | Denmark |
| 202 | PRJEB34633 | ERR3563073 | sewage | Copenhagen, co | Metagenome | MetaG | Denmark |
| 202 | PRJEB34633 | ERR3563067 | sewage | Copenhagen, co | Metagenome | MetaG | Denmark |
| 202 | PRJEB34633 | ERR3563105 | sewage | Copenhagen, co | Metagenome | MetaG | Denmark |
| 202 | PRJEB34633 | ERR3563111 | sewage | Copenhagen, co | Metagenome | MetaG | Denmark |
| 202 | PRJEB34633 | ERR3562849 | sewage | Copenhagen, co | Metagenome | MetaG | Denmark |
| 202 | PRJEB34633 | ERR3562861 | sewage | Copenhagen, co | Metagenome | MetaG | Denmark |
| 202 | PRJEB34633 | ERR3562907 | sewage | Copenhagen, co | Metagenome | MetaG | Denmark |
| 202 | PRJEB34633 | ERR3562913 | sewage | Copenhagen, co | Metagenome | MetaG | Denmark |
| 202 | PRJEB34633 | ERR3562875 | sewage | Copenhagen, co | Metagenome | MetaG | Denmark |
| 202 | PRJEB34633 | ERR3562880 | sewage | Copenhagen, co | Metagenome | MetaG | Denmark |
| 202 | PRJEB34633 | ERR3562852 | sewage | Copenhagen, co | Metagenome | MetaG | Denmark |
| 202 | PRJEB34633 | ERR3562938 | sewage | Copenhagen, co | Metagenome | MetaG | Denmark |
| 202 | PRJEB34633 | ERR3562924 | sewage | Copenhagen, co | Metagenome | MetaG | Denmark |
| 202 | PRJEB34633 | ERR3562846 | sewage | Copenhagen, co | Metagenome | MetaG | Denmark |
| 202 | PRJEB34633 | ERR3562893 | sewage | Copenhagen, co | Metagenome | MetaG | Denmark |
| 202 | PRJEB34633 | ERR3562887 | sewage | Copenhagen, co | Metagenome | MetaG | Denmark |
| 202 | PRJEB34633 | ERR3563110 | sewage | Copenhagen, co | Metagenome | MetaG | Denmark |
| 202 | PRJEB34633 | ERR3563044 | sewage | Copenhagen, co | Metagenome | MetaG | Denmark |
| 202 | PRJEB34633 | ERR3563058 | sewage | Copenhagen, co | Metagenome | MetaG | Denmark |
| 202 | PRJEB34633 | ERR3563081 | sewage | Copenhagen, co | Metagenome | MetaG | Denmark |
| 202 | PRJEB34633 | ERR3563095 | sewage | Copenhagen, co | Metagenome | MetaG | Denmark |
| 202 | PRJEB34633 | ERR3563096 | sewage | Copenhagen, co | Metagenome | MetaG | Denmark |
| 202 | PRJEB34633 | ERR3563082 | sewage | Copenhagen, co | Metagenome | MetaG | Denmark |
| 202 | PRJEB34633 | ERR3563057 | sewage | Copenhagen, co | Metagenome | MetaG | Denmark |
| 202 | PRJEB34633 | ERR3563043 | sewage | Copenhagen, co | Metagenome | MetaG | Denmark |
| 202 | PRJEB34633 | ERR3563109 | sewage | Copenhagen, co | Metagenome | MetaG | Denmark |
| 202 | PRJEB34633 | ERR3562888 | sewage | Copenhagen, co | Metagenome | MetaG | Denmark |
| 202 | PRJEB34633 | ERR3562894 | sewage | Copenhagen, co | Metagenome | MetaG | Denmark |
| 202 | PRJEB34633 | ERR3562923 | sewage | Copenhagen, co | Metagenome | MetaG | Denmark |
| 202 | PRJEB34633 | ERR3562845 | sewage | Copenhagen, co | Metagenome | MetaG | Denmark |
| 202 | PRJEB34633 | ERR3562851 | sewage | Copenhagen, co | Metagenome | MetaG | Denmark |
| 202 | PRJEB34633 | ERR3562937 | sewage | Copenhagen, co | Metagenome | MetaG | Denmark |
| 202 | PRJEB34633 | ERR3562879 | sewage | Copenhagen, co | Metagenome | MetaG | Denmark |
| 202 | PRJEB34633 | ERR3562909 | sewage | Copenhagen, co | Metagenome | MetaG | Denmark |
| 202 | PRJEB34633 | ERR3562847 | sewage | Copenhagen, co | Metagenome | MetaG | Denmark |
| 202 | PRJEB34633 | ERR3562921 | sewage | Copenhagen, co | Metagenome | MetaG | Denmark |
| 202 | PRJEB34633 | ERR3562935 | sewage | Copenhagen, co | Metagenome | MetaG | Denmark |
| 202 | PRJEB34633 | ERR3562853 | sewage | Copenhagen, co | Metagenome | MetaG | Denmark |
| 202 | PRJEB34633 | ERR3562886 | sewage | Copenhagen, co | Metagenome | MetaG | Denmark |
| 202 | PRJEB34633 | ERR3562892 | sewage | Copenhagen, co | Metagenome | MetaG | Denmark |
| 202 | PRJEB34633 | ERR3563069 | sewage | Copenhagen, co | Metagenome | MetaG | Denmark |
| 202 | PRJEB34633 | ERR3563055 | sewage | Copenhagen, co | Metagenome | MetaG | Denmark |

| | | | | | | | |
|---|---|---|---|---|---|---|---|
| 202 | PRJEB34633 | ERR3563041 | sewage | Copenhagen, co | Metagenome | MetaG | Denmark |
| 202 | PRJEB34633 | ERR3563098 | sewage | Copenhagen, co | Metagenome | MetaG | Denmark |
| 202 | PRJEB34633 | ERR3563084 | sewage | Copenhagen, co | Metagenome | MetaG | Denmark |
| 202 | PRJEB34633 | ERR3563083 | sewage | Copenhagen, co | Metagenome | MetaG | Denmark |
| 202 | PRJEB34633 | ERR3563097 | sewage | Copenhagen, co | Metagenome | MetaG | Denmark |
| 202 | PRJEB34633 | ERR3563042 | sewage | Copenhagen, co | Metagenome | MetaG | Denmark |
| 202 | PRJEB34633 | ERR3563056 | sewage | Copenhagen, co | Metagenome | MetaG | Denmark |
| 202 | PRJEB34633 | ERR3563070 | sewage | Copenhagen, co | Metagenome | MetaG | Denmark |
| 202 | PRJEB34633 | ERR3562891 | sewage | Copenhagen, co | Metagenome | MetaG | Denmark |
| 202 | PRJEB34633 | ERR3562885 | sewage | Copenhagen, co | Metagenome | MetaG | Denmark |
| 202 | PRJEB34633 | ERR3562936 | sewage | Copenhagen, co | Metagenome | MetaG | Denmark |
| 202 | PRJEB34633 | ERR3562854 | sewage | Copenhagen, co | Metagenome | MetaG | Denmark |
| 202 | PRJEB34633 | ERR3562848 | sewage | Copenhagen, co | Metagenome | MetaG | Denmark |
| 202 | PRJEB34633 | ERR3562922 | sewage | Copenhagen, co | Metagenome | MetaG | Denmark |
| 202 | PRJEB34633 | ERR3562910 | sewage | Copenhagen, co | Metagenome | MetaG | Denmark |
| 202 | PRJEB34633 | ERR3562926 | sewage | Copenhagen, co | Metagenome | MetaG | Denmark |
| 202 | PRJEB34633 | ERR3562844 | sewage | Copenhagen, co | Metagenome | MetaG | Denmark |
| 202 | PRJEB34633 | ERR3562858 | sewage | Copenhagen, co | Metagenome | MetaG | Denmark |
| 202 | PRJEB34633 | ERR3562932 | sewage | Copenhagen, co | Metagenome | MetaG | Denmark |
| 202 | PRJEB34633 | ERR3562920 | sewage | Copenhagen, co | Metagenome | MetaG | Denmark |
| 202 | PRJEB34633 | ERR3562881 | sewage | Copenhagen, co | Metagenome | MetaG | Denmark |
| 202 | PRJEB34633 | ERR3562895 | sewage | Copenhagen, co | Metagenome | MetaG | Denmark |
| 202 | PRJEB34633 | ERR3563052 | sewage | Copenhagen, co | Metagenome | MetaG | Denmark |
| 202 | PRJEB34633 | ERR3563046 | sewage | Copenhagen, co | Metagenome | MetaG | Denmark |
| 202 | PRJEB34633 | ERR3563080 | sewage | Copenhagen, co | Metagenome | MetaG | Denmark |
| 202 | PRJEB34633 | ERR3563093 | sewage | Copenhagen, co | Metagenome | MetaG | Denmark |
| 202 | PRJEB34633 | ERR3563087 | sewage | Copenhagen, co | Metagenome | MetaG | Denmark |
| 202 | PRJEB34633 | ERR3563088 | sewage | Copenhagen, co | Metagenome | MetaG | Denmark |
| 202 | PRJEB34633 | ERR3563094 | sewage | Copenhagen, co | Metagenome | MetaG | Denmark |
| 202 | PRJEB34633 | ERR3563079 | sewage | Copenhagen, co | Metagenome | MetaG | Denmark |
| 202 | PRJEB34633 | ERR3563045 | sewage | Copenhagen, co | Metagenome | MetaG | Denmark |
| 202 | PRJEB34633 | ERR3563051 | sewage | Copenhagen, co | Metagenome | MetaG | Denmark |
| 202 | PRJEB34633 | ERR3562896 | sewage | Copenhagen, co | Metagenome | MetaG | Denmark |
| 202 | PRJEB34633 | ERR3562882 | sewage | Copenhagen, co | Metagenome | MetaG | Denmark |
| 202 | PRJEB34633 | ERR3562919 | sewage | Copenhagen, co | Metagenome | MetaG | Denmark |
| 202 | PRJEB34633 | ERR3562857 | sewage | Copenhagen, co | Metagenome | MetaG | Denmark |
| 202 | PRJEB34633 | ERR3562931 | sewage | Copenhagen, co | Metagenome | MetaG | Denmark |
| 202 | PRJEB34633 | ERR3562925 | sewage | Copenhagen, co | Metagenome | MetaG | Denmark |
| 202 | PRJEB34633 | ERR3562843 | sewage | Copenhagen, co | Metagenome | MetaG | Denmark |
| 202 | PRJEB34633 | ERR3562933 | sewage | Copenhagen, co | Metagenome | MetaG | Denmark |
| 202 | PRJEB34633 | ERR3562855 | sewage | Copenhagen, co | Metagenome | MetaG | Denmark |
| 202 | PRJEB34633 | ERR3562841 | sewage | Copenhagen, co | Metagenome | MetaG | Denmark |
| 202 | PRJEB34633 | ERR3562927 | sewage | Copenhagen, co | Metagenome | MetaG | Denmark |
| 202 | PRJEB34633 | ERR3562869 | sewage | Copenhagen, co | Metagenome | MetaG | Denmark |
| 202 | PRJEB34633 | ERR3562898 | sewage | Copenhagen, co | Metagenome | MetaG | Denmark |
| 202 | PRJEB34633 | ERR3562884 | sewage | Copenhagen, co | Metagenome | MetaG | Denmark |
| 202 | PRJEB34633 | ERR3563047 | sewage | Copenhagen, co | Metagenome | MetaG | Denmark |
| 202 | PRJEB34633 | ERR3563053 | sewage | Copenhagen, co | Metagenome | MetaG | Denmark |
| 202 | PRJEB34633 | ERR3563086 | sewage | Copenhagen, co | Metagenome | MetaG | Denmark |
| 202 | PRJEB34633 | ERR3563092 | sewage | Copenhagen, co | Metagenome | MetaG | Denmark |
| 202 | PRJEB34633 | ERR3563091 | sewage | Copenhagen, co | Metagenome | MetaG | Denmark |
| 202 | PRJEB34633 | ERR3563085 | sewage | Copenhagen, co | Metagenome | MetaG | Denmark |
| 202 | PRJEB34633 | ERR3563054 | sewage | Copenhagen, co | Metagenome | MetaG | Denmark |
| 202 | PRJEB34633 | ERR3563048 | sewage | Copenhagen, co | Metagenome | MetaG | Denmark |
| 202 | PRJEB34633 | ERR3562883 | sewage | Copenhagen, co | Metagenome | MetaG | Denmark |
| 202 | PRJEB34633 | ERR3562897 | sewage | Copenhagen, co | Metagenome | MetaG | Denmark |
| 202 | PRJEB34633 | ERR3562870 | sewage | Copenhagen, co | Metagenome | MetaG | Denmark |
| 202 | PRJEB34633 | ERR3562842 | sewage | Copenhagen, co | Metagenome | MetaG | Denmark |
| 202 | PRJEB34633 | ERR3562928 | sewage | Copenhagen, co | Metagenome | MetaG | Denmark |
| 202 | PRJEB34633 | ERR3562934 | sewage | Copenhagen, co | Metagenome | MetaG | Denmark |
| 202 | PRJEB34633 | ERR3562856 | sewage | Copenhagen, co | Metagenome | MetaG | Denmark |
| 202 | PRJEB34633 | ERR3562969 | sewage | Copenhagen, co | Metagenome | MetaG | Denmark |
| 202 | PRJEB34633 | ERR3562955 | sewage | Copenhagen, co | Metagenome | MetaG | Denmark |
| 202 | PRJEB34633 | ERR3562941 | sewage | Copenhagen, co | Metagenome | MetaG | Denmark |
| 202 | PRJEB34633 | ERR3562998 | sewage | Copenhagen, co | Metagenome | MetaG | Denmark |
| 202 | PRJEB34633 | ERR3562984 | sewage | Copenhagen, co | Metagenome | MetaG | Denmark |
| 202 | PRJEB34633 | ERR3563009 | sewage | Copenhagen, co | Metagenome | MetaG | Denmark |
| 202 | PRJEB34633 | ERR3563021 | sewage | Copenhagen, co | Metagenome | MetaG | Denmark |
| 202 | PRJEB34633 | ERR3563035 | sewage | Copenhagen, co | Metagenome | MetaG | Denmark |
| 202 | PRJEB34633 | ERR3563036 | sewage | Copenhagen, co | Metagenome | MetaG | Denmark |
| 202 | PRJEB34633 | ERR3563022 | sewage | Copenhagen, co | Metagenome | MetaG | Denmark |
| 202 | PRJEB34633 | ERR3563010 | sewage | Copenhagen, co | Metagenome | MetaG | Denmark |
| 202 | PRJEB34633 | ERR3562983 | sewage | Copenhagen, co | Metagenome | MetaG | Denmark |
| 202 | PRJEB34633 | ERR3562997 | sewage | Copenhagen, co | Metagenome | MetaG | Denmark |
| 202 | PRJEB34633 | ERR3562942 | sewage | Copenhagen, co | Metagenome | MetaG | Denmark |

| | | | | | | | | |
|---|---|---|---|---|---|---|---|---|
| 202 | PRJEB34633 | ERR3562956 | sewage | Copenhagen, | co | Metagenome | MetaG | Denmark |
| 202 | PRJEB34633 | ERR3562970 | sewage | Copenhagen, | co | Metagenome | MetaG | Denmark |
| 202 | PRJEB34633 | ERR3562944 | sewage | Copenhagen, | co | Metagenome | MetaG | Denmark |
| 202 | PRJEB34633 | ERR3562958 | sewage | Copenhagen, | co | Metagenome | MetaG | Denmark |
| 202 | PRJEB34633 | ERR3562981 | sewage | Copenhagen, | co | Metagenome | MetaG | Denmark |
| 202 | PRJEB34633 | ERR3562995 | sewage | Copenhagen, | co | Metagenome | MetaG | Denmark |
| 202 | PRJEB34633 | ERR3563038 | sewage | Copenhagen, | co | Metagenome | MetaG | Denmark |
| 202 | PRJEB34633 | ERR3563024 | sewage | Copenhagen, | co | Metagenome | MetaG | Denmark |
| 202 | PRJEB34633 | ERR3563023 | sewage | Copenhagen, | co | Metagenome | MetaG | Denmark |
| 202 | PRJEB34633 | ERR3563037 | sewage | Copenhagen, | co | Metagenome | MetaG | Denmark |
| 202 | PRJEB34633 | ERR3562996 | sewage | Copenhagen, | co | Metagenome | MetaG | Denmark |
| 202 | PRJEB34633 | ERR3562982 | sewage | Copenhagen, | co | Metagenome | MetaG | Denmark |
| 202 | PRJEB34633 | ERR3562957 | sewage | Copenhagen, | co | Metagenome | MetaG | Denmark |
| 202 | PRJEB34633 | ERR3562943 | sewage | Copenhagen, | co | Metagenome | MetaG | Denmark |
| 202 | PRJEB34633 | ERR3562947 | sewage | Copenhagen, | co | Metagenome | MetaG | Denmark |
| 202 | PRJEB34633 | ERR3562953 | sewage | Copenhagen, | co | Metagenome | MetaG | Denmark |
| 202 | PRJEB34633 | ERR3562986 | sewage | Copenhagen, | co | Metagenome | MetaG | Denmark |
| 202 | PRJEB34633 | ERR3562992 | sewage | Copenhagen, | co | Metagenome | MetaG | Denmark |
| 202 | PRJEB34633 | ERR3563033 | sewage | Copenhagen, | co | Metagenome | MetaG | Denmark |
| 202 | PRJEB34633 | ERR3563027 | sewage | Copenhagen, | co | Metagenome | MetaG | Denmark |
| 202 | PRJEB34633 | ERR3563028 | sewage | Copenhagen, | co | Metagenome | MetaG | Denmark |
| 202 | PRJEB34633 | ERR3563034 | sewage | Copenhagen, | co | Metagenome | MetaG | Denmark |
| 202 | PRJEB34633 | ERR3562991 | sewage | Copenhagen, | co | Metagenome | MetaG | Denmark |
| 202 | PRJEB34633 | ERR3562985 | sewage | Copenhagen, | co | Metagenome | MetaG | Denmark |
| 202 | PRJEB34633 | ERR3562954 | sewage | Copenhagen, | co | Metagenome | MetaG | Denmark |
| 202 | PRJEB34633 | ERR3562836 | sewage | Copenhagen, | co | Metagenome | MetaG | Denmark |
| 202 | PRJEB34633 | ERR3562948 | sewage | Copenhagen, | co | Metagenome | MetaG | Denmark |
| 202 | PRJEB34633 | ERR3562838 | sewage | Copenhagen, | co | Metagenome | MetaG | Denmark |
| 202 | PRJEB34633 | ERR3562952 | sewage | Copenhagen, | co | Metagenome | MetaG | Denmark |
| 202 | PRJEB34633 | ERR3562946 | sewage | Copenhagen, | co | Metagenome | MetaG | Denmark |
| 202 | PRJEB34633 | ERR3562980 | sewage | Copenhagen, | co | Metagenome | MetaG | Denmark |
| 202 | PRJEB34633 | ERR3562993 | sewage | Copenhagen, | co | Metagenome | MetaG | Denmark |
| 202 | PRJEB34633 | ERR3562987 | sewage | Copenhagen, | co | Metagenome | MetaG | Denmark |
| 202 | PRJEB34633 | ERR3563026 | sewage | Copenhagen, | co | Metagenome | MetaG | Denmark |
| 202 | PRJEB34633 | ERR3563032 | sewage | Copenhagen, | co | Metagenome | MetaG | Denmark |
| 202 | PRJEB34633 | ERR3563020 | sewage | Copenhagen, | co | Metagenome | MetaG | Denmark |
| 202 | PRJEB34633 | ERR3563019 | sewage | Copenhagen, | co | Metagenome | MetaG | Denmark |
| 202 | PRJEB34633 | ERR3563031 | sewage | Copenhagen, | co | Metagenome | MetaG | Denmark |
| 202 | PRJEB34633 | ERR3563025 | sewage | Copenhagen, | co | Metagenome | MetaG | Denmark |
| 202 | PRJEB34633 | ERR3562988 | sewage | Copenhagen, | co | Metagenome | MetaG | Denmark |
| 202 | PRJEB34633 | ERR3562994 | sewage | Copenhagen, | co | Metagenome | MetaG | Denmark |
| 202 | PRJEB34633 | ERR3562979 | sewage | Copenhagen, | co | Metagenome | MetaG | Denmark |
| 202 | PRJEB34633 | ERR3562945 | sewage | Copenhagen, | co | Metagenome | MetaG | Denmark |
| 202 | PRJEB34633 | ERR3562837 | sewage | Copenhagen, | co | Metagenome | MetaG | Denmark |
| 202 | PRJEB34633 | ERR3562951 | sewage | Copenhagen, | co | Metagenome | MetaG | Denmark |
| 203 | PRJEB13831 | ERR1725973 | sewage | Helsinki, Finla | co | Metagenome | MetaG | Finland |
| 203 | PRJEB13831 | ERR1713356 | sewage | Helsinki, Finla | co | Metagenome | MetaG | Finland |
| 203 | PRJEB13831 | ERR2592255 | sewage | Helsinki, Finla | co | Metagenome | MetaG | Finland |
| 213 | PRJEB27054 | ERR2607605 | sewage | Lusaka, Zamb | co | Metagenome | MetaG | Zambia |
| 213 | PRJEB27054 | ERR2607596 | sewage | Lusaka, Zamb | co | Metagenome | MetaG | Zambia |
| 213 | PRJEB27054 | ERR2607597 | sewage | Lusaka, Zamb | co | Metagenome | MetaG | Zambia |
| 221 | PRJEB27054 | ERR2607604 | sewage | Kitwe, Zambi | co | Metagenome | MetaG | Zambia |
| 221 | PRJEB27054 | ERR2607603 | sewage | Kitwe, Zambi | co | Metagenome | MetaG | Zambia |
| 221 | PRJEB27054 | ERR2607602 | sewage | Kitwe, Zambi | co | Metagenome | MetaG | Zambia |
| 221 | PRJEB27054 | ERR2607601 | sewage | Kitwe, Zambi | co | Metagenome | MetaG | Zambia |
| 221 | PRJEB27054 | ERR2607600 | sewage | Kitwe, Zambi | co | Metagenome | MetaG | Zambia |
| 221 | PRJEB27054 | ERR2607599 | sewage | Kitwe, Zambi | co | Metagenome | MetaG | Zambia |
| 221 | PRJEB27054 | ERR2607598 | sewage | Kitwe, Zambi | co | Metagenome | MetaG | Zambia |
| 214 | PRJEB27054 | ERR2607595 | sewage | Pretoria, Sout | co | Metagenome | MetaG | South Africa |
| 214 | PRJEB27054 | ERR2607594 | sewage | Pretoria, Sout | co | Metagenome | MetaG | South Africa |
| 214 | PRJEB27054 | ERR2607593 | sewage | Pretoria, Sout | co | Metagenome | MetaG | South Africa |
| 215 | PRJEB27054 | ERR2607592 | sewage | Kosovo, Albar | single | Metagenome | MetaG | Albania |
| 216 | PRJEB27054 | ERR2607591 | sewage | Vietnam | single | Metagenome | MetaG | Vietnam |
| 217 | PRJEB27054 | ERR2607590 | sewage | Atlanta, GA, I | single | Metagenome | MetaG | USA |
| 218 | PRJEB27054 | ERR2607589 | sewage | Boulder, CO, | single | Metagenome | MetaG | USA |
| 219 | PRJEB27054 | ERR2607588 | sewage | Adams Count | single | Metagenome | MetaG | USA |
| 220 | PRJEB27054 | ERR2607587 | sewage | El Paso, TX, U | co | Metagenome | MetaG | USA |
| 220 | PRJEB27054 | ERR2607586 | sewage | El Paso, TX, U | co | Metagenome | MetaG | USA |
| 220 | PRJEB27054 | ERR2607585 | sewage | El Paso, TX, U | co | Metagenome | MetaG | USA |
| 220 | PRJEB27054 | ERR2607584 | sewage | El Paso, TX, U | co | Metagenome | MetaG | USA |
| 220 | PRJEB27054 | ERR2607583 | sewage | El Paso, TX, U | co | Metagenome | MetaG | USA |
| 220 | PRJEB27054 | ERR2607582 | sewage | El Paso, TX, U | co | Metagenome | MetaG | USA |
| 220 | PRJEB27054 | ERR2607581 | sewage | El Paso, TX, U | co | Metagenome | MetaG | USA |
| 220 | PRJEB27054 | ERR2607580 | sewage | El Paso, TX, U | co | Metagenome | MetaG | USA |
| 220 | PRJEB27054 | ERR2607579 | sewage | El Paso, TX, U | co | Metagenome | MetaG | USA |

| | | | | | | | | |
|---|---|---|---|---|---|---|---|---|
| 220 | PRJEB27054 | ERR2607578 | sewage | El Paso, TX, U | co | Metagenome | MetaG | USA |
| 220 | PRJEB27054 | ERR2607577 | sewage | El Paso, TX, U | co | Metagenome | MetaG | USA |
| 220 | PRJEB27054 | ERR2607576 | sewage | El Paso, TX, U | co | Metagenome | MetaG | USA |
| 220 | PRJEB27054 | ERR2607575 | sewage | El Paso, TX, U | co | Metagenome | MetaG | USA |
| 220 | PRJEB27054 | ERR2607574 | sewage | El Paso, TX, U | co | Metagenome | MetaG | USA |
| 220 | PRJEB27054 | ERR2607573 | sewage | El Paso, TX, U | co | Metagenome | MetaG | USA |
| 220 | PRJEB27054 | ERR2607572 | sewage | El Paso, TX, U | co | Metagenome | MetaG | USA |
| 220 | PRJEB27054 | ERR2607571 | sewage | El Paso, TX, U | co | Metagenome | MetaG | USA |
| 220 | PRJEB27054 | ERR2607570 | sewage | El Paso, TX, U | co | Metagenome | MetaG | USA |
| 220 | PRJEB27054 | ERR2607569 | sewage | El Paso, TX, U | co | Metagenome | MetaG | USA |
| 220 | PRJEB27054 | ERR2607568 | sewage | El Paso, TX, U | co | Metagenome | MetaG | USA |
| 220 | PRJEB27054 | ERR2607562 | sewage | El Paso, TX, U | co | Metagenome | MetaG | USA |
| 220 | PRJEB27054 | ERR2607563 | sewage | El Paso, TX, U | co | Metagenome | MetaG | USA |
| 222 | PRJEB27054 | ERR2607567 | sewage | Portland, OR, | co | Metagenome | MetaG | USA |
| 222 | PRJEB27054 | ERR2607566 | sewage | Portland, OR, | co | Metagenome | MetaG | USA |
| 222 | PRJEB27054 | ERR2607565 | sewage | Portland, OR, | co | Metagenome | MetaG | USA |
| 222 | PRJEB27054 | ERR2607564 | sewage | Portland, OR, | co | Metagenome | MetaG | USA |
| 223 | PRJEB27054 | ERR2607561 | sewage | Chicago, IL, U | co | Metagenome | MetaG | USA |
| 223 | PRJEB27054 | ERR2607560 | sewage | Chicago, IL, U | co | Metagenome | MetaG | USA |
| 223 | PRJEB27054 | ERR2607559 | sewage | Chicago, IL, U | co | Metagenome | MetaG | USA |
| 223 | PRJEB27054 | ERR2607558 | sewage | Chicago, IL, U | co | Metagenome | MetaG | USA |
| 223 | PRJEB27054 | ERR2607557 | sewage | Chicago, IL, U | co | Metagenome | MetaG | USA |
| 223 | PRJEB27054 | ERR2607556 | sewage | Chicago, IL, U | co | Metagenome | MetaG | USA |
| 223 | PRJEB27054 | ERR2607555 | sewage | Chicago, IL, U | co | Metagenome | MetaG | USA |
| 224 | PRJEB27054 | ERR2607554 | sewage | Seattle, WA, | co | Metagenome | MetaG | USA |
| 224 | PRJEB27054 | ERR2607553 | sewage | Seattle, WA, | co | Metagenome | MetaG | USA |
| 224 | PRJEB27054 | ERR2607552 | sewage | Seattle, WA, | co | Metagenome | MetaG | USA |
| 224 | PRJEB27054 | ERR2607551 | sewage | Seattle, WA, | co | Metagenome | MetaG | USA |
| 225 | PRJEB27054 | ERR2607550 | sewage | Kilimanjaro, T | single | Metagenome | MetaG | Tanzania |
| 226 | PRJEB27054 | ERR2607549 | sewage | Tatlar, Turkey | co | Metagenome | MetaG | Turkey |
| 226 | PRJEB27054 | ERR2607548 | sewage | Tatlar, Turkey | co | Metagenome | MetaG | Turkey |
| 226 | PRJEB27054 | ERR2607547 | sewage | Tatlar, Turkey | co | Metagenome | MetaG | Turkey |
| 227 | PRJEB27054 | ERR2607546 | sewage | Agbadahonou | single | Metagenome | MetaG | Togo |
| 228 | PRJEB27054 | ERR2607545 | sewage | N'Djamena, C | co | Metagenome | MetaG | Chad |
| 228 | PRJEB27054 | ERR2607544 | sewage | N'Djamena, C | co | Metagenome | MetaG | Chad |
| 228 | PRJEB27054 | ERR2607543 | sewage | N'Djamena, C | co | Metagenome | MetaG | Chad |
| 228 | PRJEB27054 | ERR2607542 | sewage | N'Djamena, C | co | Metagenome | MetaG | Chad |
| 228 | PRJEB27054 | ERR2607541 | sewage | N'Djamena, C | co | Metagenome | MetaG | Chad |
| 229 | PRJEB27054 | ERR2607540 | sewage | Uppsala, Swe | single | Metagenome | MetaG | Sweden |
| 230 | PRJEB27054 | ERR2607539 | sewage | Gothenburg, S | co | Metagenome | MetaG | Sweden |
| 230 | PRJEB27054 | ERR2607538 | sewage | Gothenburg, S | co | Metagenome | MetaG | Sweden |
| 230 | PRJEB27054 | ERR2607537 | sewage | Gothenburg, S | co | Metagenome | MetaG | Sweden |
| 231 | PRJEB27054 | ERR2607536 | sewage | Ljubljana, Slo | co | Metagenome | MetaG | Slovenia |
| 231 | PRJEB27054 | ERR2607535 | sewage | Ljubljana, Slo | co | Metagenome | MetaG | Slovenia |
| 231 | PRJEB27054 | ERR2607534 | sewage | Ljubljana, Slo | co | Metagenome | MetaG | Slovenia |
| 231 | PRJEB27054 | ERR2607533 | sewage | Ljubljana, Slo | co | Metagenome | MetaG | Slovenia |
| 232 | PRJEB27054 | ERR2607532 | sewage | Bratislava, Sl | co | Metagenome | MetaG | Slovakia |
| 232 | PRJEB27054 | ERR2607531 | sewage | Bratislava, Sl | co | Metagenome | MetaG | Slovakia |
| 233 | PRJEB27054 | ERR2607530 | sewage | Belgrade, Ser | co | Metagenome | MetaG | Serbia |
| 233 | PRJEB27054 | ERR2607529 | sewage | Belgrade, Ser | co | Metagenome | MetaG | Serbia |
| 233 | PRJEB27054 | ERR2607528 | sewage | Belgrade, Ser | co | Metagenome | MetaG | Serbia |
| 234 | PRJEB27054 | ERR2607527 | sewage | Singapore | co | Metagenome | MetaG | Singapore |
| 234 | PRJEB27054 | ERR2607526 | sewage | Singapore | co | Metagenome | MetaG | Singapore |
| 234 | PRJEB27054 | ERR2607525 | sewage | Singapore | co | Metagenome | MetaG | Singapore |
| 235 | PRJEB27054 | ERR2607524 | sewage | Dakar, Seneg | single | Metagenome | MetaG | Senegal |
| 236 | PRJEB27054 | ERR2607523 | sewage | Pulawy, Polar | co | Metagenome | MetaG | Poland |
| 236 | PRJEB27054 | ERR2607522 | sewage | Pulawy, Polar | co | Metagenome | MetaG | Poland |
| 237 | PRJEB27054 | ERR2607521 | sewage | Pariachi, Peru | single | Metagenome | MetaG | Peru |
| 238 | PRJEB27054 | ERR2607520 | sewage | Jamshed Tow | single | Metagenome | MetaG | Pakistan |
| 239 | PRJEB27054 | ERR2607519 | sewage | Tainui, New Z | single | Metagenome | MetaG | New Zealand |
| 240 | PRJEB27054 | ERR2607518 | sewage | Kathmandu, N | single | Metagenome | MetaG | Nepal |
| 241 | PRJEB27054 | ERR2607517 | sewage | Vollen, Norwa | co | Metagenome | MetaG | Norway |
| 241 | PRJEB27054 | ERR2607516 | sewage | Vollen, Norwa | co | Metagenome | MetaG | Norway |
| 241 | PRJEB27054 | ERR2607515 | sewage | Vollen, Norwa | co | Metagenome | MetaG | Norway |
| 242 | PRJEB27054 | ERR2607514 | sewage | Amsterdam, | co | Metagenome | MetaG | Netherlands |
| 242 | PRJEB27054 | ERR2607513 | sewage | Amsterdam, | co | Metagenome | MetaG | Netherlands |
| 242 | PRJEB27054 | ERR2607512 | sewage | Amsterdam, | co | Metagenome | MetaG | Netherlands |
| 242 | PRJEB27054 | ERR2607511 | sewage | Amsterdam, | co | Metagenome | MetaG | Netherlands |
| 242 | PRJEB27054 | ERR2607510 | sewage | Amsterdam, | co | Metagenome | MetaG | Netherlands |
| 243 | PRJEB27054 | ERR2607509 | sewage | Lagos State, I | co | Metagenome | MetaG | Nigeria |
| 243 | PRJEB27054 | ERR2607508 | sewage | Lagos State, I | co | Metagenome | MetaG | Nigeria |
| 243 | PRJEB27054 | ERR2607507 | sewage | Lagos State, I | co | Metagenome | MetaG | Nigeria |
| 244 | PRJEB27054 | ERR2607506 | sewage | Kuala Lumpur | co | Metagenome | MetaG | Malaysia |
| 244 | PRJEB27054 | ERR2607505 | sewage | Kuala Lumpur | co | Metagenome | MetaG | Malaysia |
| 244 | PRJEB27054 | ERR2607504 | sewage | Kuala Lumpur | co | Metagenome | MetaG | Malaysia |

| | | | | | | | | |
|---|---|---|---|---|---|---|---|---|
| 244 | PRJEB27054 | ERR2607503 | sewage | Kuala Lumpur | co | Metagenome | MetaG | Malaysia |
| 244 | PRJEB27054 | ERR2607502 | sewage | Kuala Lumpur | co | Metagenome | MetaG | Malaysia |
| 245 | PRJEB27054 | ERR2607501 | sewage | Malta | co | Metagenome | MetaG | Malta |
| 245 | PRJEB27054 | ERR2607500 | sewage | Malta | co | Metagenome | MetaG | Malta |
| 245 | PRJEB27054 | ERR2607499 | sewage | Malta | co | Metagenome | MetaG | Malta |
| 246 | PRJEB27054 | ERR2607498 | sewage | Skopje, North | co | Metagenome | MetaG | North Macedonia |
| 246 | PRJEB27054 | ERR2607497 | sewage | Skopje, North | co | Metagenome | MetaG | North Macedonia |
| 246 | PRJEB27054 | ERR2607496 | sewage | Skopje, North | co | Metagenome | MetaG | North Macedonia |
| 246 | PRJEB27054 | ERR2607495 | sewage | Skopje, North | co | Metagenome | MetaG | North Macedonia |
| 247 | PRJEB27054 | ERR2607494 | sewage | Moldova | co | Metagenome | MetaG | Moldova |
| 247 | PRJEB27054 | ERR2607493 | sewage | Moldova | co | Metagenome | MetaG | Moldova |
| 247 | PRJEB27054 | ERR2607492 | sewage | Moldova | co | Metagenome | MetaG | Moldova |
| 248 | PRJEB27054 | ERR2607491 | sewage | Riga, Latia | co | Metagenome | MetaG | Latia |
| 248 | PRJEB27054 | ERR2607490 | sewage | Riga, Latia | co | Metagenome | MetaG | Latia |
| 248 | PRJEB27054 | ERR2607489 | sewage | Riga, Latia | co | Metagenome | MetaG | Latia |
| 248 | PRJEB27054 | ERR2607488 | sewage | Riga, Latia | co | Metagenome | MetaG | Latia |
| 249 | PRJEB27054 | ERR2607487 | sewage | Walferdange, | single | Metagenome | MetaG | Luxembourg |
| 250 | PRJEB27054 | ERR2607486 | sewage | Colombo, Sri | single | Metagenome | MetaG | Sri Lanka |
| 251 | PRJEB27054 | ERR2607485 | sewage | Khan Mean C | single | Metagenome | MetaG | Cambodia |
| 252 | PRJEB27054 | ERR2607484 | sewage | Nairobi, Keny | co | Metagenome | MetaG | Kenya |
| 252 | PRJEB27054 | ERR2607483 | sewage | Nairobi, Keny | co | Metagenome | MetaG | Kenya |
| 252 | PRJEB27054 | ERR2607482 | sewage | Nairobi, Keny | co | Metagenome | MetaG | Kenya |
| 252 | PRJEB27054 | ERR2607481 | sewage | Nairobi, Keny | co | Metagenome | MetaG | Kenya |
| 253 | PRJEB27054 | ERR2607480 | sewage | Almaty, Kaza | single | Metagenome | MetaG | Kazakhstan |
| 254 | PRJEB27054 | ERR2607479 | sewage | Rome, Italy | single | Metagenome | MetaG | Italy |
| 255 | PRJEB27054 | ERR2607478 | sewage | Jerusalem, Is | co | Metagenome | MetaG | Israel |
| 255 | PRJEB27054 | ERR2607477 | sewage | Jerusalem, Is | co | Metagenome | MetaG | Israel |
| 255 | PRJEB27054 | ERR2607476 | sewage | Jerusalem, Is | co | Metagenome | MetaG | Israel |
| 256 | PRJEB27054 | ERR2607475 | sewage | Reykjavik, Ice | co | Metagenome | MetaG | Iceland |
| 256 | PRJEB27054 | ERR2607474 | sewage | Reykjavik, Ice | co | Metagenome | MetaG | Iceland |
| 256 | PRJEB27054 | ERR2607473 | sewage | Reykjavik, Ice | co | Metagenome | MetaG | Iceland |
| 257 | PRJEB27054 | ERR2607472 | sewage | Iran | co | Metagenome | MetaG | Iran |
| 257 | PRJEB27054 | ERR2607471 | sewage | Iran | co | Metagenome | MetaG | Iran |
| 257 | PRJEB27054 | ERR2607470 | sewage | Iran | co | Metagenome | MetaG | Iran |
| 258 | PRJEB27054 | ERR2607469 | sewage | Claddagh, Ire | co | Metagenome | MetaG | Ireland |
| 258 | PRJEB27054 | ERR2607468 | sewage | Claddagh, Ire | co | Metagenome | MetaG | Ireland |
| 258 | PRJEB27054 | ERR2607467 | sewage | Claddagh, Ire | co | Metagenome | MetaG | Ireland |
| 259 | PRJEB27054 | ERR2607466 | sewage | Kerala, India | co | Metagenome | MetaG | India |
| 259 | PRJEB27054 | ERR2607465 | sewage | Kerala, India | co | Metagenome | MetaG | India |
| 259 | PRJEB27054 | ERR2607464 | sewage | Kerala, India | co | Metagenome | MetaG | India |
| 260 | PRJEB27054 | ERR2607463 | sewage | Budapest, Hu | single | Metagenome | MetaG | Hungary |
| 261 | PRJEB27054 | ERR2607462 | sewage | Zagreb, Croat | co | Metagenome | MetaG | Croatia |
| 261 | PRJEB27054 | ERR2607461 | sewage | Zagreb, Croat | co | Metagenome | MetaG | Croatia |
| 261 | PRJEB27054 | ERR2607460 | sewage | Zagreb, Croat | co | Metagenome | MetaG | Croatia |
| 261 | PRJEB27054 | ERR2607459 | sewage | Zagreb, Croat | co | Metagenome | MetaG | Croatia |
| 261 | PRJEB27054 | ERR2607458 | sewage | Zagreb, Croat | co | Metagenome | MetaG | Croatia |
| 262 | PRJEB27054 | ERR2607457 | sewage | Gambia | single | Metagenome | MetaG | Gambia |
| 263 | PRJEB27054 | ERR2607456 | sewage | Tamale, Ghar | single | Metagenome | MetaG | Ghana |
| 264 | PRJEB27054 | ERR2607455 | sewage | Gombori Pass | co | Metagenome | MetaG | Georgia |
| 264 | PRJEB27054 | ERR2607454 | sewage | Gombori Pass | co | Metagenome | MetaG | Georgia |
| 264 | PRJEB27054 | ERR2607453 | sewage | Gombori Pass | co | Metagenome | MetaG | Georgia |
| 264 | PRJEB27054 | ERR2607452 | sewage | Gombori Pass | co | Metagenome | MetaG | Georgia |
| 264 | PRJEB27054 | ERR2607451 | sewage | Gombori Pass | co | Metagenome | MetaG | Georgia |
| 265 | PRJEB27054 | ERR2607450 | sewage | Helsinki, Finla | co | Metagenome | MetaG | Finland |
| 265 | PRJEB27054 | ERR2607449 | sewage | Helsinki, Finla | co | Metagenome | MetaG | Finland |
| 266 | PRJEB27054 | ERR2607448 | sewage | Akaki Kaliti, E | single | Metagenome | MetaG | Ethiopia |
| 267 | PRJEB27054 | ERR2607447 | sewage | Barcelona, Sp | co | Metagenome | MetaG | Spain |
| 267 | PRJEB27054 | ERR2607446 | sewage | Barcelona, Sp | co | Metagenome | MetaG | Spain |
| 267 | PRJEB27054 | ERR2607445 | sewage | Barcelona, Sp | co | Metagenome | MetaG | Spain |
| 267 | PRJEB27054 | ERR2607444 | sewage | Barcelona, Sp | co | Metagenome | MetaG | Spain |
| 267 | PRJEB27054 | ERR2607443 | sewage | Barcelona, Sp | co | Metagenome | MetaG | Spain |
| 267 | PRJEB27054 | ERR2607442 | sewage | Barcelona, Sp | co | Metagenome | MetaG | Spain |
| 268 | PRJEB27054 | ERR2607441 | sewage | Quito, Ecuado | single | Metagenome | MetaG | Ecuador |
| 269 | PRJEB27054 | ERR2607440 | sewage | San Cristobal, | co | Metagenome | MetaG | Ecuador |
| 269 | PRJEB27054 | ERR2607439 | sewage | San Cristobal, | co | Metagenome | MetaG | Ecuador |
| 270 | PRJEB27054 | ERR2607438 | sewage | Copenhagen, | co | Metagenome | MetaG | Denmark |
| 270 | PRJEB27054 | ERR2607437 | sewage | Copenhagen, | co | Metagenome | MetaG | Denmark |
| 270 | PRJEB27054 | ERR2607436 | sewage | Copenhagen, | co | Metagenome | MetaG | Denmark |
| 271 | PRJEB27054 | ERR2607435 | sewage | Werkring, Ge | co | Metagenome | MetaG | Germany |
| 271 | PRJEB27054 | ERR2607434 | sewage | Werkring, Ge | co | Metagenome | MetaG | Germany |
| 271 | PRJEB27054 | ERR2607433 | sewage | Werkring, Ge | co | Metagenome | MetaG | Germany |
| 271 | PRJEB27054 | ERR2607432 | sewage | Werkring, Ge | co | Metagenome | MetaG | Germany |
| 271 | PRJEB27054 | ERR2607431 | sewage | Werkring, Ge | co | Metagenome | MetaG | Germany |
| 272 | PRJEB27054 | ERR2607430 | sewage | Prague, Czech | co | Metagenome | MetaG | Czech Republich |
| 272 | PRJEB27054 | ERR2607429 | sewage | Prague, Czech | co | Metagenome | MetaG | Czech Republich |

| | | | | | | | | |
|---|---|---|---|---|---|---|---|---|
| 272 | PRJEB27054 | ERR2607428 | sewage | Prague, Czech | co | Metagenome | MetaG | Czech Republich |
| 273 | PRJEB27054 | ERR2607427 | sewage | El Charquito, | single | Metagenome | MetaG | Colombia |
| 274 | PRJEB27054 | ERR2607426 | sewage | Cocody, Cote | co | Metagenome | MetaG | Cote d'Ivoire |
| 274 | PRJEB27054 | ERR2607425 | sewage | Cocody, Cote | co | Metagenome | MetaG | Cote d'Ivoire |
| 274 | PRJEB27054 | ERR2607424 | sewage | Cocody, Cote | co | Metagenome | MetaG | Cote d'Ivoire |
| 275 | PRJEB27054 | ERR2607423 | sewage | Guangdong P | single | Metagenome | MetaG | China |
| 276 | PRJEB27054 | ERR2607422 | sewage | Bern, Switzer | co | Metagenome | MetaG | Switzerland |
| 276 | PRJEB27054 | ERR2607421 | sewage | Bern, Switzer | co | Metagenome | MetaG | Switzerland |
| 276 | PRJEB27054 | ERR2607420 | sewage | Bern, Switzer | co | Metagenome | MetaG | Switzerland |
| 276 | PRJEB27054 | ERR2607419 | sewage | Bern, Switzer | co | Metagenome | MetaG | Switzerland |
| 276 | PRJEB27054 | ERR2607418 | sewage | Bern, Switzer | co | Metagenome | MetaG | Switzerland |
| 276 | PRJEB27054 | ERR2607417 | sewage | Bern, Switzer | co | Metagenome | MetaG | Switzerland |
| 276 | PRJEB27054 | ERR2607416 | sewage | Bern, Switzer | co | Metagenome | MetaG | Switzerland |
| 277 | PRJEB27054 | ERR2607415 | sewage | Regina, Cana | single | Metagenome | MetaG | Canada |
| 278 | PRJEB27054 | ERR2607414 | sewage | Ottowa, Cana | single | Metagenome | MetaG | Canada |
| 279 | PRJEB27054 | ERR2607413 | sewage | Toronto, Cana | co | Metagenome | MetaG | Canada |
| 279 | PRJEB27054 | ERR2607412 | sewage | Toronto, Cana | co | Metagenome | MetaG | Canada |
| 279 | PRJEB27054 | ERR2607411 | sewage | Toronto, Cana | co | Metagenome | MetaG | Canada |
| 279 | PRJEB27054 | ERR2607410 | sewage | Toronto, Cana | co | Metagenome | MetaG | Canada |
| 280 | PRJEB27054 | ERR2607409 | sewage | Calgary, Cana | co | Metagenome | MetaG | Canada |
| 280 | PRJEB27054 | ERR2607408 | sewage | Calgary, Cana | co | Metagenome | MetaG | Canada |
| 280 | PRJEB27054 | ERR2607407 | sewage | Calgary, Cana | co | Metagenome | MetaG | Canada |
| 280 | PRJEB27054 | ERR2607406 | sewage | Calgary, Cana | co | Metagenome | MetaG | Canada |
| 280 | PRJEB27054 | ERR2607405 | sewage | Calgary, Cana | co | Metagenome | MetaG | Canada |
| 280 | PRJEB27054 | ERR2607404 | sewage | Calgary, Cana | co | Metagenome | MetaG | Canada |
| 281 | PRJEB27054 | ERR2607403 | sewage | Regina, Cana | co | Metagenome | MetaG | Canada |
| 281 | PRJEB27054 | ERR2607402 | sewage | Regina, Cana | co | Metagenome | MetaG | Canada |
| 282 | PRJEB27054 | ERR2607401 | sewage | Gaborone, Bo | co | Metagenome | MetaG | Botswana |
| 282 | PRJEB27054 | ERR2607400 | sewage | Gaborone, Bo | co | Metagenome | MetaG | Botswana |
| 282 | PRJEB27054 | ERR2607399 | sewage | Gaborone, Bo | co | Metagenome | MetaG | Botswana |
| 282 | PRJEB27054 | ERR2607398 | sewage | Gaborone, Bo | co | Metagenome | MetaG | Botswana |
| 283 | PRJEB27054 | ERR2607397 | sewage | Sabara, Brazi | co | Metagenome | MetaG | Brazil |
| 284 | PRJEB27054 | ERR2607396 | sewage | Belem, Brazil | co | Metagenome | MetaG | Brazil |
| 284 | PRJEB27054 | ERR2607395 | sewage | Belem, Brazil | co | Metagenome | MetaG | Brazil |
| 284 | PRJEB27054 | ERR2607394 | sewage | Belem, Brazil | co | Metagenome | MetaG | Brazil |
| 285 | PRJEB27054 | ERR2607393 | sewage | Sabara, Brazi | co | Metagenome | MetaG | Brazil |
| 285 | PRJEB27054 | ERR2607392 | sewage | Sabara, Brazi | co | Metagenome | MetaG | Brazil |
| 285 | PRJEB27054 | ERR2607391 | sewage | Sabara, Brazi | co | Metagenome | MetaG | Brazil |
| 286 | PRJEB27054 | ERR2607390 | sewage | Kremikovci, B | co | Metagenome | MetaG | Bulgaria |
| 286 | PRJEB27054 | ERR2607389 | sewage | Kremikovci, B | co | Metagenome | MetaG | Bulgaria |
| 286 | PRJEB27054 | ERR2607388 | sewage | Kremikovci, B | co | Metagenome | MetaG | Bulgaria |
| 287 | PRJEB27054 | ERR2607387 | sewage | Vienna, Austr | co | Metagenome | MetaG | Austria |
| 287 | PRJEB27054 | ERR2607386 | sewage | Vienna, Austr | co | Metagenome | MetaG | Austria |
| 287 | PRJEB27054 | ERR2607385 | sewage | Vienna, Austr | co | Metagenome | MetaG | Austria |
| 288 | PRJEB27054 | ERR2607384 | sewage | District of Be | single | Metagenome | MetaG | Australia |
| 289 | PRJEB27054 | ERR2607383 | sewage | Melbourne, A | co | Metagenome | MetaG | Australia |
| 289 | PRJEB27054 | ERR2607382 | sewage | Melbourne, A | co | Metagenome | MetaG | Australia |
| 289 | PRJEB27054 | ERR2607381 | sewage | Melbourne, A | co | Metagenome | MetaG | Australia |
| 289 | PRJEB27054 | ERR2607380 | sewage | Melbourne, A | co | Metagenome | MetaG | Australia |
| 289 | PRJEB27054 | ERR2607379 | sewage | Melbourne, A | co | Metagenome | MetaG | Australia |
| 289 | PRJEB27054 | ERR2607378 | sewage | Melbourne, A | co | Metagenome | MetaG | Australia |
| 290 | PRJEB27054 | ERR2607377 | sewage | District of Be | co | Metagenome | MetaG | Australia |
| 290 | PRJEB27054 | ERR2607376 | sewage | District of Be | co | Metagenome | MetaG | Australia |
| 290 | PRJEB27054 | ERR2607374 | sewage | District of Be | co | Metagenome | MetaG | Australia |
| 291 | PRJEB27054 | ERR2607373 | sewage | Tirana, Alban | co | Metagenome | MetaG | Albania |
| 291 | PRJEB27054 | ERR2607372 | sewage | Tirana, Alban | co | Metagenome | MetaG | Albania |
| 291 | PRJEB27054 | ERR2607371 | sewage | Tirana, Alban | co | Metagenome | MetaG | Albania |
| 208 | PRJNA438281 | SRR6846456 | wwtp influent | Singapore | co | Metavirome | MetaV: WGS | Singapore |
| 208 | PRJNA438281 | SRR6846470 | wwtp influent | Singapore | co | Metavirome | MetaV: WGS | Singapore |
| 208 | PRJNA438281 | SRR6846454 | wwtp influent | Singapore | co | Metavirome | MetaV: WGS | Singapore |
| 208 | PRJNA438281 | SRR6846460 | wwtp influent | Singapore | co | Metavirome | MetaV: WGS | Singapore |
| 212 | PRJNA526679 | SRR8715492 | hospital wastewater | Tel Aviv, Isra | co | Metavirome | MetaV: WGS | Israel |
| 212 | PRJNA526679 | SRR8715494 | hospital wastewater | Tel Aviv, Isra | co | Metavirome | MetaV: WGS | Israel |
| 212 | PRJNA526679 | SRR8715495 | hospital wastewater | Tel Aviv, Isra | co | Metavirome | MetaV: WGS | Israel |
| 292 | PRJEB47975 | ERR7015403 | hospital wastewater | Finland | co | Metagenome | MetaG | Finland |
| 292 | PRJEB47975 | ERR7015402 | hospital wastewater | Finland | co | Metagenome | MetaG | Finland |
| 292 | PRJEB47975 | ERR7015401 | hospital wastewater | Finland | co | Metagenome | MetaG | Finland |
| 292 | PRJEB47975 | ERR7015400 | hospital wastewater | Finland | co | Metagenome | MetaG | Finland |
| 292 | PRJEB47975 | ERR7015399 | hospital wastewater | Finland | co | Metagenome | MetaG | Finland |
| 292 | PRJEB47975 | ERR7015398 | hospital wastewater | Finland | co | Metagenome | MetaG | Finland |
| 292 | PRJEB47975 | ERR7015397 | hospital wastewater | Finland | co | Metagenome | MetaG | Finland |
| 292 | PRJEB47975 | ERR7015396 | hospital wastewater | Finland | co | Metagenome | MetaG | Finland |
| 292 | PRJEB47975 | ERR7015395 | hospital wastewater | Finland | co | Metagenome | MetaG | Finland |
| 293 | PRJEB47975 | ERR7015394 | hospital septic tank | Burkina Faso, | co | Metagenome | MetaG | Africa |
| 293 | PRJEB47975 | ERR7015393 | hospital septic tank | Burkina Faso, | co | Metagenome | MetaG | Africa |

| | | | | | | | | |
|---|---|---|---|---|---|---|---|---|
| 293 | PRJEB47975 | ERR7015392 | hospital septic tank | Burkina Faso, | co | Metagenome | MetaG | Africa |
| 293 | PRJEB47975 | ERR7015391 | hospital septic tank | Burkina Faso, | co | Metagenome | MetaG | Africa |
| 293 | PRJEB47975 | ERR7015390 | hospital septic tank | Burkina Faso, | co | Metagenome | MetaG | Africa |
| 293 | PRJEB47975 | ERR7015389 | hospital septic tank | Burkina Faso, | co | Metagenome | MetaG | Africa |
| 293 | PRJEB47975 | ERR7015388 | hospital septic tank | Burkina Faso, | co | Metagenome | MetaG | Africa |
| 293 | PRJEB47975 | ERR7015387 | hospital septic tank | Burkina Faso, | co | Metagenome | MetaG | Africa |
| 293 | PRJEB47975 | ERR7015386 | hospital septic tank | Burkina Faso, | co | Metagenome | MetaG | Africa |
| 293 | PRJEB47975 | ERR7015385 | hospital septic tank | Burkina Faso, | co | Metagenome | MetaG | Africa |
| 293 | PRJEB47975 | ERR7015384 | hospital septic tank | Burkina Faso, | co | Metagenome | MetaG | Africa |
| 293 | PRJEB47975 | ERR7015383 | hospital septic tank | Burkina Faso, | co | Metagenome | MetaG | Africa |
| 293 | PRJEB47975 | ERR7015382 | hospital septic tank | Burkina Faso, | co | Metagenome | MetaG | Africa |
| 293 | PRJEB47975 | ERR7015381 | hospital septic tank | Burkina Faso, | co | Metagenome | MetaG | Africa |
| 293 | PRJEB47975 | ERR7015380 | hospital septic tank | Burkina Faso, | co | Metagenome | MetaG | Africa |
| 293 | PRJEB47975 | ERR7015379 | hospital septic tank | Burkina Faso, | co | Metagenome | MetaG | Africa |
| 293 | PRJEB47975 | ERR7015378 | hospital septic tank | Burkina Faso, | co | Metagenome | MetaG | Africa |
| 293 | PRJEB47975 | ERR7015377 | hospital septic tank | Burkina Faso, | co | Metagenome | MetaG | Africa |
| 293 | PRJEB47975 | ERR7015376 | hospital septic tank | Burkina Faso, | co | Metagenome | MetaG | Africa |
| 293 | PRJEB47975 | ERR7015375 | hospital septic tank | Burkina Faso, | co | Metagenome | MetaG | Africa |
| 293 | PRJEB47975 | ERR7015374 | hospital septic tank | Burkina Faso, | co | Metagenome | MetaG | Africa |
| 293 | PRJEB47975 | ERR7015373 | hospital septic tank | Burkina Faso, | co | Metagenome | MetaG | Africa |
| 293 | PRJEB47975 | ERR7015372 | hospital septic tank | Burkina Faso, | co | Metagenome | MetaG | Africa |
| 293 | PRJEB47975 | ERR7015371 | hospital septic tank | Burkina Faso, | co | Metagenome | MetaG | Africa |
| 293 | PRJEB47975 | ERR7015370 | hospital septic tank | Burkina Faso, | co | Metagenome | MetaG | Africa |
| 293 | PRJEB47975 | ERR7015369 | hospital septic tank | Burkina Faso, | co | Metagenome | MetaG | Africa |
| 293 | PRJEB47975 | ERR7015368 | hospital septic tank | Burkina Faso, | co | Metagenome | MetaG | Africa |
| 293 | PRJEB47975 | ERR7015367 | hospital septic tank | Burkina Faso, | co | Metagenome | MetaG | Africa |
| 293 | PRJEB47975 | ERR7015366 | hospital septic tank | Burkina Faso, | co | Metagenome | MetaG | Africa |
| 293 | PRJEB47975 | ERR7015365 | hospital septic tank | Burkina Faso, | co | Metagenome | MetaG | Africa |
| 293 | PRJEB47975 | ERR7015364 | hospital septic tank | Burkina Faso, | co | Metagenome | MetaG | Africa |
| 293 | PRJEB47975 | ERR7015363 | hospital septic tank | Burkina Faso, | co | Metagenome | MetaG | Africa |
| 293 | PRJEB47975 | ERR7015362 | hospital septic tank | Burkina Faso, | co | Metagenome | MetaG | Africa |
| 293 | PRJEB47975 | ERR7015361 | hospital septic tank | Burkina Faso, | co | Metagenome | MetaG | Africa |
| 293 | PRJEB47975 | ERR7015360 | hospital septic tank | Burkina Faso, | co | Metagenome | MetaG | Africa |
| 293 | PRJEB47975 | ERR7015359 | hospital septic tank | Burkina Faso, | co | Metagenome | MetaG | Africa |
| 293 | PRJEB47975 | ERR7015358 | hospital septic tank | Burkina Faso, | co | Metagenome | MetaG | Africa |
| 293 | PRJEB47975 | ERR7015357 | hospital septic tank | Burkina Faso, | co | Metagenome | MetaG | Africa |
| 294 | PRJEB47975 | ERR7015351 | hospital septic tank | Benin | co | Metagenome | MetaG | Benin |
| 294 | PRJEB47975 | ERR7015350 | hospital septic tank | Benin | co | Metagenome | MetaG | Benin |
| 294 | PRJEB47975 | ERR7015348 | hospital septic tank | Benin | co | Metagenome | MetaG | Benin |
| 294 | PRJEB47975 | ERR7015347 | hospital septic tank | Benin | co | Metagenome | MetaG | Benin |
| 294 | PRJEB47975 | ERR7015346 | hospital septic tank | Benin | co | Metagenome | MetaG | Benin |
| 294 | PRJEB47975 | ERR7015345 | hospital septic tank | Benin | co | Metagenome | MetaG | Benin |
| 294 | PRJEB47975 | ERR7015344 | hospital septic tank | Benin | co | Metagenome | MetaG | Benin |
| 294 | PRJEB47975 | ERR7015343 | hospital septic tank | Benin | co | Metagenome | MetaG | Benin |
| 294 | PRJEB47975 | ERR7015342 | hospital septic tank | Benin | co | Metagenome | MetaG | Benin |
| 294 | PRJEB47975 | ERR7015340 | hospital septic tank | Benin | co | Metagenome | MetaG | Benin |
| 294 | PRJEB47975 | ERR7015339 | hospital septic tank | Benin | co | Metagenome | MetaG | Benin |
| 294 | PRJEB47975 | ERR7015338 | hospital septic tank | Benin | co | Metagenome | MetaG | Benin |
| 294 | PRJEB47975 | ERR7015337 | hospital septic tank | Benin | co | Metagenome | MetaG | Benin |
| 294 | PRJEB47975 | ERR7015335 | hospital septic tank | Benin | co | Metagenome | MetaG | Benin |
| 294 | PRJEB47975 | ERR7015334 | hospital septic tank | Benin | co | Metagenome | MetaG | Benin |
| 294 | PRJEB47975 | ERR7015333 | hospital septic tank | Benin | co | Metagenome | MetaG | Benin |
| 294 | PRJEB47975 | ERR7015332 | hospital septic tank | Benin | co | Metagenome | MetaG | Benin |
| 294 | PRJEB47975 | ERR7015330 | hospital septic tank | Benin | co | Metagenome | MetaG | Benin |
| 294 | PRJEB47975 | ERR7015329 | hospital septic tank | Benin | co | Metagenome | MetaG | Benin |
| 294 | PRJEB47975 | ERR7015328 | hospital septic tank | Benin | co | Metagenome | MetaG | Benin |
| 294 | PRJEB47975 | ERR7015327 | hospital septic tank | Benin | co | Metagenome | MetaG | Benin |
| 294 | PRJEB47975 | ERR7015321 | hospital septic tank | Benin | co | Metagenome | MetaG | Benin |
| 294 | PRJEB47975 | ERR7015320 | hospital septic tank | Benin | co | Metagenome | MetaG | Benin |
| 294 | PRJEB47975 | ERR7015318 | hospital septic tank | Benin | co | Metagenome | MetaG | Benin |
| 294 | PRJEB47975 | ERR7015317 | hospital septic tank | Benin | co | Metagenome | MetaG | Benin |
| 294 | PRJEB47975 | ERR7015316 | hospital septic tank | Benin | co | Metagenome | MetaG | Benin |
| 294 | PRJEB47975 | ERR7015315 | hospital septic tank | Benin | co | Metagenome | MetaG | Benin |
| 294 | PRJEB47975 | ERR7015313 | hospital septic tank | Benin | co | Metagenome | MetaG | Benin |
| 294 | PRJEB47975 | ERR7015312 | hospital septic tank | Benin | co | Metagenome | MetaG | Benin |
| 294 | PRJEB47975 | ERR7015311 | hospital septic tank | Benin | co | Metagenome | MetaG | Benin |
| 295 | PRJEB52722 | ERR9723913 | wwtp influent | United Kingd | co | Metagenome | MetaG | United Kingdom |
| 295 | PRJEB52722 | ERR9723914 | wwtp influent | United Kingd | co | Metagenome | MetaG | United Kingdom |
| 295 | PRJEB52722 | ERR9723915 | wwtp influent | United Kingd | co | Metagenome | MetaG | United Kingdom |
| 295 | PRJEB52722 | ERR9723916 | wwtp influent | United Kingd | co | Metagenome | MetaG | United Kingdom |
| 295 | PRJEB52722 | ERR9723917 | wwtp influent | United Kingd | co | Metagenome | MetaG | United Kingdom |
| 295 | PRJEB52722 | ERR9723918 | wwtp influent | United Kingd | co | Metagenome | MetaG | United Kingdom |
| 295 | PRJEB52722 | ERR9723919 | wwtp influent | United Kingd | co | Metagenome | MetaG | United Kingdom |
| 296 | PRJNA857474 | SRR20074303 | wwtp influent | China | single | Metagenome | MetaG | China |
| 297 | PRJNA768945 | SRR16214424 | raw sewage | Manitoba, Ca | co | Metagenome | MetaG | Canada |

| | | | | | | | | |
|---|---|---|---|---|---|---|---|---|
| 297 | PRJNA768945 | SRR16214422 | raw sewage | Manitoba, Ca | co | Metagenome | MetaG | Canada |
| 297 | PRJNA768945 | SRR16214421 | raw sewage | Manitoba, Ca | co | Metagenome | MetaG | Canada |
| 297 | PRJNA768945 | SRR16214420 | raw sewage | Manitoba, Ca | co | Metagenome | MetaG | Canada |
| 298 | PRJNA768945 | SRR16214419 | raw sewage | Manitoba, Ca | co | Metavirome | MetaV: WGS | Canada |
| 298 | PRJNA768945 | SRR16214418 | raw sewage | Manitoba, Ca | co | Metavirome | MetaV: WGS | Canada |
| 298 | PRJNA768945 | SRR16214417 | raw sewage | Manitoba, Ca | co | Metavirome | MetaV: WGS | Canada |
| 298 | PRJNA768945 | SRR16214416 | raw sewage | Manitoba, Ca | co | Metavirome | MetaV: WGS | Canada |
| 299 | PRJNA749165 | SRR15237802 | wwtp influent | HeFei, China | single | Metagenome | MetaG | China |
| 300 | PRJNA749165 | SRR15237812 | wwtp influent | BaoTou, China | single | Metagenome | MetaG | China |
| 301 | PRJNA749165 | SRR15237818 | wwtp influent | TongLu, China | single | Metagenome | MetaG | China |
| 302 | PRJNA737178 | SRR14800556 | wwtp influent | Xinjiang, Chin | co | Metagenome | MetaG | China |
| 302 | PRJNA737178 | SRR14800553 | wwtp influent | Xinjiang, Chin | co | Metagenome | MetaG | China |
| 302 | PRJNA737178 | SRR14800550 | wwtp influent | Xinjiang, Chin | co | Metagenome | MetaG | China |
| 303 | PRJEB38014 | ERR4450760 | wwtp influent | Jaipur, India | co | Metagenome | MetaG | India |
| 303 | PRJEB38014 | ERR4450761 | wwtp influent | Jaipur, India | co | Metagenome | MetaG | India |
| 303 | PRJEB38014 | ERR4450762 | wwtp influent | Jaipur, India | co | Metagenome | MetaG | India |
| 303 | PRJEB38014 | ERR4450754 | wwtp influent | Jaipur, India | co | Metagenome | MetaG | India |
| 303 | PRJEB38014 | ERR4450753 | wwtp influent | Jaipur, India | co | Metagenome | MetaG | India |
| 303 | PRJEB38014 | ERR4450752 | wwtp influent | Jaipur, India | co | Metagenome | MetaG | India |
| 303 | PRJEB38014 | ERR4450745 | wwtp influent | Jaipur, India | co | Metagenome | MetaG | India |
| 303 | PRJEB38014 | ERR4450744 | wwtp influent | Jaipur, India | co | Metagenome | MetaG | India |
| 303 | PRJEB38014 | ERR4450743 | wwtp influent | Jaipur, India | co | Metagenome | MetaG | India |
| 303 | PRJEB38014 | ERR4450736 | wwtp influent | Jaipur, India | co | Metagenome | MetaG | India |
| 303 | PRJEB38014 | ERR4450735 | wwtp influent | Jaipur, India | co | Metagenome | MetaG | India |
| 304 | PRJNA730932 | SRR14597296 | farm ww | Donggang, Ch | co | Metagenome | MetaG | China |
| 304 | PRJNA730932 | SRR14597297 | farm ww | Donggang, Ch | co | Metagenome | MetaG | China |
| 304 | PRJNA730932 | SRR14597298 | farm ww | Donggang, Ch | co | Metagenome | MetaG | China |
| 304 | PRJNA730932 | SRR14597299 | farm ww | Donggang, Ch | co | Metagenome | MetaG | China |
| 304 | PRJNA730932 | SRR14597300 | farm ww | Donggang, Ch | co | Metagenome | MetaG | China |
| 304 | PRJNA730932 | SRR14597301 | farm ww | Donggang, Ch | co | Metagenome | MetaG | China |
| 304 | PRJNA730932 | SRR14597279 | farm ww | Donggang, Ch | co | Metagenome | MetaG | China |
| 304 | PRJNA730932 | SRR14597278 | farm ww | Donggang, Ch | co | Metagenome | MetaG | China |
| 305 | PRJNA730932 | SRR14597282 | urban ww | Donggang, Ch | co | Metagenome | MetaG | China |
| 305 | PRJNA730932 | SRR14597281 | urban ww | Donggang, Ch | co | Metagenome | MetaG | China |
| 305 | PRJNA730932 | SRR14597280 | urban ww | Donggang, Ch | co | Metagenome | MetaG | China |
| 305 | PRJNA730932 | SRR14597277 | urban ww | Donggang, Ch | co | Metagenome | MetaG | China |
| 305 | PRJNA730932 | SRR14597276 | urban ww | Donggang, Ch | co | Metagenome | MetaG | China |
| 305 | PRJNA730932 | SRR14597291 | urban ww | Donggang, Ch | co | Metagenome | MetaG | China |
| 305 | PRJNA730932 | SRR14597302 | urban ww | Donggang, Ch | co | Metagenome | MetaG | China |
| 305 | PRJNA730932 | SRR14597303 | urban ww | Donggang, Ch | co | Metagenome | MetaG | China |
| 306 | PRJNA730932 | SRR14597283 | farm ww | Zhanjiang, Ch | co | Metagenome | MetaG | China |
| 306 | PRJNA730932 | SRR14597284 | farm ww | Zhanjiang, Ch | co | Metagenome | MetaG | China |
| 306 | PRJNA730932 | SRR14597285 | farm ww | Zhanjiang, Ch | co | Metagenome | MetaG | China |
| 306 | PRJNA730932 | SRR14597286 | farm ww | Zhanjiang, Ch | co | Metagenome | MetaG | China |
| 306 | PRJNA730932 | SRR14597287 | farm ww | Zhanjiang, Ch | co | Metagenome | MetaG | China |
| 306 | PRJNA730932 | SRR14597288 | farm ww | Zhanjiang, Ch | co | Metagenome | MetaG | China |
| 307 | PRJNA730932 | SRR14597289 | urban ww | Zhanjiang, Ch | co | Metagenome | MetaG | China |
| 307 | PRJNA730932 | SRR14597290 | urban ww | Zhanjiang, Ch | co | Metagenome | MetaG | China |
| 307 | PRJNA730932 | SRR14597292 | urban ww | Zhanjiang, Ch | co | Metagenome | MetaG | China |
| 307 | PRJNA730932 | SRR14597293 | urban ww | Zhanjiang, Ch | co | Metagenome | MetaG | China |
| 307 | PRJNA730932 | SRR14597294 | urban ww | Zhanjiang, Ch | co | Metagenome | MetaG | China |
| 307 | PRJNA730932 | SRR14597295 | urban ww | Zhanjiang, Ch | co | Metagenome | MetaG | China |
| 308 | PRJEB41538 | ERR4878532 | household ww | Kleve, Germa | co | Metagenome | MetaG | Germany |
| 308 | PRJEB41538 | ERR4878531 | household ww | Kleve, Germa | co | Metagenome | MetaG | Germany |
| 308 | PRJEB41538 | ERR4878530 | household ww | Kleve, Germa | co | Metagenome | MetaG | Germany |
| 308 | PRJEB41538 | ERR4878529 | household ww | Kleve, Germa | co | Metagenome | MetaG | Germany |
| 308 | PRJEB41538 | ERR4878528 | household ww | Kleve, Germa | co | Metagenome | MetaG | Germany |
| 308 | PRJEB41538 | ERR4878527 | household ww | Kleve, Germa | co | Metagenome | MetaG | Germany |
| 309 | PRJEB41538 | ERR4878517 | wwtp influent | Kleve, Germa | co | Metagenome | MetaG | Germany |
| 309 | PRJEB41538 | ERR4878516 | wwtp influent | Kleve, Germa | co | Metagenome | MetaG | Germany |
| 309 | PRJEB41538 | ERR4878515 | wwtp influent | Kleve, Germa | co | Metagenome | MetaG | Germany |
| 309 | PRJEB41538 | ERR4878514 | wwtp influent | Kleve, Germa | co | Metagenome | MetaG | Germany |
| 310 | PRJEB18607 | ERR2041951 | sewage | Orsay, France | single | Metagenome | MetaG | France |
| 311 | PRJEB15519 | ERR1661349 | wwtp influent | Thuwal, Saud | co | Metavirome | MetaV: WGS | Saudi Arabia |
| 311 | PRJEB15519 | ERR1661348 | wwtp influent | Thuwal, Saud | co | Metavirome | MetaV: WGS | Saudi Arabia |
| 311 | PRJEB15519 | ERR1661343 | wwtp influent | Thuwal, Saud | co | Metavirome | MetaV: WGS | Saudi Arabia |
| 311 | PRJEB15519 | ERR1661342 | wwtp influent | Thuwal, Saud | co | Metavirome | MetaV: WGS | Saudi Arabia |
| 312 | PRJEB15084 | ERR1560011 | hospital SH ww | Saudi Arabia | co | Metagenome | MetaG | Saudi Arabia |
| 312 | PRJEB15084 | ERR1560010 | hospital SH ww | Saudi Arabia | co | Metagenome | MetaG | Saudi Arabia |
| 312 | PRJEB15084 | ERR1560009 | hospital SH ww | Saudi Arabia | co | Metagenome | MetaG | Saudi Arabia |
| 312 | PRJEB15084 | ERR1560008 | hospital SH ww | Saudi Arabia | co | Metagenome | MetaG | Saudi Arabia |
| 312 | PRJEB15084 | ERR1560007 | hospital SH ww | Saudi Arabia | co | Metagenome | MetaG | Saudi Arabia |
| 312 | PRJEB15084 | ERR1560006 | hospital SH ww | Saudi Arabia | co | Metagenome | MetaG | Saudi Arabia |
| 313 | PRJEB15084 | ERR1559999 | hospital IH ww | Saudi Arabia | co | Metagenome | MetaG | Saudi Arabia |
| 313 | PRJEB15084 | ERR1559995 | hospital IH ww | Saudi Arabia | co | Metagenome | MetaG | Saudi Arabia |

| | | | | | | | | |
|---|---|---|---|---|---|---|---|---|
| 313 | PRJEB15084 | ERR1559996 | hospital IH ww | Saudi Arabia | co | Metagenome | MetaG | Saudi Arabia |
| 313 | PRJEB15084 | ERR1559997 | hospital IH ww | Saudi Arabia | co | Metagenome | MetaG | Saudi Arabia |
| 313 | PRJEB15084 | ERR1559998 | hospital IH ww | Saudi Arabia | co | Metagenome | MetaG | Saudi Arabia |
| 209 | PRJEB30546 | ERR3026572 | sewage | Global | co | Metavirome | MetaV: WGS | Global |
| 209 | PRJEB30546 | ERR3026571 | sewage | Global | co | Metavirome | MetaV: WGS | Global |
| 209 | PRJEB30546 | ERR3026563 | sewage | Global | co | Metavirome | MetaV: WGS | Global |
| 209 | PRJEB30546 | ERR3026569 | sewage | Global | co | Metavirome | MetaV: WGS | Global |
| 209 | PRJEB30546 | ERR3026575 | sewage | Global | co | Metavirome | MetaV: WGS | Global |
| 209 | PRJEB30546 | ERR3026576 | sewage | Global | co | Metavirome | MetaV: WGS | Global |
| 209 | PRJEB30546 | ERR3026570 | sewage | Global | co | Metavirome | MetaV: WGS | Global |
| 209 | PRJEB30546 | ERR3026564 | sewage | Global | co | Metavirome | MetaV: WGS | Global |
| 209 | PRJEB30546 | ERR3026566 | sewage | Global | co | Metavirome | MetaV: WGS | Global |
| 209 | PRJEB30546 | ERR3026554 | sewage | Global | co | Metavirome | MetaV: WGS | Global |
| 209 | PRJEB30546 | ERR3026553 | sewage | Global | co | Metavirome | MetaV: WGS | Global |
| 209 | PRJEB30546 | ERR3026565 | sewage | Global | co | Metavirome | MetaV: WGS | Global |
| 209 | PRJEB30546 | ERR3026567 | sewage | Global | co | Metavirome | MetaV: WGS | Global |
| 209 | PRJEB30546 | ERR3026568 | sewage | Global | co | Metavirome | MetaV: WGS | Global |
| 209 | PRJEB30546 | ERR3026543 | sewage | Global | co | Metavirome | MetaV: WGS | Global |
| 209 | PRJEB30546 | ERR3026515 | sewage | Global | co | Metavirome | MetaV: WGS | Global |
| 209 | PRJEB30546 | ERR3026509 | sewage | Global | co | Metavirome | MetaV: WGS | Global |
| 209 | PRJEB30546 | ERR3026510 | sewage | Global | co | Metavirome | MetaV: WGS | Global |
| 209 | PRJEB30546 | ERR3026516 | sewage | Global | co | Metavirome | MetaV: WGS | Global |
| 209 | PRJEB30546 | ERR3026544 | sewage | Global | co | Metavirome | MetaV: WGS | Global |
| 209 | PRJEB30546 | ERR3026512 | sewage | Global | co | Metavirome | MetaV: WGS | Global |
| 209 | PRJEB30546 | ERR3026518 | sewage | Global | co | Metavirome | MetaV: WGS | Global |
| 209 | PRJEB30546 | ERR3026503 | sewage | Global | co | Metavirome | MetaV: WGS | Global |
| 209 | PRJEB30546 | ERR3026504 | sewage | Global | co | Metavirome | MetaV: WGS | Global |
| 209 | PRJEB30546 | ERR3026517 | sewage | Global | co | Metavirome | MetaV: WGS | Global |
| 209 | PRJEB30546 | ERR3026511 | sewage | Global | co | Metavirome | MetaV: WGS | Global |
| 209 | PRJEB30546 | ERR3026507 | sewage | Global | co | Metavirome | MetaV: WGS | Global |
| 209 | PRJEB30546 | ERR3026521 | sewage | Global | co | Metavirome | MetaV: WGS | Global |
| 209 | PRJEB30546 | ERR3026494 | sewage | Global | co | Metavirome | MetaV: WGS | Global |
| 209 | PRJEB30546 | ERR3026493 | sewage | Global | co | Metavirome | MetaV: WGS | Global |
| 209 | PRJEB30546 | ERR3026522 | sewage | Global | co | Metavirome | MetaV: WGS | Global |
| 209 | PRJEB30546 | ERR3026508 | sewage | Global | co | Metavirome | MetaV: WGS | Global |
| 209 | PRJEB30546 | ERR3026520 | sewage | Global | co | Metavirome | MetaV: WGS | Global |
| 209 | PRJEB30546 | ERR3026506 | sewage | Global | co | Metavirome | MetaV: WGS | Global |
| 209 | PRJEB30546 | ERR3026534 | sewage | Global | co | Metavirome | MetaV: WGS | Global |
| 209 | PRJEB30546 | ERR3026533 | sewage | Global | co | Metavirome | MetaV: WGS | Global |
| 209 | PRJEB30546 | ERR3026505 | sewage | Global | co | Metavirome | MetaV: WGS | Global |
| 209 | PRJEB30546 | ERR3026519 | sewage | Global | co | Metavirome | MetaV: WGS | Global |
| 209 | PRJEB30546 | ERR3026524 | sewage | Global | co | Metavirome | MetaV: WGS | Global |
| 209 | PRJEB30546 | ERR3026536 | sewage | Global | co | Metavirome | MetaV: WGS | Global |
| 209 | PRJEB30546 | ERR3026530 | sewage | Global | co | Metavirome | MetaV: WGS | Global |
| 209 | PRJEB30546 | ERR3026497 | sewage | Global | co | Metavirome | MetaV: WGS | Global |
| 209 | PRJEB30546 | ERR3026498 | sewage | Global | co | Metavirome | MetaV: WGS | Global |
| 209 | PRJEB30546 | ERR3026492 | sewage | Global | co | Metavirome | MetaV: WGS | Global |
| 209 | PRJEB30546 | ERR3026529 | sewage | Global | co | Metavirome | MetaV: WGS | Global |
| 209 | PRJEB30546 | ERR3026535 | sewage | Global | co | Metavirome | MetaV: WGS | Global |
| 209 | PRJEB30546 | ERR3026523 | sewage | Global | co | Metavirome | MetaV: WGS | Global |
| 209 | PRJEB30546 | ERR3026531 | sewage | Global | co | Metavirome | MetaV: WGS | Global |
| 209 | PRJEB30546 | ERR3026537 | sewage | Global | co | Metavirome | MetaV: WGS | Global |
| 209 | PRJEB30546 | ERR3026496 | sewage | Global | co | Metavirome | MetaV: WGS | Global |
| 209 | PRJEB30546 | ERR3026495 | sewage | Global | co | Metavirome | MetaV: WGS | Global |
| 209 | PRJEB30546 | ERR3026538 | sewage | Global | co | Metavirome | MetaV: WGS | Global |
| 209 | PRJEB30546 | ERR3026532 | sewage | Global | co | Metavirome | MetaV: WGS | Global |
| 209 | PRJEB30546 | ERR3026528 | sewage | Global | co | Metavirome | MetaV: WGS | Global |
| 209 | PRJEB30546 | ERR3026542 | sewage | Global | co | Metavirome | MetaV: WGS | Global |
| 209 | PRJEB30546 | ERR3026499 | sewage | Global | co | Metavirome | MetaV: WGS | Global |
| 209 | PRJEB30546 | ERR3026500 | sewage | Global | co | Metavirome | MetaV: WGS | Global |
| 209 | PRJEB30546 | ERR3026541 | sewage | Global | co | Metavirome | MetaV: WGS | Global |
| 209 | PRJEB30546 | ERR3026527 | sewage | Global | co | Metavirome | MetaV: WGS | Global |
| 209 | PRJEB30546 | ERR3026539 | sewage | Global | co | Metavirome | MetaV: WGS | Global |
| 209 | PRJEB30546 | ERR3026525 | sewage | Global | co | Metavirome | MetaV: WGS | Global |
| 209 | PRJEB30546 | ERR3026513 | sewage | Global | co | Metavirome | MetaV: WGS | Global |
| 209 | PRJEB30546 | ERR3026502 | sewage | Global | co | Metavirome | MetaV: WGS | Global |
| 209 | PRJEB30546 | ERR3026501 | sewage | Global | co | Metavirome | MetaV: WGS | Global |
| 209 | PRJEB30546 | ERR3026514 | sewage | Global | co | Metavirome | MetaV: WGS | Global |
| 209 | PRJEB30546 | ERR3026526 | sewage | Global | co | Metavirome | MetaV: WGS | Global |
| 209 | PRJEB30546 | ERR3026540 | sewage | Global | co | Metavirome | MetaV: WGS | Global |
| 209 | PRJEB30546 | ERR3026557 | sewage | Global | co | Metavirome | MetaV: WGS | Global |
| 209 | PRJEB30546 | ERR3026551 | sewage | Global | co | Metavirome | MetaV: WGS | Global |
| 209 | PRJEB30546 | ERR3026552 | sewage | Global | co | Metavirome | MetaV: WGS | Global |
| 209 | PRJEB30546 | ERR3026558 | sewage | Global | co | Metavirome | MetaV: WGS | Global |
| 209 | PRJEB30546 | ERR3026550 | sewage | Global | co | Metavirome | MetaV: WGS | Global |

| | | | | | | | | |
|---|---|---|---|---|---|---|---|---|
| 209 | PRJEB30546 | ERR3026556 | sewage | Global | co | Metavirome | MetaV: WGS | Global |
| 209 | PRJEB30546 | ERR3026555 | sewage | Global | co | Metavirome | MetaV: WGS | Global |
| 209 | PRJEB30546 | ERR3026549 | sewage | Global | co | Metavirome | MetaV: WGS | Global |
| 209 | PRJEB30546 | ERR3026545 | sewage | Global | co | Metavirome | MetaV: WGS | Global |
| 209 | PRJEB30546 | ERR3026559 | sewage | Global | co | Metavirome | MetaV: WGS | Global |
| 209 | PRJEB30546 | ERR3026573 | sewage | Global | co | Metavirome | MetaV: WGS | Global |
| 209 | PRJEB30546 | ERR3026574 | sewage | Global | co | Metavirome | MetaV: WGS | Global |
| 209 | PRJEB30546 | ERR3026560 | sewage | Global | co | Metavirome | MetaV: WGS | Global |
| 209 | PRJEB30546 | ERR3026546 | sewage | Global | co | Metavirome | MetaV: WGS | Global |
| 209 | PRJEB30546 | ERR3026562 | sewage | Global | co | Metavirome | MetaV: WGS | Global |
| 209 | PRJEB30546 | ERR3026548 | sewage | Global | co | Metavirome | MetaV: WGS | Global |
| 209 | PRJEB30546 | ERR3026547 | sewage | Global | co | Metavirome | MetaV: WGS | Global |
| 209 | PRJEB30546 | ERR3026561 | sewage | Global | co | Metavirome | MetaV: WGS | Global |
| 67 | PRJEB13831 | ERR9855099 | sewage | China | single | Metagenome | MetaG | China |
| 68 | PRJEB13831 | ERR9855098 | sewage | Malaysia | single | Metagenome | MetaG | Malaysia |
| 69 | PRJEB13831 | ERR9855097 | sewage | Brazil | single | Metagenome | MetaG | Brazil |
| 70 | PRJEB13831 | ERR9855096 | sewage | Tanzania | single | Metagenome | MetaG | Tanzania |
| 71 | PRJEB13831 | ERR9855095 | sewage | Iran | single | Metagenome | MetaG | Iran |
| 72 | PRJEB13831 | ERR9855094 | sewage | Victoria, Aust | single | Metagenome | MetaG | Australia |
| 73 | PRJEB13831 | ERR9855093 | sewage | Iran | single | Metagenome | MetaG | Iran |
| 74 | PRJEB13831 | ERR9855092 | sewage | Germany | single | Metagenome | MetaG | Germany |
| 75 | PRJEB13831 | ERR9855091 | sewage | Brazil | single | Metagenome | MetaG | Brazil |
| 76 | PRJEB13831 | ERR9855090 | sewage | Tanzania | single | Metagenome | MetaG | Tanzania |
| 77 | PRJEB13831 | ERR9855089 | sewage | Germany | single | Metagenome | MetaG | Germany |
| 78 | PRJEB13831 | ERR9855088 | sewage | Canada | single | Metagenome | MetaG | Canada |
| 79 | PRJEB13831 | ERR9855087 | sewage | Canada | single | Metagenome | MetaG | Canada |
| 80 | PRJEB13831 | ERR9855086 | sewage | Victoria, Aust | single | Metagenome | MetaG | Australia |
| 81 | PRJEB13831 | ERR9855085 | sewage | Malaysia | single | Metagenome | MetaG | Malaysia |
| 82 | PRJEB13831 | ERR9855084 | sewage | China | single | Metagenome | MetaG | China |
| 315 | UNM_1 | SRR_UNM_1 | ww | Albuquerque, | co | Metagenome | MetaG | USA |
| 326 | PRJNA648659 | SRR13727143 | wwtp influent | Antioquia, Co | co | Metagenome | MetaG | Colombia |
| 326 | PRJNA648659 | SRR12327456 | wwtp influent | Antioquia, Co | co | Metagenome | MetaG | Colombia |
| 326 | PRJNA648659 | SRR12326541 | wwtp influent | Antioquia, Co | co | Metagenome | MetaG | Colombia |
| 326 | PRJNA648659 | SRR12326496 | wwtp influent | Antioquia, Co | co | Metagenome | MetaG | Colombia |
| 326 | PRJNA648659 | SRR12326204 | wwtp influent | Antioquia, Co | co | Metagenome | MetaG | Colombia |
| 326 | PRJNA648659 | SRR12325978 | wwtp influent | Antioquia, Co | co | Metagenome | MetaG | Colombia |
| 327 | PRJNA524094 | SRR8648017 | wwtp influent | Gottingen, G | co | Metagenome | MetaG | Germany |
| 327 | PRJNA524094 | SRR11088433 | wwtp influent | Gottingen, G | co | Metagenome | MetaG | Germany |
| 327 | PRJNA524094 | SRR11088376 | wwtp influent | Gottingen, G | co | Metagenome | MetaG | Germany |
| 327 | PRJNA524094 | SRR11088457 | wwtp influent | Gottingen, G | co | Metagenome | MetaG | Germany |
| 327 | PRJNA524094 | SRR11088438 | wwtp influent | Gottingen, G | co | Metagenome | MetaG | Germany |
| 327 | PRJNA524094 | SRR11088419 | wwtp influent | Gottingen, G | co | Metagenome | MetaG | Germany |
| 327 | PRJNA524094 | SRR11088401 | wwtp influent | Gottingen, G | co | Metagenome | MetaG | Germany |
| 327 | PRJNA524094 | SRR11088478 | wwtp influent | Gottingen, G | co | Metagenome | MetaG | Germany |
| 328 | PRJNA524094 | SRR11088379 | wwtp influent | Greifswald, G | co | Metagenome | MetaG | Germany |
| 328 | PRJNA524094 | SRR11088384 | wwtp influent | Greifswald, G | co | Metagenome | MetaG | Germany |
| 328 | PRJNA524094 | SRR11088366 | wwtp influent | Greifswald, G | co | Metagenome | MetaG | Germany |
| 328 | PRJNA524094 | SRR11088446 | wwtp influent | Greifswald, G | co | Metagenome | MetaG | Germany |
| 328 | PRJNA524094 | SRR11088427 | wwtp influent | Greifswald, G | co | Metagenome | MetaG | Germany |
| 328 | PRJNA524094 | SRR11088408 | wwtp influent | Greifswald, G | co | Metagenome | MetaG | Germany |
| 328 | PRJNA524094 | SRR11088390 | wwtp influent | Greifswald, G | co | Metagenome | MetaG | Germany |
| 328 | PRJNA524094 | SRR11088467 | wwtp influent | Greifswald, G | co | Metagenome | MetaG | Germany |
| 329 | PRJNA524094 | SRR8648011 | hospital effluent | Gottingen, G | co | Metagenome | MetaG | Germany |
| 329 | PRJNA524094 | SRR11088386 | hospital effluent | Gottingen, G | co | Metagenome | MetaG | Germany |
| 329 | PRJNA524094 | SRR11088369 | hospital effluent | Gottingen, G | co | Metagenome | MetaG | Germany |
| 329 | PRJNA524094 | SRR11088449 | hospital effluent | Gottingen, G | co | Metagenome | MetaG | Germany |
| 329 | PRJNA524094 | SRR11088430 | hospital effluent | Gottingen, G | co | Metagenome | MetaG | Germany |
| 329 | PRJNA524094 | SRR11088412 | hospital effluent | Gottingen, G | co | Metagenome | MetaG | Germany |
| 329 | PRJNA524094 | SRR11088393 | hospital effluent | Gottingen, G | co | Metagenome | MetaG | Germany |
| 329 | PRJNA524094 | SRR11088470 | hospital effluent | Gottingen, G | co | Metagenome | MetaG | Germany |
| 330 | PRJNA524094 | SRR11088455 | hospital effluent | Greifswald, G | co | Metagenome | MetaG | Germany |
| 330 | PRJNA524094 | SRR11088459 | hospital effluent | Greifswald, G | co | Metagenome | MetaG | Germany |
| 330 | PRJNA524094 | SRR11088440 | hospital effluent | Greifswald, G | co | Metagenome | MetaG | Germany |
| 330 | PRJNA524094 | SRR11088421 | hospital effluent | Greifswald, G | co | Metagenome | MetaG | Germany |
| 330 | PRJNA524094 | SRR11088403 | hospital effluent | Greifswald, G | co | Metagenome | MetaG | Germany |
| 330 | PRJNA524094 | SRR11088480 | hospital effluent | Greifswald, G | co | Metagenome | MetaG | Germany |
| 330 | PRJNA524094 | SRR11088387 | hospital effluent | Greifswald, G | co | Metagenome | MetaG | Germany |
| 337 | PRJNA532515 | SRR8944124 | wwtp influent | Porto, Portug | co | Metagenome | MetaG | Portugal |
| 337 | PRJNA532515 | SRR8944125 | wwtp influent | Porto, Portug | co | Metagenome | MetaG | Portugal |
| 337 | PRJNA532515 | SRR8944126 | wwtp influent | Porto, Portug | co | Metagenome | MetaG | Portugal |
| 2851 | PRJNA1353891 | SRR36147035 | wwtp influent | Los Alamos, N | single | Metagenome | MetaG | USA |

**Supplementary Table 2:** Count of wastewater sources per environment used in the study

| Wastewater source | Count |
|---|---|
| Sewage | 995 |
| WWTP influent | 122 |
| Hospital | 152 |
| Agricultural | 53 |
| Industrial | 1 |
| Urban | 20 |
| Undefined | 1 |
| **Sum** | **1344** |

**Supplementary Table 3:** Distribution of uSGBs and kSGBs across continent and their pathogenic potential

| Continent | Total SGBs | kSGBs | uSGBs | Pathogenic SGBs | Ratio Unknown | Ratio Pathogens |
|---|---|---|---|---|---|---|
| Global | 26 | 22 | 4 | 4 | 0.15 | 0.15 |
| North America | 636 | 237 | 399 | 28 | 0.63 | 0.04 |
| Euro | 804 | 432 | 372 | 50 | 0.46 | 0.06 |
| Asia | 1404 | 495 | 909 | 77 | 0.65 | 0.05 |
| Africa | 900 | 210 | 690 | 39 | 0.77 | 0.04 |
| South America | 140 | 44 | 96 | 6 | 0.69 | 0.04 |
| Oceania | 97 | 57 | 40 | 8 | 0.41 | 0.08 |
