## Supplementary Tables and Figures for "Global untreated wastewater hosts a vast reservoir of previously uncharacterized microbial lineages": global_ww_supp_figs.pdf

1)

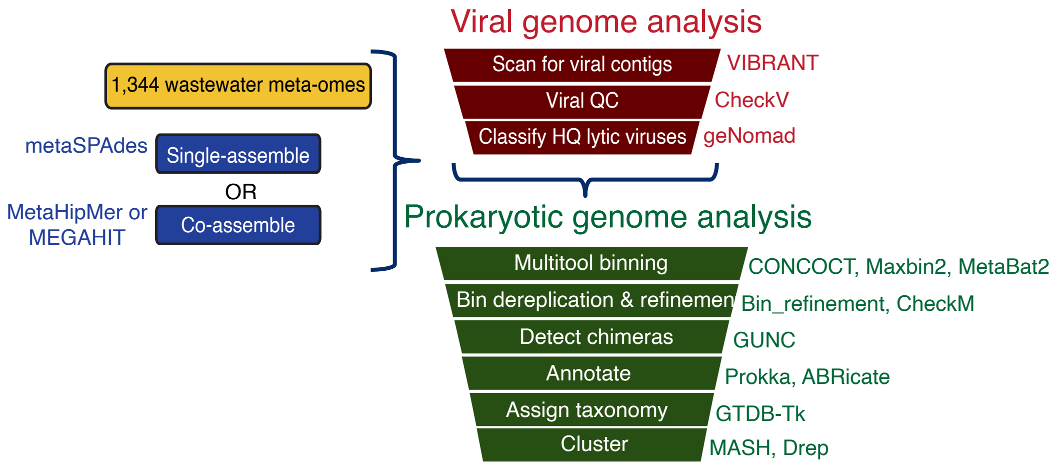


**Supplementary Figure 1**: Bioinformatic pipeline for assembling shotgun metagenomic samples, scanning for viruses, binning, and MAG analysis.

2)

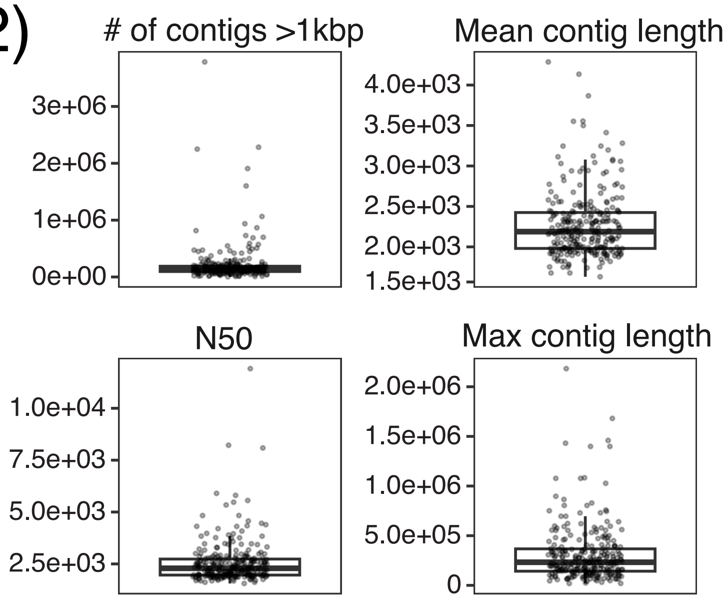


**Supplementary Figure 2:** Statistics of all assembled reads. Each dot represents an assembly.

3)

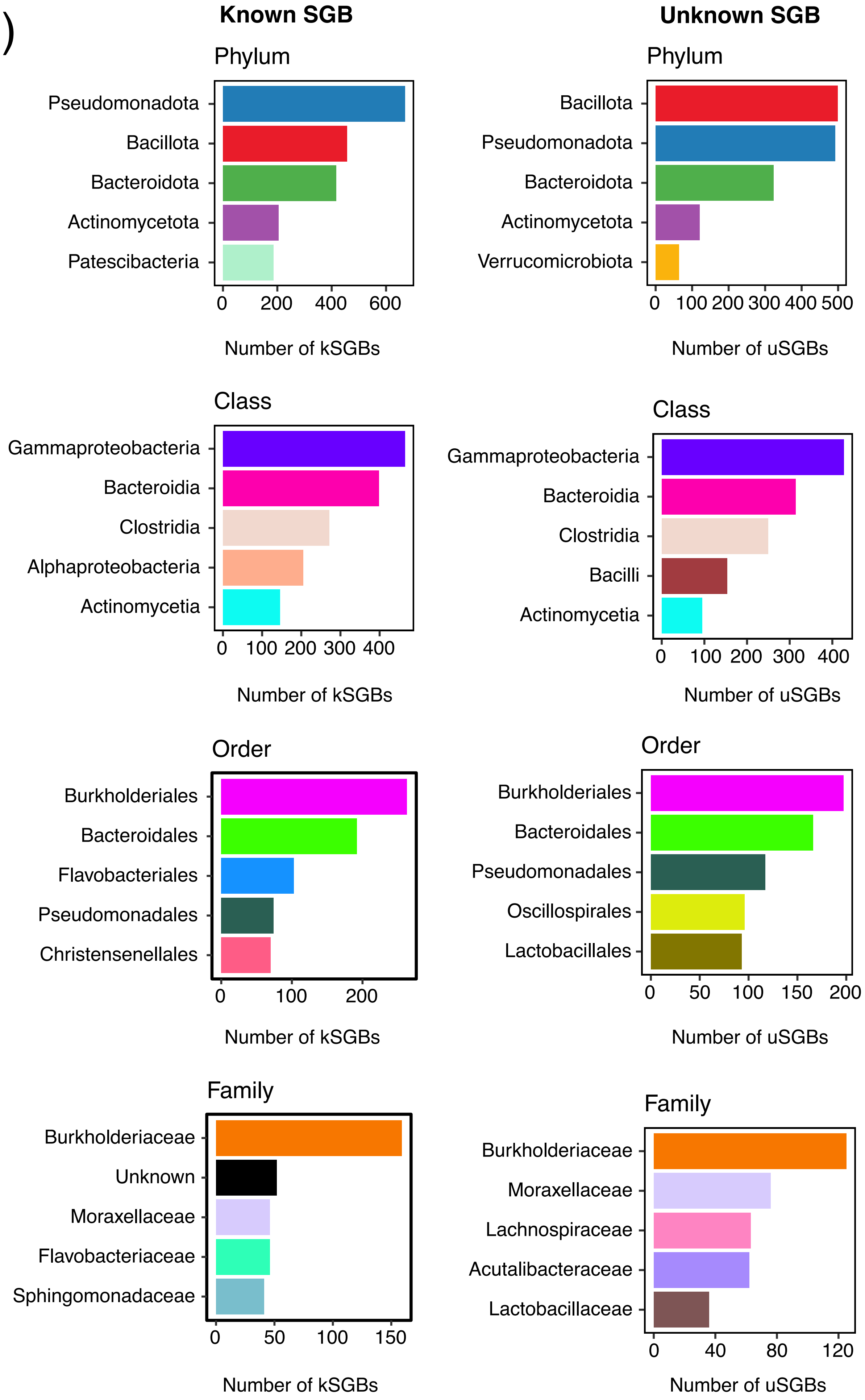


**Supplementary Figure 3:** Top 5 common taxonomic classifications of kSGBs and uSGBs. Genus-level classifications were excluded due to large diversity within each group.

4)

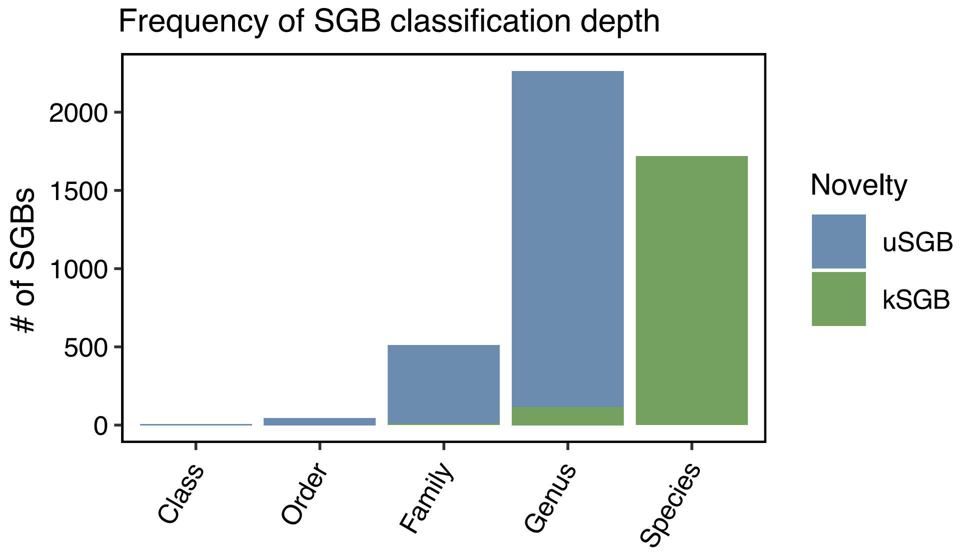


**Supplementary Figure 4:** Frequency of SGB classification depth, colored by known or unknown status. All SGBs clustered to 95% ANI with reference genomes or given a species-level taxonomy by GTDB-Tk were considered known.

5)

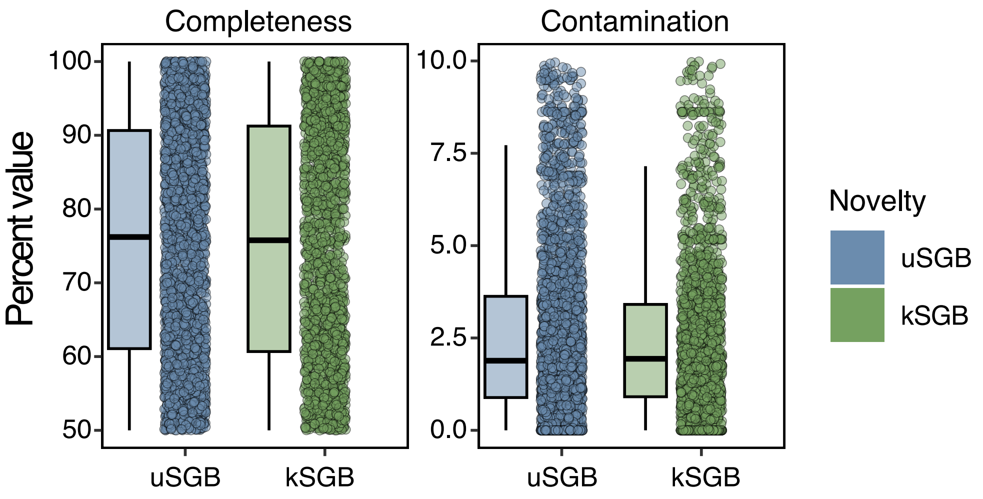


**Supplementary Figure 5:** Completeness and contamination metrics for uSGBs and kSGBs, predicted by CheckM2.

6)

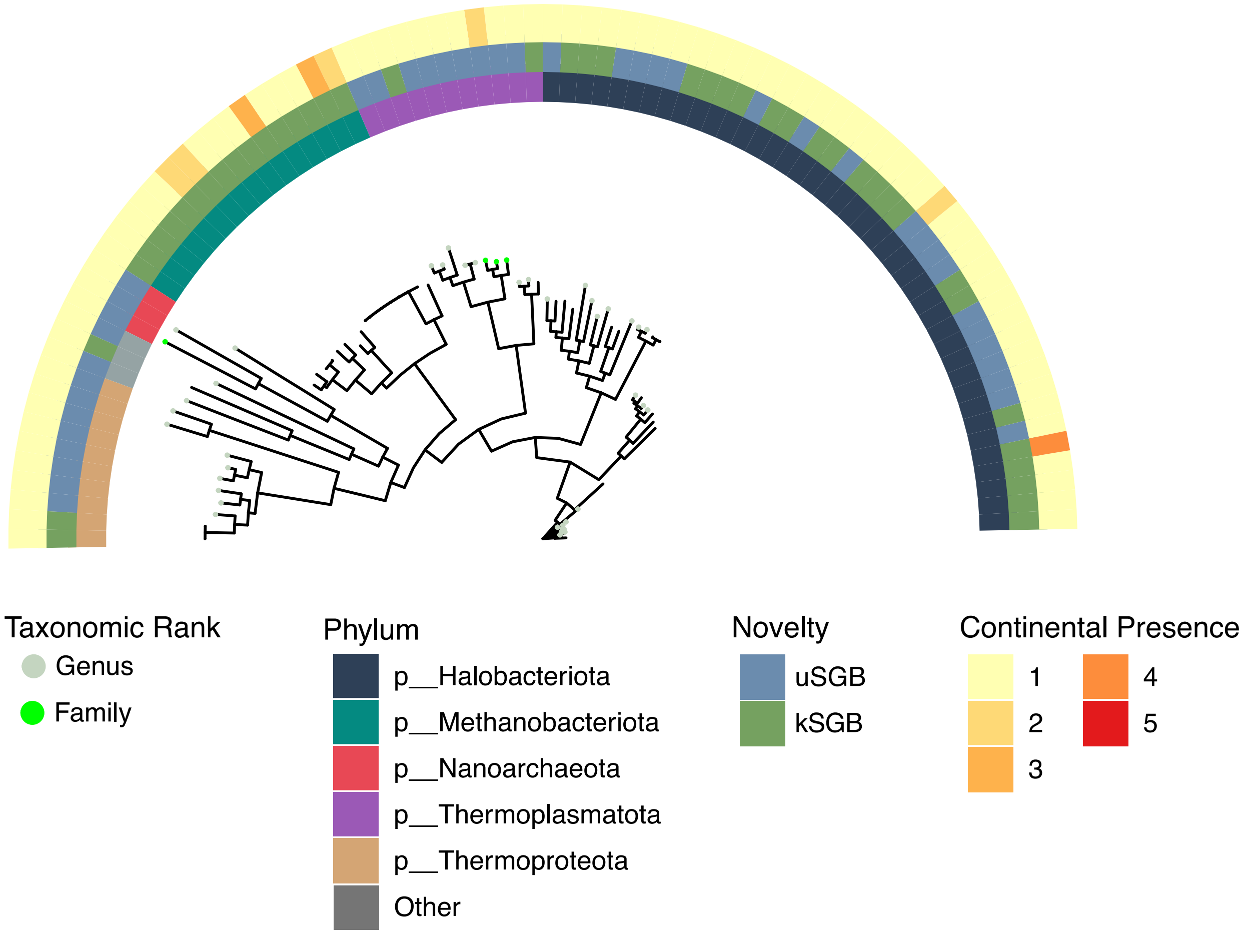


**Supplementary Figure 6:** Archaeal SGB phylogeny. Tip colors represent the assigned taxonomic rank of each SGB excluding species. The inner ring indicates the five most common phyla, the middle represents known or unknown status of the SGB, and the outer ring indicates on how many continents the species was present.

7)

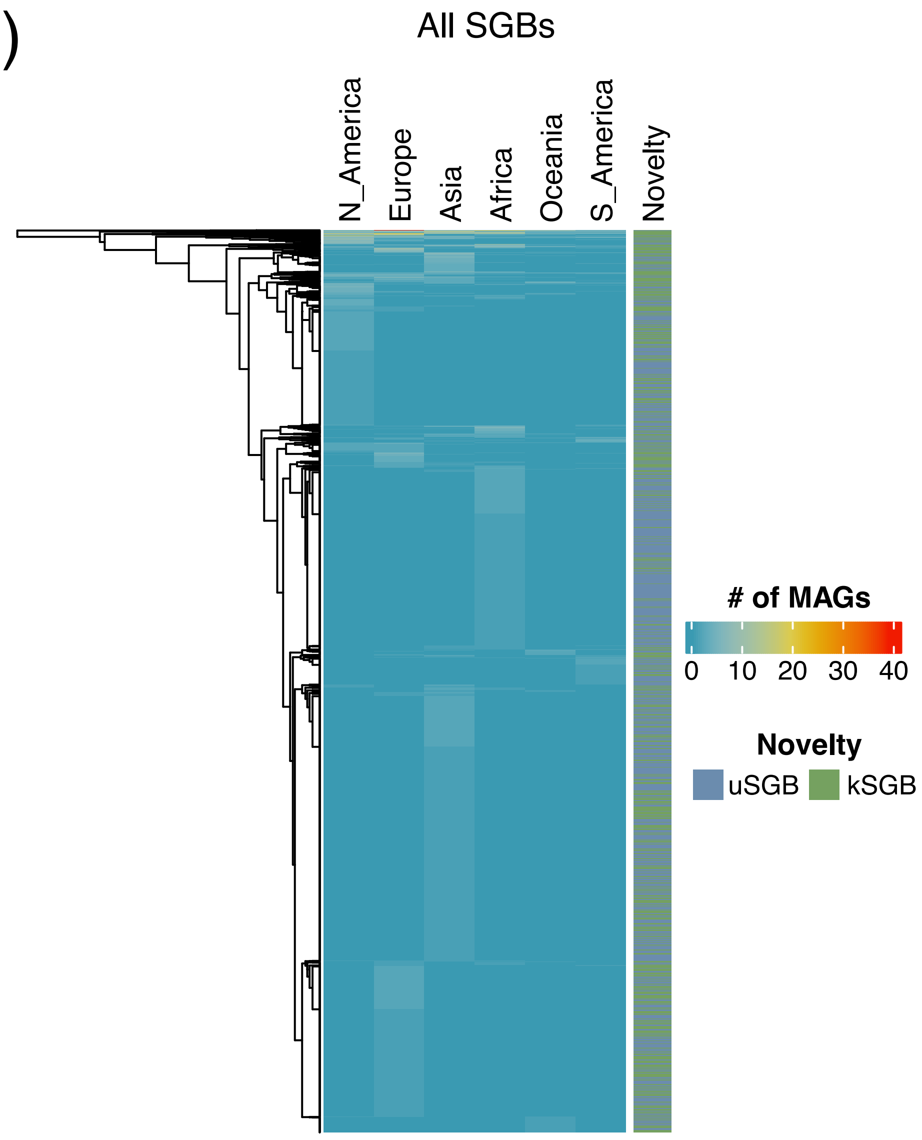


**Supplementary Figure 7:** Heatmap of all SGBs using continuous continental presence.

8)

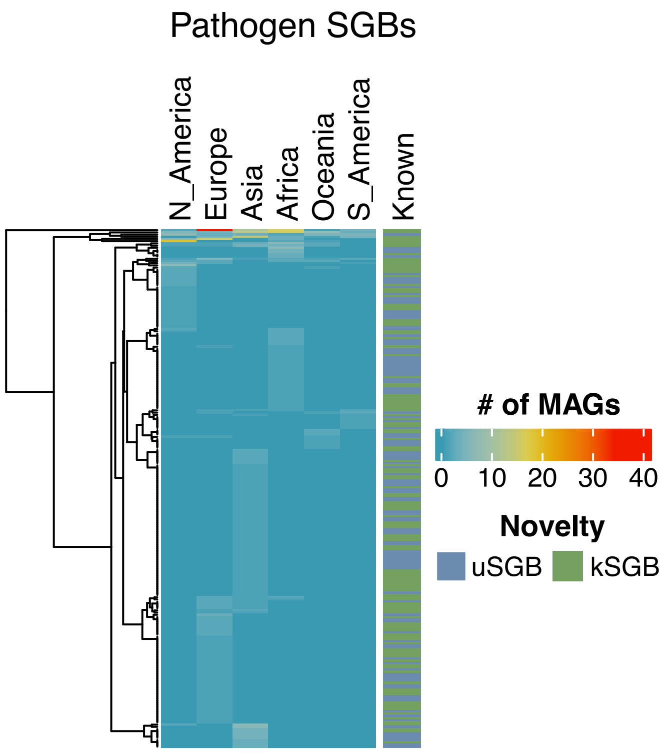


**Supplementary Figure 8:** Heatmap of all pathogenic SGBs using binary continental presence.

9)

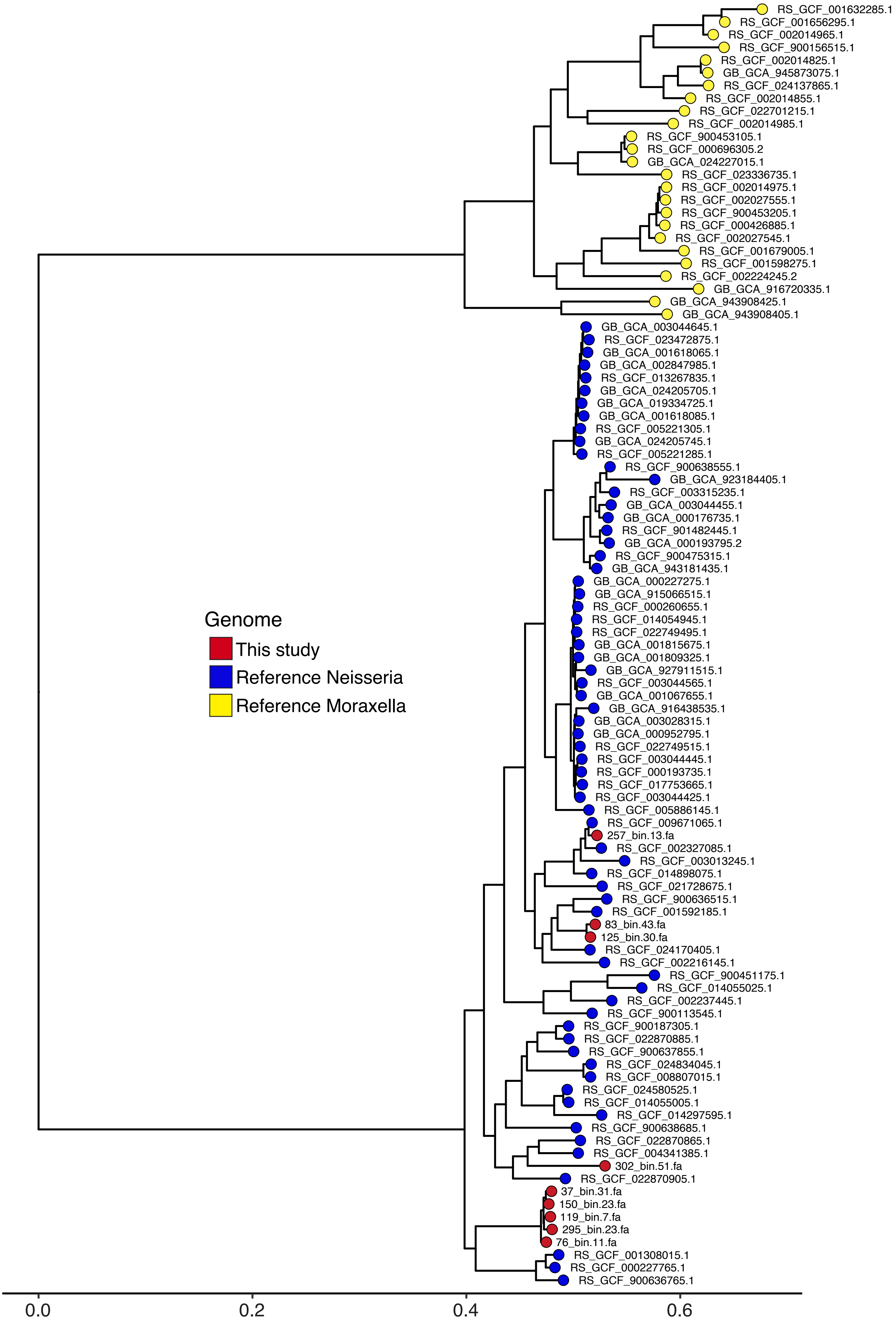


**Supplementary Figure 9:** Phylogeny of unknown Neisseria SGBs and reference Neisseria using Moraxella genomes as the outgroup.
